# Cross-species interactome analysis uncovers a conserved selective autophagy mechanism for protein quality control in plants

**DOI:** 10.1101/2024.09.08.611708

**Authors:** Víctor Sánchez de Medina Hernández, Marintia Mayola Nava García, Marion Clavel, Ranjith K. Papareddy, Veselin I. Andreev, Varsha Mathur, Azadeh Mohseni, Marta García-León, Peng Gao, Juan Carlos de la Concepción, Lorenzo Picchianti, Nenad Grujic, Roksolana Kobylinska, Alibek Abdrakhmanov, Héloïse Duvergé, Gaurav Anand, Nils Leibrock, Anita Bianchi, Margot Raffeiner, Timothy Scott Crawford, Luca Argirò, Mateusz Matuszkiewicz, Cheuk-Ling Wun, Jakob Valdbjørn Kanne, Anton Meinhart, Elisabeth Roitinger, Isabel Bäurle, Byung Ho Kang, Morten Petersen, Suayib Üstün, Yogesh Kulathu, Tim Clausen, Silvia Ramundo, Yasin Dagdas

**Author notes:** Correspondence: Yasin Dagdas. These authors contributed equally to this work (Víctor Sánchez de Medina Hernández, Marintia Mayola Nava García).

## Abstract

Selective autophagy is a fundamental protein quality control pathway that safeguards proteostasis by degrading damaged or surplus cellular components, particularly under stress. This process is orchestrated by selective autophagy receptors (SARs) that direct specific cargo for degradation. While significant strides have been made in understanding the molecular framework of selective autophagy, the diversity of SAR repertoires across species remain largely unexplored. Through a comparative interactome analysis across five model organisms, we identified a suite of conserved and lineage-specific SAR candidates. Among these, we validated CESAR as a conserved SAR critical for proteostasis under heat stress. CESAR specifically facilitates the degradation of hydrophobic, ubiquitinated protein aggregates and is indispensable for heat stress tolerance. Our study offers a rich resource for SAR discovery and positions CESAR as a pivotal regulator of proteostasis, with broad implications for improving stress resilience in plants.

## Introduction

Macroautophagy (hereafter autophagy) is a key quality control mechanism, essential for maintaining organismal homeostasis amidst fluctuating environmental conditions^1^. In both plants and other eukaryotic organisms, autophagy recycles damaged or unnecessary macromolecules and organelles, thereby facilitating cellular renewal and adaptation^2^. Extensive genetic research underscores the vital role of autophagy in enabling organisms to withstand a wide spectrum of abiotic and biotic stressors, including nutrient deprivation, heat stress, and pathogen attacks^3,4^. As global climate change accelerates, leading to more unpredictable weather patterns and heightened pathogen pressures^5^, an improved understanding of plant autophagy mechanisms emerges as a key opportunity to enhance crop resilience and global food security.

Autophagy is characterized by the sequestration of cellular cargo into newly formed double-membrane vesicles known as autophagosomes, which are subsequently directed to lytic compartments for degradation^6^. The formation of autophagosomes is tightly coordinated by a highly conserved set of Autophagy Related Gene (ATG) proteins^6^. Accumulating evidence has demonstrated that autophagic recycling is not merely a bulk degradation process but is executed with remarkable specificity, mediated by selective autophagy receptors (SARs)^7^. SARs act as critical intermediaries, linking autophagic cargo marked with ‘eat-me’ signals, such as polyubiquitin chains, to the autophagy machinery through direct interaction with the core autophagy protein ATG8^8^. SAR-ATG8 interaction is mediated by ATG8-interacting motifs (AIMs)—short linear motifs conserved across eukaryotes, typically defined by the sequence Θ-X-X-Γ, where Θ is an aromatic residue (W/F/Y), Γ is an aliphatic residue (L/V/I), and X represents any amino acid. The aromatic and aliphatic residues crucially anchor to the hydrophobic W and L sites on ATG8, forming the AIM Docking Site (ADS)^10^. *In vitro* reconstitution and cellular biology assays have substantiated a cargo-driven model of autophagosome biogenesis, suggesting that the cargo itself actively participates in its own sequestration by recruiting SARs and other autophagic components^9^. This model suggests the existence of both lineage-specific SARs, adapted to the unique needs of an organism, and more generalist SARs that target broad classes of proteostatic threats, such as those triggered by heat stress. Despite these insights, a comprehensive exploration of SAR diversity across different organisms and how these pathways have evolved in response to distinct evolutionary pressures has yet to be undertaken.

Here, we present a streamlined biochemical approach for identifying novel SARs. Using a peptide competition-coupled affinity purification method, we selectively enriched for direct ATG8 interactors and analysed the ATG8 interactome across five species within the Viridiplantae, generating a valuable resource for SAR discovery. This resource revealed both conserved and lineage-specific interactors, highlighting the flexibility of selective autophagy across different species. Through further characterization, we identified CESAR, a highly conserved SAR in the Streptophyta that plays a critical role in clearing protein aggregates and maintaining protein quality control under proteotoxic stress. Our findings position CESAR as a promising target for enhancing stress tolerance in plants.

## Results

### *Marchantia polymorpha* is a powerful model to investigate selective autophagy

The bryophyte *Marchantia polymorpha* has emerged as a new model plant species due to its low genetic redundancy, the conservation of key molecular pathways found in other land plants, fully sequenced genome, and the availability of genetic tools^10–12^. These features prompted us to establish *M*. *polymorpha* as a model system to study plant selective autophagy. Unlike other model plants such as *Physcomitrium patens* (Pp) and *Arabidopsis thaliana* (At), that encode 6 and 9 ATG8 isoforms, respectively, *M. polymorpha* has only two ATG8 isoforms, MpATG8A and MpATG8B. Phylogenetically, these isoforms represent the two major clades (I and II) of plant ATG8s (Figure 1A, Table S1). To validate *M. polymorpha* as a suitable model system to study selective autophagy, we first performed a GFP release assay^13^ using transgenic *M. polymorpha* Tak-1 (*wild*-type) and *atg7*, an autophagy deficient mutant, expressing GFP-MpATG8A and GFP-MpATG8B. Inhibiting the Tor kinase with Torin treatment successfully led to autophagy induction, with both reporter constructs releasing a free GFP fragment, further stabilized by the inhibition of vacuolar proteases with E64d/pepstatin A treatment^14^. Importantly, the GFP fragment was absent in the *atg7* mutant, confirming both ATG8 isoforms undergo autophagic degradation (Figures 1B, 1C, S1A, S1B).

**Figure 1.**
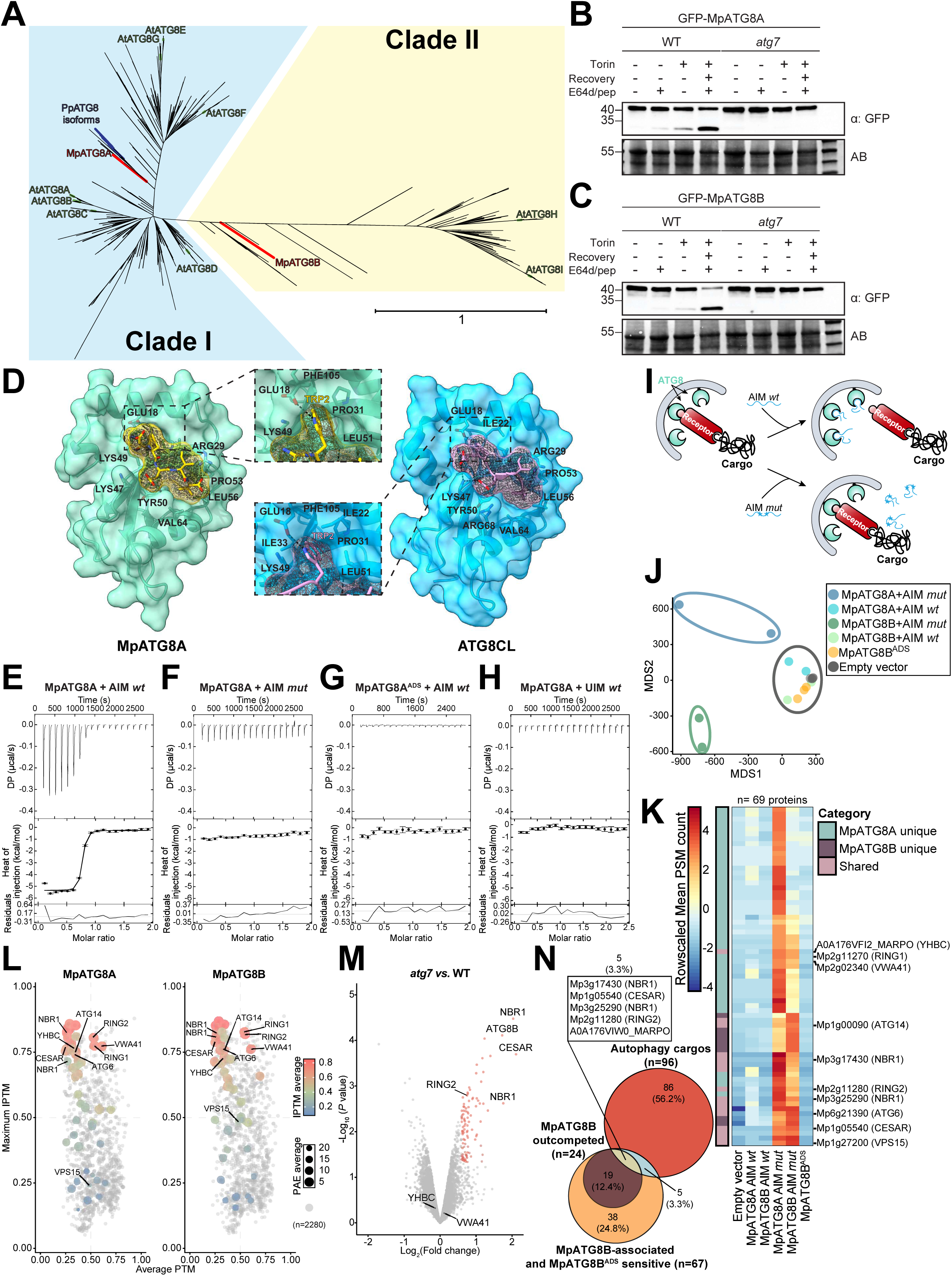
Canonical ATG8 function is conserved in *M. polymorpha*. **(A) *M. polymorpha* (Mp) ATG8 isoforms represent the two major clades of plant ATG8.** Unrooted maximum likelihood phylogram of 621 ATG8 homologs from 80 different plant and algal species with blue and yellow boxes highlighting clades I and II, respectively. *M. polymorpha* (Mp), *P. patens* (Pp) and *A. thaliana* (At) ATG8 isoforms are highlighted in red, blue and green, respectively. **(B-C) MpATG8 isoforms undergo vacuolar degradation in an autophagy-dependent manner.** Autophagic flux assays of transgenic GFP-MpATG8 plants in wild*-type* (Tak-1) or *atg7* mutant background. 5-days old *M. polymorpha* gemmae were incubated in either DMSO or 12 µM Torin-containing medium for 5 hours, after which plants were either harvested or recovered for additional 16 hours with fresh media containing 10 µM E64d and 10 µM pepstatin A (E64d/pep) before harvesting. Protein extracts were analysed by immunoblotting with anti-GFP antibody. Total protein loading control was analysed by Amidoblack (AB) staining. A second independent replicate for each isoform is shown in Fig. S1A,B. **(D) MpATG8 binds AIM in a canonical manner.** Schematic representation of the X-ray crystal structure of MpATG8A bound to the canonical AIM *wt* peptide (left) and its comparison to potato ATG8CL structure (PDB: 5L83) bound to the same peptide (right). MpATG8A and ATG8CL are shown as a surface representation in green and blue, respectively, and the peptide is shown as sticks with yellow (left) or pink (right) carbon atoms. The electron density omit map of the peptide ligand is shown as mesh and contoured at 2 σ. Electrostatic interactions are indicated with black dashed lines. Residues involved in the interaction are highlighted. Tryptophanś side chains docking on the hydrophobic pockets on ATG8 are shown in insets. (E-F) MpATG8A binds AIM *wt* but not AIM *mut*. Isothermal titration calorimetry (ITC) experiments showing titrations of AIM *wt* (E) or AIM *mut* (**F**) peptides to MpATG8A. The concentration of reactants is 50 µM for MpATG8A (in cell) and 500 µM peptide (in syringe). Three independent replicates for AIM *wt* and AIM *mut* are shown in Fig. S1C and Fig. S2A, respectively. (G) MpATG8A^ADS^ does not bind AIM *wt*. ITC experiments showing titrations of AIM *wt* peptide to MpATG8A^ADS^. The concentration of reactants is 25 µM for MpATG8A (in cell) and 250 µM peptide (in syringe). Three independent replicates are shown in Fig. S2B. (H) MpATG8A does not bind UIM *wt*. ITC experiments showing titrations of UIM *wt* (D) peptide to MpATG8A. The concentration of reactants is 50 µM for MpATG8A (in cell) and 500 µM peptide (in syringe). Three independent replicates are shown in Fig. S2D. Global analysis was performed using a hetero-association model A + B. The top panels show the SVD-reconstructed thermograms, the middle panels show the isotherms as integrated heat of injection (dots) and best fit (solid line), and the bottom panels show the residuals (difference between the fit to a single hetero-association model and the experimental data. Extracted global parameters and their 68.3% confidence interval are reported in Fig. S1C,D. **(I) Cartoon representing the peptide competition approach.** Addition of AIM *wt* would outcompete direct ATG8 interactors, whereas addition of AIM *mut* would have no effect. **(J) Peptide competition depletes ATG8 interactors and mimics the ADS mutation.** Multidimensional scaling (MDS) plot of MpATG8 interactome highlighting the global variation between experiments. **(K) Peptide competition enriches for direct ATG8 interactors.** Protein abundance pattern represented by a heatmap (Log2(Mean PSM+1)-Mean PSM per protein) for the 69 associated and outcompeted proteins (see Fig. S3E,F). Proteins are annotated based on their enrichment in one (MpATG8A/B unique) or multiple (shared) samples. Each column represents the row-scaled mean PSM count. **(L) AF2 multimer-based interactome analysis partially captures peptide outcompeted candidates.** Dot plot representing AF2 multimer interaction predictions of whole interactome dataset including the proteins associating with empty vector control against each MpATG8 isoform. The 69 associated and outcompeted proteins showed in Fig. 1K are highlighted based on their average **P**redicted **A**ligned **E**rror (PAE) and **I**nterface **P**redicted **T**emplate **M**odelling (IPTM) values. (M) Quantitative proteomics of *M. polymorpha* reveals autophagy-dependent degradome. Protein enrichment in *atg7* plants compared to wild*-type* (Tak-1) plants represented by a volcano plot. Protein abundance was measured in untreated 10-day-old *M. polymorpha* plants using 16-plex TMT labelling. Enriched proteins are highlighted in red. **(N) Selection of bona-fide autophagy receptor candidates.** Venn diagram of three overlapping pairwise comparisons. Autophagy cargos (red circle) are defined as in Fig. 1M. MpATG8B-associated and MpATG8B^ADS^ sensitive proteins (orange circle) are defined as in Fig. S3G. MpATG8B outcompeted proteins (orange circle) are defined as AIM-dependent proteins in Fig. S3C.

Next, we tested whether MpATG8 interacts with the ATG8-interacting motif (AIM). We determined the X-ray crystal structure of MpATG8a in complex with a canonical AIM peptide (AIM *wt*)^15,16^ at 1.75 Å resolution (Figure 1D, Table 1). The structure revealed that MpATG8A binds the AIM peptide via the AIM-docking site (ADS), similar to yeast, human, and plant ATG8s^7,17^. Next, we assessed binding of MpATG8A and MpATG8B to AIM *wt* using isothermal titration calorimetry (ITC). Both isoforms interacted with AIM *wt*, with a KD∼60 nM and 750 nM, respectively (Figures 1E, S1C and S1D). These affinities are within similar range as the *A. thaliana* AtATG8A and human GABARAP ITC measurements that we previously assessed for the same peptide^16,18^. Mutating the Θ and Γ positions to Alanines in the AIM peptide (AIM *mut*) abrogated binding to MpATG8 (Figures 1F, S2A) and consistently, MpATG8 AIM Docking Site (ADS) mutants failed to bind AIM *wt* peptide (Figures 1G, S2B). Finally, we tested the *in vivo* association of MpATG8 isoforms with the AIM containing SAR MpNBR1. We showed that MpNBR1 co-immunoprecipitated with both MpATG8 isoforms, while competition with the AIM *wt* peptide or mutating the ADS of MpATG8 isoforms prevented NBR1-MpATG8 interaction (Figure S2C). Overall, these experiments demonstrate that the ATG8 AIM-ADS interface is conserved in *M. polymorpha*.

**Table 1:**
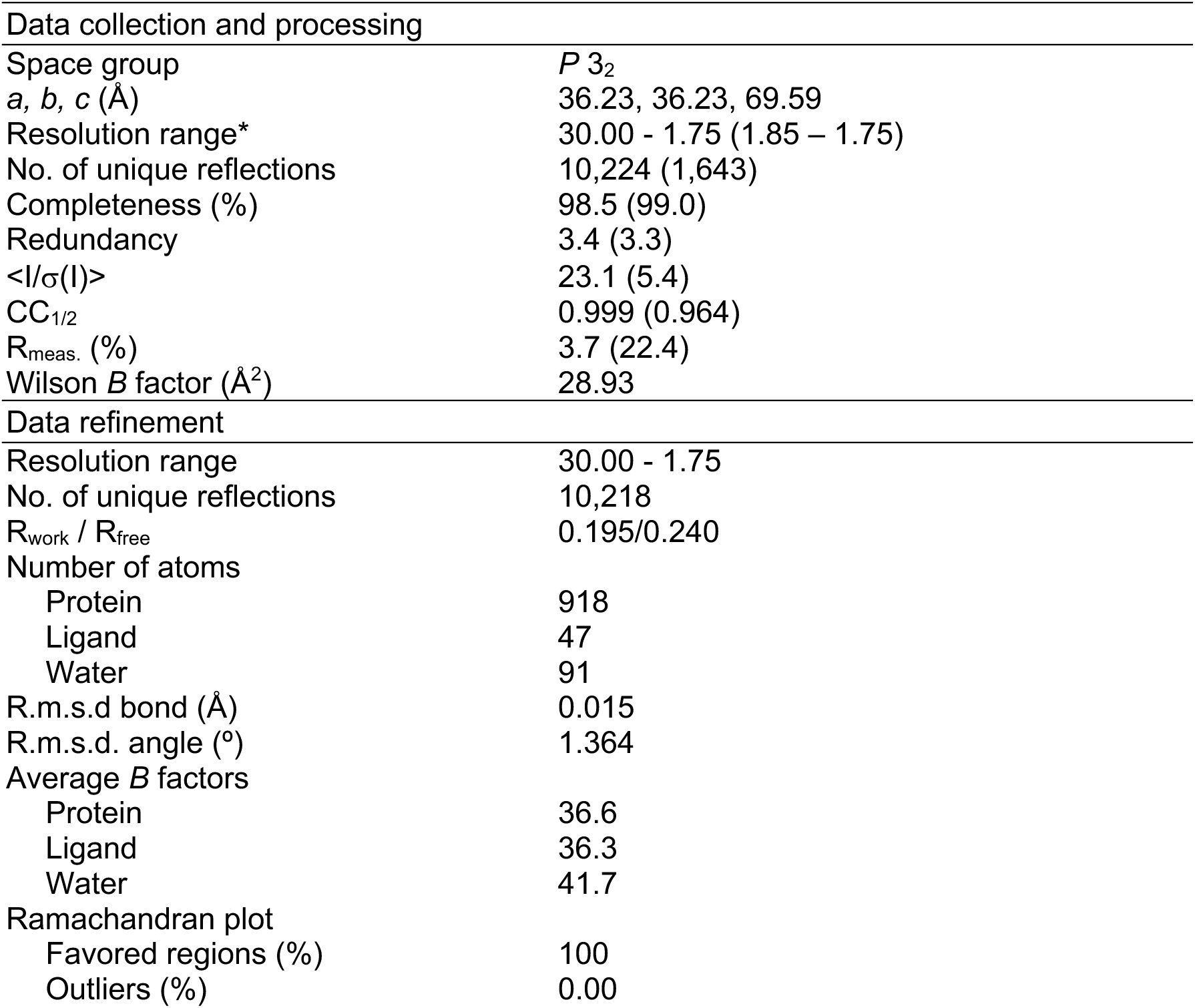
Diffraction data collection and model refinement.

Recent work suggests a category of SARs contain an additional binding motif – the ubiquitin-interacting motif (UIM) - and bind to the UIM-docking site (UDS) within ATG8, located opposite to the ADS^7,17^. To test if MpATG8 could bind to a UIM peptide, we performed ITC experiments using the AtRPN10 UIM peptide (UIM *wt*). We detected no binding between MpATG8 and UIM *wt* (Figures 1H, S2D). To further probe ATG8-UIM interaction, we purified the C-terminal portion of AtRPN10 (AtRPN10^201–386^) and assessed its binding to AtATG8A via ITC. Unlike previously reported^17^, AtRPN10^201-386^ did not bind AtATG8A (Figure S2E). Since we could not recapitulate ATG8-UIM interaction and UDS residues have been recently shown to interact with the autophagosome membranes^19^, we decided to focus only on AIM-dependent ATG8 interactions.

### Peptide competition-coupled affinity purification fast-tracks the identification of AIM-dependent ATG8 interactors

We sought to use our *M*. *polymorpha* system to identify novel candidate SARs using affinity purification coupled with mass spectrometry (AP-MS). Since all SARs identified to date contain AIMs, we hypothesized that adding an AIM peptide competition step to ATG8 affinity purification could selectively deplete AIM-dependent ATG8 interactors and facilitate SAR discovery. To test our hypothesis, during the affinity purification, we incubated the *M. polymorpha* lysates with an excess of either AIM *wt* or AIM *mut* peptide and analysed both fractions with mass spectrometry (Figure 1I). We identified a total of 272 MpATG8-associated proteins, 93 of them shared between MpATG8 isoforms (Figures S3A-D). Consistent with our hypothesis, multidimensional scaling (MDS) analysis revealed that peptide competition reduced the complexity of the interactome (Figure 1J). Strikingly, 61 out of 244 MpATG8A associated proteins (Figure S3E) and 24 out of 121 for MpATG8B associated proteins (Figure S3F) were outcompeted by the AIM *wt* peptide, leading to a 4-fold and 5-fold reduction of the MpATG8A and MpATG8B interactomes, respectively (Figure 1K, Table S2). Consistent with our previous experiment (Figure S2C), MpNBR1 was amongst the outcompeted proteins, as well as VPS15, ATG14, ATG1, and ATG6^20^, all core autophagy proteins necessary for the completion of autophagy.

To further test the peptide competition approach, we compared the outcompeted MpATG8B interactome with the MpATG8B^ADS^ interactome, as ADS mutation should phenocopy the interaction depletion caused by the AIM *wt* peptide (Figures 1G, S2B, S2C). We identified 67 MpATG8B-associated proteins that were depleted in MpATG8B^ADS^ samples (Figure S3G). All 24 peptide-outcompeted proteins were sensitive to the ADS mutation (Figure S3H, Table S2), suggesting that the peptide competition approach reduces the complexity of ATG8 interactomes and enriches for direct ATG8 interactors.

Recently, Alphafold2 (AF2) has emerged as a powerful *in silico* approach for predicting protein-protein interactions^21,22^. To test if AIM-dependent candidate detection could be further refined by *in-silico* AF2 predictions, we calculated the interaction scores of the 2280 proteins identified in our interactome against MpATG8 isoforms. Out of the identified 69 non-redundant outcompeted candidates, AF2 predicted 48 proteins (∼70%) as having well-supported interaction interfaces (iPTM>0.5) (Figure 1L, Table S3), allowing us to further define high confidence interactors.

Even though we identified known AIM containing proteins such as MpNBR1 in our datasets, to test if the peptide competition approach could facilitate the identification of novel AIM containing proteins, we selected 5 uncharacterized proteins: MpYHBC, a ribosome maturation factor (RimP) domain-containing protein; MpRING1/2, two RING-finger domain-containing paralogous proteins; and MpVWA41/43, two vault inter-trypsin (VIT) and von Willebrand factor (VWA) domain-containing paralogous proteins (Figure S4A, S5A, S5D, S6A, S6D). We first tested their ability to bind MpATG8 using *in vitro* pulldown assays. All tested candidates bound MpATG8 in an AIM-dependent manner, as their binding was fully abrogated by the addition of the AIM *wt* peptide, and they did not bind MpATG8^ADS^ (Figure S4B, S5B, S5E, S6B and S6E). We identified the relevant AIMs within the candidates using mutagenesis. This approach revealed a single AIM within MpYHBC (Figure S4C), an AIM shared by both MpVWA paralogues (Figure S6C, S6F) and two synergistic AIMs within both MpRING proteins (Figure S5C, S5F). Interestingly, phylogenetic analyses revealed different evolutionary histories for each candidate (Figures S4D, S5G, S6G). Unlike YHBC, in which the RimP domain originated in a prokaryotic ancestor, both VWA and RING paralogs have a eukaryotic origin. YHBC and RING proteins are conserved across the Viridiplantae, whereas VWA are bryophyte-specific and not found in the Tracheophyta (vascular plants). Furthermore, although the AIM residues are conserved across the Viridiplantae for RING proteins, only the bryophyte orthologs of YHBC and VWA have a conserved AIM (Figures S4D, S5G, S6G). These findings suggest peptide-competition enriches for AIM-dependent ATG8 interactors and underscore the dynamic evolutionary history of ATG8 interacting proteins.

Since another canonical feature of SARs is undergoing autophagic degradation together with their cargo^23^, we hypothesized that SAR protein levels would be higher in *atg7* mutants when compared to *wild*-type plants. To test our hypothesis, we performed tandem mass tag (TMT)-based quantitative proteomics and compared the proteomes of *M. polymorpha* Tak-1 and *atg7* mutant plants under control conditions. Consistent with our hypothesis, MpNBR1 proteins were among the most significantly enriched proteins in *atg7* with a 4-fold enrichment, together with MpATG8B and a previously uncharacterised protein that contains a CUE (**C**oupling of **U**biquitin to **E**R degradation) and ELKS (protein rich in the amino acids **E**, **L**, **K** and **S**) domain that we named as CESAR (CUE and ELKS domain Containing SAR) (Figures 1K, 1M). Among the significantly enriched proteins, we also found the identified outcompeted protein MpRING2 (Figure 1N), indicating they are degraded via autophagy. Other identified outcompeted proteins, such as MpYHBC or MpVWA41, were not enriched in *atg7* under these conditions (Figure 1M), suggesting they may only serve as SARs under certain conditions. Altogether, we showed that peptide competition streamlines the discovery of potential selective autophagy players and paves the way for the systematic analysis of ATG8 interactomes under different stress conditions or in different model organisms.

### Cross-Species Analysis of ATG8 Interactomes Reveals novel SAR candidates

To explore the diversity of selective autophagy networks, we conducted a cross-species analysis of ATG8 interactomes (Figure 2A). We generated lines that express fluorescent protein fusions of ATG8 and performed peptide competition coupled AP-MS in the green algae *Chlamydomonas reinhardtii*, the moss *Physcomitrium patens*, and two angiosperms, *Arabidopsis thaliana* and *Nicotiana benthamiana*, which together span approximately one billion years of evolution (See Materials and Methods for details) (Figure 2A, Tables S4, S5, S6, S7). This analysis identified 289 (*C. reinhardtii*), 222 (*P. patens*), 339 (*A. thaliana*), and 56 (*N. benthamiana*) ATG8-associated proteins, respectively. Consistent with our findings in *M. polymorpha*, peptide competition reduced the ATG8 interactomes by approximately 20-fold in *C. reinhardtii* (17 out of 289), 4-fold in *P. patens* (59 out of 222), and by about 2-fold in both *A. thaliana* (192 out of 339) and *N. benthamiana* (27 out of 56) (Tables S4, S5, S6, S7).

**Figure 2.**
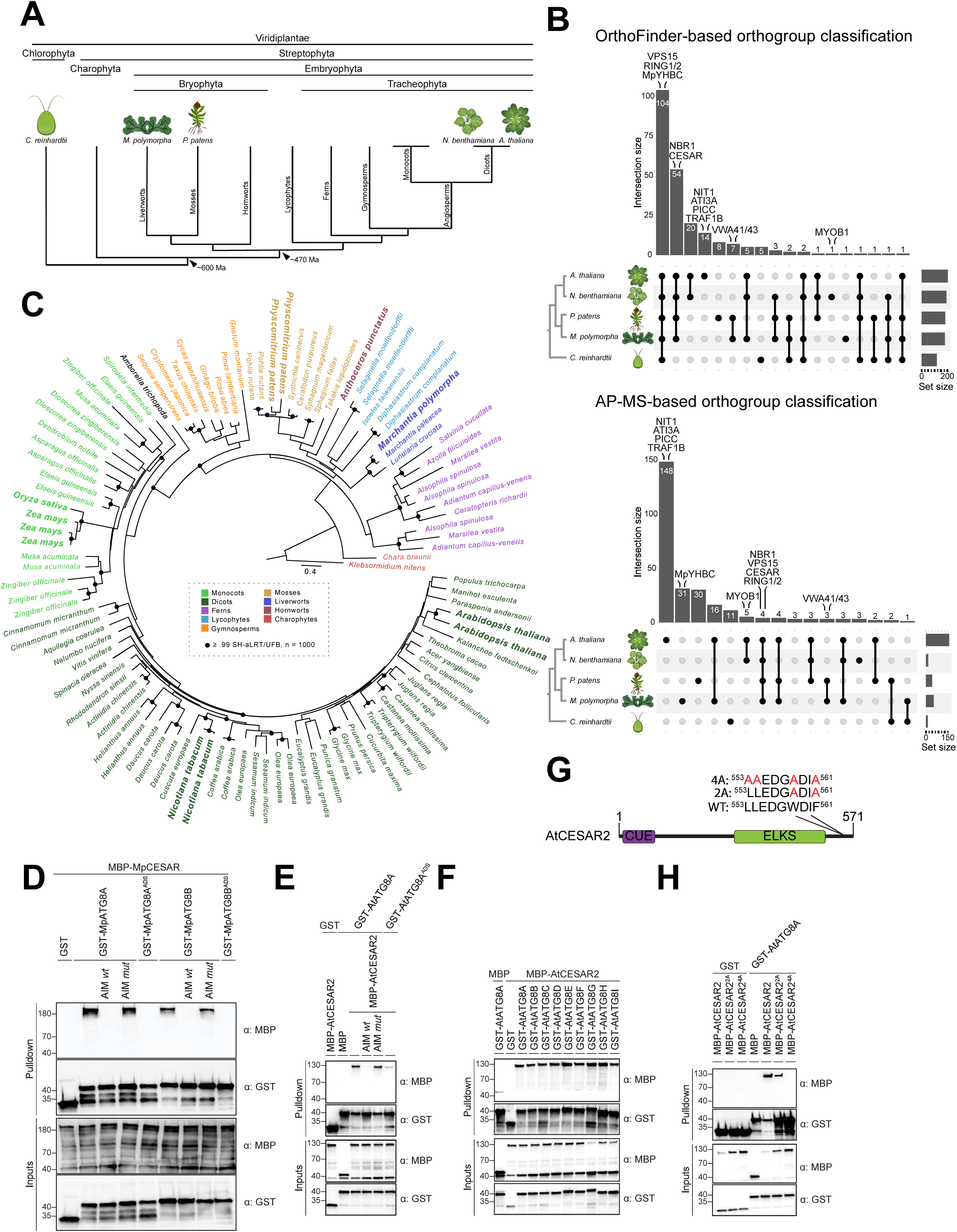
Peptide competition-coupled comparative ATG8 interactome analyses identifies CESAR as an evolutionary conserved ATG8 interactor. **(A) Phylogenetic relationships between species used for ATG8 interactome analyses.** Schematic phylogram of the Viridiplantae. Species used for the comparative AP-MS interactome analyses are represented. **(B) Orthologous cluster analysis of outcompeted proteins highlights the dynamism of autophagy pathways.** UpSet plots displaying unique and shared orthologous clusters determined by OrthoFinder among the depicted species represented in the left cladogram phylogeny. The intersection size is shown by the vertical bar chart, and the set size is shown by the right horizontal bar chart. Ortholog groups are ordered by the intersection size, and overlaps are represented by vertical lines. Orthogroups are classified either based on the identification of orthologous sequences from any given outcompeted protein (top panel) or strictly limited to experimentally identified orthologous proteins in the tested species (bottom panel). Proteins of interest are highlighted above their clusters. **(C) CESAR evolved in Streptophyta.** Maximum likelihood phylogenetic tree of 107 non-redundant CESAR homologs from 72 different plant and algal species constructed using the LG4M+F+R6 substitution model in IQ-TREE. Charophytes are used for outgroup-based tree-rooting. Branches are colored for sequences belonging to the indicated clade/family, except *Amborella trichopoda* (black). CESAR orthologs from the species used for biochemical assays in Fig. 2 and Fig. S7 are highlighted in bold. Shimodaira-Hasewaga with approximate likelihood-ratio test (SH-aLRT) and ultrafast bootstrap support (UFB) values (n = 1,000) greater than 99% are shown on the branches as black circles. (D-E) MpCESAR and AtCESAR2 bind ATG8 in an AIM-dependent manner *in vitro* (MpATG8A^ADS^=MpATG8A^(Y52A,L53A)^, MpATG8B^ADS^=MpATG8B^(K51A)^, AtATG8A^ADS^=AtATG8A^(Y50A,L51A)^. AIM *wt* and AIM *mut* peptides were added to a final concentration of 200 µM. (F) AtCESAR2 binds all AtATG8 isoforms *in vitro*. (G) AtCESAR2 has an AIM in its C-terminal. Protein domain architecture of AtCESAR2. The CUE and ELKS domains are highlighted in purple and green, respectively. AIM residues, their positions and mutagenesis are shown. **(H) AIM mutagenesis of AtCESAR2 abrogates AtATG8A binding** AtCESAR2^2A^=AtCESAR2^(W558A,F561A)^, AtCESAR2^4A^=AtCESAR2^(L553A,L554A,W558A,F561A)^. Bacterial lysates containing recombinant protein were mixed and pulled down with glutathione magnetic agarose beads. Input and bound proteins were visualized by immunoblotting with anti-GST and anti-MBP antibodies.

To further understand the evolutionary history of these outcompeted proteins, we identified their orthologues across all five species using Orthofinder^24^ and visualized their intersections (Figure 2B, Table S8). Although most of the outcompeted proteins have orthologous sequences within all five species, we observed cases where proteins were detected in only a single species, likely reflecting lineage-specific gains or losses of these proteins. This is exemplified by the MpVWA-domain containing proteins that based on our phylogenetic analyses have bryophyte-specific origins (Figure S6G). Interestingly, only 12% of the candidate proteins were classified in species-specific orthologous groups, with *A. thaliana* and *P. patens* having the largest number of observations (Figure 2B, upper panel). This suggests that most of these gene families were present in the last common ancestor of the Viridiplantae and were co-opted for autophagic function in a lineage-specific manner.

To test this idea further, we checked whether our experimentally recovered AIM-dependent ATG8 interactors matched their orthology-based groupings. We repeated the analysis, this time restricting the classification of orthogroups to those based on experimental validation, meaning that only orthologous proteins that were simultaneously identified in different species were considered as belonging to the same orthogroup (Figure 2B, lower panel, Table S9). We could recapitulate the bryophyte-specific origin of MpVWA, for which we mapped its AIM (Figure S6). However, some of the highly conserved orthologous sequences were experimentally identified as being species-specific, such as MpYHBC. This may indicate that the simple acquisition of an AIM can confer a new role in autophagy to a pre-existing protein. Notably, we found that NBR1, VPS15, CESAR and MpRING1/2 were identified across all streptophytes suggesting their long-lasting evolutionary conservation in both orthology and function as SARs and/or core autophagy proteins.

In summary, our cross-species ATG8 interactome analysis uncovers novel AIM-dependent ATG8 interactors and highlights the evolutionary diversity of ATG8 interactome.

### CESAR localizes to autophagosomes and undergoes autophagic degradation

Next, we decided to functionally characterize one of the candidate SAR proteins that so far remained as an uncharacterized protein in streptophytes. We selected CESAR for further investigation because **(i)** it interacts with ATG8 and is outcompeted by the AIM *wt* peptide in all four land plant species tested, and **(ii)** in our quantitative proteomics assays, it accumulates higher in *atg7* mutants compared to *wild*-type *M. polymorpha* plants and **(iii)** it is identified as a high-confidence interactor in our AF2 analysis (Figures 1L, 1M, 1N, 2B). Phylogenetic analyses revealed that CESAR first evolved in Charophyta algae and subsequently underwent gene duplication events in mosses, followed by additional duplication and gene loss throughout plant evolution (Figure 2C). To test the conservation of CESAR-ATG8 interaction across plants, we expressed CESAR orthologs from seven different plant species spanning various branches of plant phylogeny and demonstrated that all tested CESAR orthologs bind ATG8 in an AIM-dependent manner (Figures 2D-F, S7). Further AIM-mapping within the *A. thaliana* CESAR2 ortholog confirmed the presence of an AIM in the C-terminal region of the protein (Figures 2G, 2H).

Next, we sought to functionally characterize the CESAR-mediated autophagy pathway. Since CESAR shared enrichment patterns with the aggrephagy receptor NBR1 in all our assays, we hypothesized that CESAR could be involved in response to proteotoxic stress. To test this hypothesis, we took advantage of the well-established proteotoxic stress assays in *Arabidopsis thaliana* and performed two further ATG8 interactome analyses using two different proteotoxic stress treatments: **(i)** heat stress (HS) and **(ii)** heat stress supplemented with CB5083 (CB) treatments. CB inhibits CDC48, an essential AAA+ ATPase that presents ubiquitinated proteins to the proteasome for degradation^25,26^. We hypothesized that blocking CDC48 would channel ubiquitinated proteins to autophagy. Peptide competition coupled AP-MS analyses with two different AtATG8 isoforms (AtATG8A and AtATG8E) revealed 33 AIM-dependent interactors that responded to proteotoxic stress (Figures 3A, S8, S9A, Table S10). We observed that AtCESAR2 was amongst the top candidate proteins enriched in ATG8 IP upon proteotoxic stress, while simultaneously being outcompeted, a feature that was shared with NBR1 (Figures 3B, S9B-E). Additionally, we performed proximity labelling proteomics with TurboID (TID) tagged AtATG8E, AtATG8E^ADS^ and AtATG8E^G118A^ (a lipidation deficient mutant that cannot be conjugated to the membrane of autophagosomes^27^). We chose AtATG8E as a representative of Clade I ATG8s, which contains MpATG8A and AtATG8A (Figure 1A). We reasoned that the comparison of ATG8 proxitomes in our treatment conditions with that of the G118A mutant would reveal proteins that are uniquely present in stress-induced autophagosomes. These experiments revealed additional ATG8 proximal proteins upon proteotoxic stress that were not captured with AP-MS (Figures 3C, S10, Table S11). Overall, 34 proteins are shared between the AP-MS and the TurboID experiments, including AtCESAR1 and AtCESAR2 (Figure 3D, Table S11). Altogether, these results are consistent with our hypothesis that CESAR-ATG8 interaction is induced by proteotoxic stress and CESAR may localize to autophagosomes (Figure 3C).

**Figure 3.**
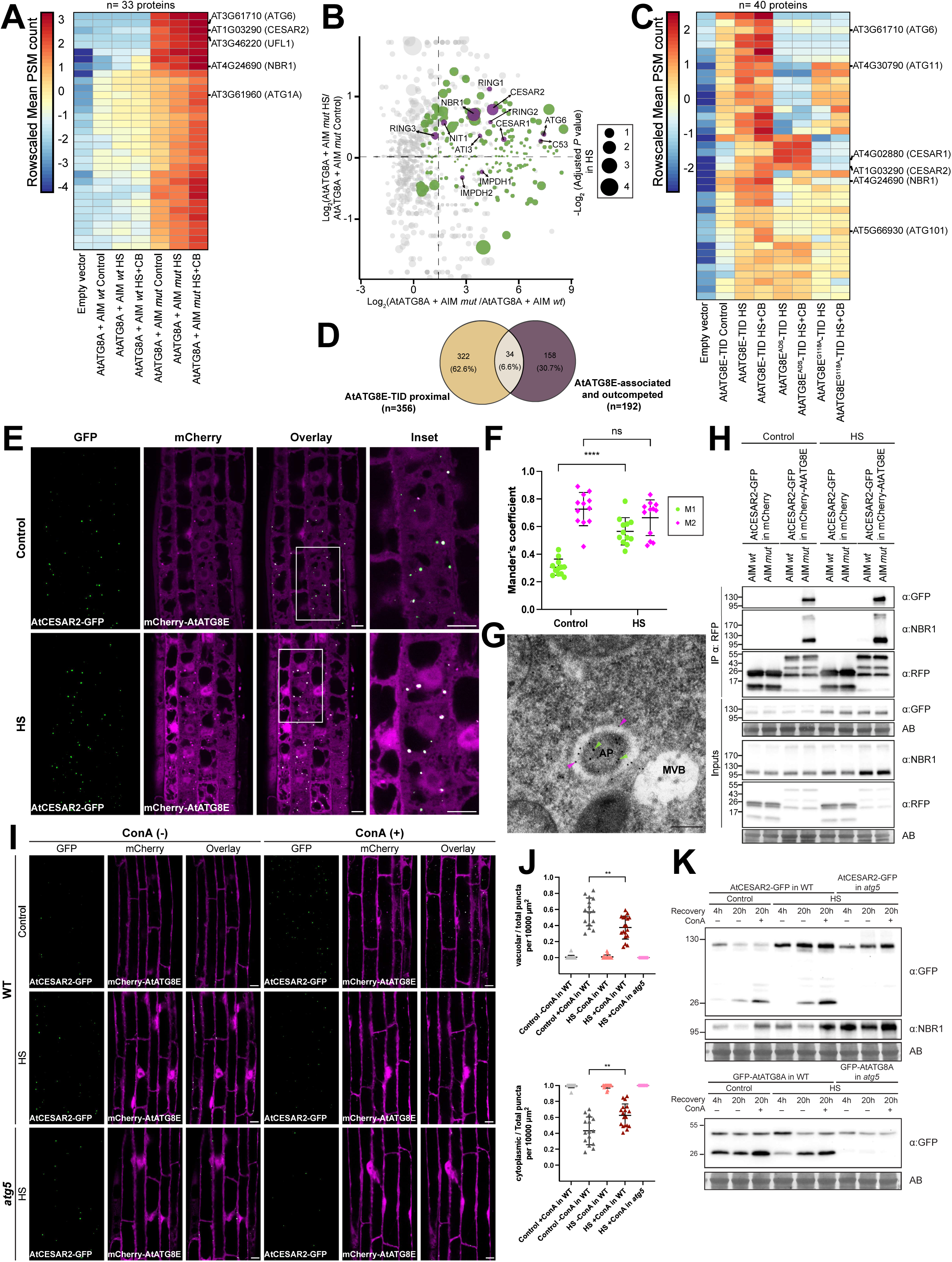
CESAR localizes to autophagosomes and undergoes autophagic degradation. **(A) AtATG8A interactome upon HS+CB treatment.** Protein abundance pattern represented by a heatmap (Log2 (Mean PSM+1) – Mean PSM per protein) for the 33 AtATG8A associated and outcompeted proteins identified as induced upon HS+CB. Each column represents the row-scaled mean PSM count of 3 independent biological replicates. **(B) AtATG8A interactome upon HS treatment.** Dot plot representing protein abundance as Log2 (Fold change) for the ratio of two pairwise comparisons comparing the peptide effect (AtATG8A+AIM *mut* vs. AtATG8A+AIM *wt*) on the x-axis and the treatment effect (AtATG8A+AIM *mut* HS vs. AtATG8A+AIM *mut* Control) on the y-axis. The centroid for x-axis and y-axis is represented as dashed lines. Dot size is mapped to reflect significant induction in heat-treated samples represented as – Log2 (*P* value). The 172 AtATG8A associated and outcompeted proteins are colored based on their response to heat stress (red, induced; blue, not induced). **(C) AtATG8E proxitome upon HS or HS+CB treatments.** Protein abundance pattern represented by a heatmap (Log2 (Mean PSM+1) – Mean PSM per protein) for the 40 AtATG8E proximal proteins identified as induced upon either HS or HS+CB. Each column represents the row-scaled mean PSM count of 3 independent biological replicates. **(D) Comparison between AtATG8E proxitome and interactome.** Venn diagram of two overlapping pairwise comparisons. AtATG8E-TID proximal proteins (beige circle) are defined as in Fig. S10A. AtATG8E associated and outcompeted proteins (purple circle) are defined as AIM-dependent in Fig. S8E. **(E) AtCESAR2 co-localizes with AtATG8E decorated autophagosomes.** Confocal microscopy images of *A. thaliana* root epidermal cells at the transition zone co-expressing AtCESAR2-GFP with mCherry- AtATG8E. 5-d-old *A. thaliana* seedlings were grown in 1% agar ½ MS + MES + 1% sucrose plates and incubated at 21°C (Control, C) or 37°C (Heat stress, HS) for 3h followed by a recovery of 3h at 21°C prior to imaging. Representative images of 12 biological replicates are shown. Area highlighted in the white-boxed region in the merge panel was further enlarged and presented in the inset panel. Scale bars, 10 μm. Inset scale bars, 10 μm. **(F) AtCESAR2 localization to autophagosomes increases upon HS.** Quantification of confocal experiments in Fig. 3E as assessed by the Mander’s colocalization coefficients between AtCESAR2-GFP and mCherry-AtATG8E. M1, fraction of AtCESAR2-GFP signal that overlaps with mCherry-AtATG8E signal. M2, fraction of mCherry-AtATG8E signal that overlaps with AtCESAR2-GFP signal. Bars indicate the mean ± SD of 12 biological replicates. Two-tailed Mann-Whitney test was performed to analyse the significance of the differences of M1 and M2 values between Control (21°C) and HS (37°C) conditions. ****, *P* value < 0.0001. **(G) AtCESAR2 localizes inside of autophagosomes.** Transmission electron microscopy (TEM) micrograph showing either immune-gold labelled AtCESAR2-GFP (15 nm gold particles) or mCherry-AtATG8E (10 nm gold particles), at the autophagosomes in *A. thaliana* root cells. 5-d-old *A. thaliana* seedlings were grown at 21°C in 1% agar ½ MS + MES + 1% sucrose plates before cryofixation. Sections from AtCESAR2-GFP and mCherry-AtATG8E co-expressing samples were labelled with anti-GFP and anti-mCherry primary antibodies and secondary antibodies conjugated with either 15 nm gold particles (for GFP, green arrows) or 10 nm gold particles (for mCherry, magenta arrows). Scale bars, 200 nm. AP, autophagosome. MVB, multivesicular body. **(H) AtCESAR2 co-immunoprecipitates with AtATG8E *in vivo*.** RFP-Trap co-immunoprecipitation coupled to peptide competition of whole-seedling extracts of 7-d-old *A. thaliana* seedlings co-expressing AtCESAR2-GFP with either mCherry-AtATG8E or mCherry-EV (mCherry). Seedlings were incubated either at 21°C (Control, C) or 37°C (Heat stress, HS) for 6h followed by a 4h recovery at 21°C in fresh ½ MS + MES + 1% sucrose media. AIM *wt* and AIM *mut* peptides were added to a final concentration of 100 μM. Protein extracts were immunoblotted with anti-GFP and anti-RFP antibodies. Two independent biological replicates are shown **in** Fig. S14. **(I) AtCESAR2 undergoes autophagic degradation.** Confocal microscopy images of *A. thaliana* root epidermal cells at the elongation zone co-expressing AtCESAR2-GFP with mCherry-AtATG8E in wild-*type* (WT) and *atg5* mutant backgrounds. 5-d-old *A. thaliana* seedlings were incubated at 21°C (Control, C) or 37°C (Heat stress, HS) for 3h followed by a 3h recovery at 21°C in fresh ½ MS + MES + 1% Sucrose media ± 1 μM Concanamycin A (ConA). Representative images are shown. Scale bars, 10 μm. **(J) The rate of autophagic degradation decreases upon HS.** Quantification of the number of AtCESAR2-GFP puncta inside the vacuole (upper panel) or in the cytoplasm (lower panel) per normalized area (10,000 μm^2^). Bars indicate the mean ± SD of 15 biological replicates for Control ± ConA in WT; 17 for HS –ConA in WT; 18 for HS + ConA in WT; and 8 for HS +ConA in *atg5*. Two-tailed unpaired Student’s t-tests with Welch’s corrections were performed to analyse the differences of puncta ratio numbers between control and heat stress recovery conditions. **, *P* value < 0.01 (0.0017). **(K) AtCESAR2 undergoes autophagic degradation during prolonged recovery post-HS.** Western blots showing AtCESAR2-GFP (upper panel) and GFP-AtATG8A (lower panel) autophagic flux assays during heat stress recovery. *A. thaliana* seedlings were grown in ½ MS + MES + 1% sucrose media for 7 days before incubation at 21°C (Control, C) or 37°C (Heat stress, HS) for 4h, followed by either a 4h or a 20h recovery in fresh media ± 1 μM ConA at 21°C. 20 μg of total protein extract was loaded and immunoblotted with the indicated antibodies. Two independent biological replicates are shown (see Fig. S15C). Reference protein sizes are indicated on the left side of the blots (unit: kDa). Total protein loading control was analysed by Amidoblack (AB) staining.

Next, we generated *A. thaliana* transgenic lines co-expressing fluorescently tagged AtCESAR1 and AtCESAR2 and determined their subcellular localization. Live-cell imaging studies revealed that AtCESAR1-mCherry and AtCESAR2-GFP colocalized in puncta in *A*. *thaliana* root epidermal cells (Figure S11). Since they showed high degree of colocalization, we focused on AtCESAR2 as a representative member for further characterization. To test if AtCESAR2 localizes to autophagosomes, we performed confocal microscopy on transgenic lines co-expressing AtCESAR2-GFP and mCherry-AtATG8E. Already under control conditions, some AtCESAR2-GFP puncta colocalized with mCherry-AtATG8E-labelled autophagosomes (Figures 3E, 3F, S12A). Upon heat stress, the degree of colocalization further increased, suggesting that CESAR autophagy flux is heightened upon heat-induced proteotoxic stress (Figures 3E, 3F). MpCESAR also co-localized with MpATG8 puncta in *M*. *polymorpha*, consistent with the evolutionary conserved nature of CESAR-ATG8 interaction (Figure S12B).

To visualize AtCESAR2 puncta at ultrastructural level, we performed immunogold labelling transmission electron microscopy with two different sizes of gold particles (15nm for AtCESAR2-GFP and 10nm for mCherry-AtATG8E). Electron micrographs obtained from Arabidopsis root cell sections confirmed that AtCESAR2 colocalizes at double membraned AtATG8E labelled autophagosomes, consistent with our live-cell imaging results (Figures 3G, S13).

To orthogonally validate AtCESAR2-AtATG8 association, we performed co-immunoprecipitation (Co-IP) experiments under control and heat stress conditions. This approach confirmed that AtCESAR2 associates with AtATG8E in an AIM-dependent manner and the interaction is further strengthened upon heat stress treatments, similar to NBR1 (Figures 3H, S14). Altogether, these results demonstrate that CESAR recruitment to autophagosomes is increased upon heat stress.

We then tested if CESAR undergoes autophagic degradation. Inhibition of vacuolar degradation by Concanamycin A (ConA) under control conditions led to AtCESAR2-GFP-decorated autophagic bodies in *wild*-type root cells (Figure 3I). ConA treatment also induced the stabilization of AtCESAR2-GFP puncta during heat stress (Figure 3I). Notably, the ratio of vacuolar puncta to total puncta per normalized area was higher in control samples compared to those under heat stress (Figure 3J, upper panel), while the cytoplasmic/total puncta ratio was higher in heat stress samples (Figure 3J, lower panel). These findings suggest that AtCESAR2-GFP undergoes vacuolar degradation and its autophagic flux is slower during heat stress. We further assessed AtCESAR2-GFP levels in autophagy-deficient mutant *atg5*. We could not detect any AtCESAR2 vacuolar puncta, confirming that AtCESAR2 delivery to the vacuole depends on the autophagy machinery (Figures 3I and 3J).

To support our microscopy-based flux assays, we performed western blot-based autophagic flux assays. Initially, we determined how AtCESAR2 protein levels change under different heat stress conditions (Figure S15A). We observed that heat stress stabilized the full-length AtCESAR2-GFP after 1 hour, reaching maximum levels at 4 hours. The stabilization was further enhanced following 4 hours of recovery in control conditions (Figures S15A and S15B). When we performed overnight recovery experiments, we detected a free GFP band, indicating vacuolar degradation. Consistently, the free GFP band was stabilized upon ConA treatment and absent in the *atg5* mutant (Figures 3K, S15C). Comparable kinetics were observed for NBR1 and GFP-AtATG8A autophagic flux (Figures 3K, S15C). In summary, these results demonstrate that AtCESAR2 localizes to autophagosomes and undergoes autophagic degradation. Furthermore, heat stress induces the accumulation of AtCESAR2, which then undergoes autophagic degradation during recovery.

### Proteasome inhibition induces AtCESAR2 autophagic flux

Given that heat stress disrupts protein homeostasis by promoting protein misfolding in plants, we investigated whether inducing proteotoxic stress through proteasome inhibition would similarly affect the autophagic flux of AtCESAR2-GFP. We performed time-course experiments using the proteasome inhibitor Bortezomib. Strikingly, 50 µM Bortezomib treatments induced a rapid degradation of AtCESAR2-GFP, reaching its peak after 4 hours of incubation followed by 4 hours of recovery (Figures S16A, S16B). To determine whether this degradation is autophagy-dependent, we combined Bortezomib treatments with ConA treatment in *wild-*type and *atg5* mutant plants. Remarkably, in the *wild-type* background, both the total number of puncta per normalized area as well as the puncta size significantly decreased upon Bortezomib treatment (Figures S16C-E). In contrast, in the *atg5* mutant background, the total number of puncta did not change, while the puncta size increased significantly. This coincided with the lack of free GFP visible by blotting method in the autophagic flux assay, suggesting that impaired vacuolar delivery led to stabilization of enlarged cytoplasmic CESAR2 puncta (Figure S17). Finally, we tested if CB treatment could also induce AtCESAR2 flux. A time-course showed a significant increase in AtCESAR2 flux, consistent with our proteasome inhibition assays (Figure S18). In summary, these findings indicate that the autophagic degradation of AtCESAR2 is induced upon proteotoxic stress.

### CESAR targets hydrophobic ubiquitinated proteins for autophagic degradation upon proteotoxic stress

We aimed to define the cargo and mechanism of action of CESAR-mediated autophagy. AF2-based structural predictions revealed a highly conserved three-helix bundle domain that is characteristic for CUE (Coupling of Ubiquitin to ER degradation) domains^28^ on the N-terminal region (Figures 4A, S19A). Since previous studies have demonstrated that CUE domains from different organisms could bind ubiquitin^28–30^, we hypothesized that CESAR functions as an aggrephagy receptor and degrades ubiquitinated proteins. To test our hypothesis, we first performed *in vitro* pulldown assays using MpCESAR, since there is only one CESAR protein in *M. polymorpha*. Truncation and structure-guided mutagenesis experiments revealed that MpCESAR interacts with monoubiquitin and M1-linked di- and tetra-ubiquitin chains in a CUE domain-dependent manner (Figures 4B-D, S19B). To test for specificity towards different linked ubiquitin chains, we purified MpCESAR CUE domain and showed it binds to 7 different linkage types (M1, K6, K11, K29, K33, K48, and K63), indicating no clear specificity (Figure 4E).

**Figure 4.**
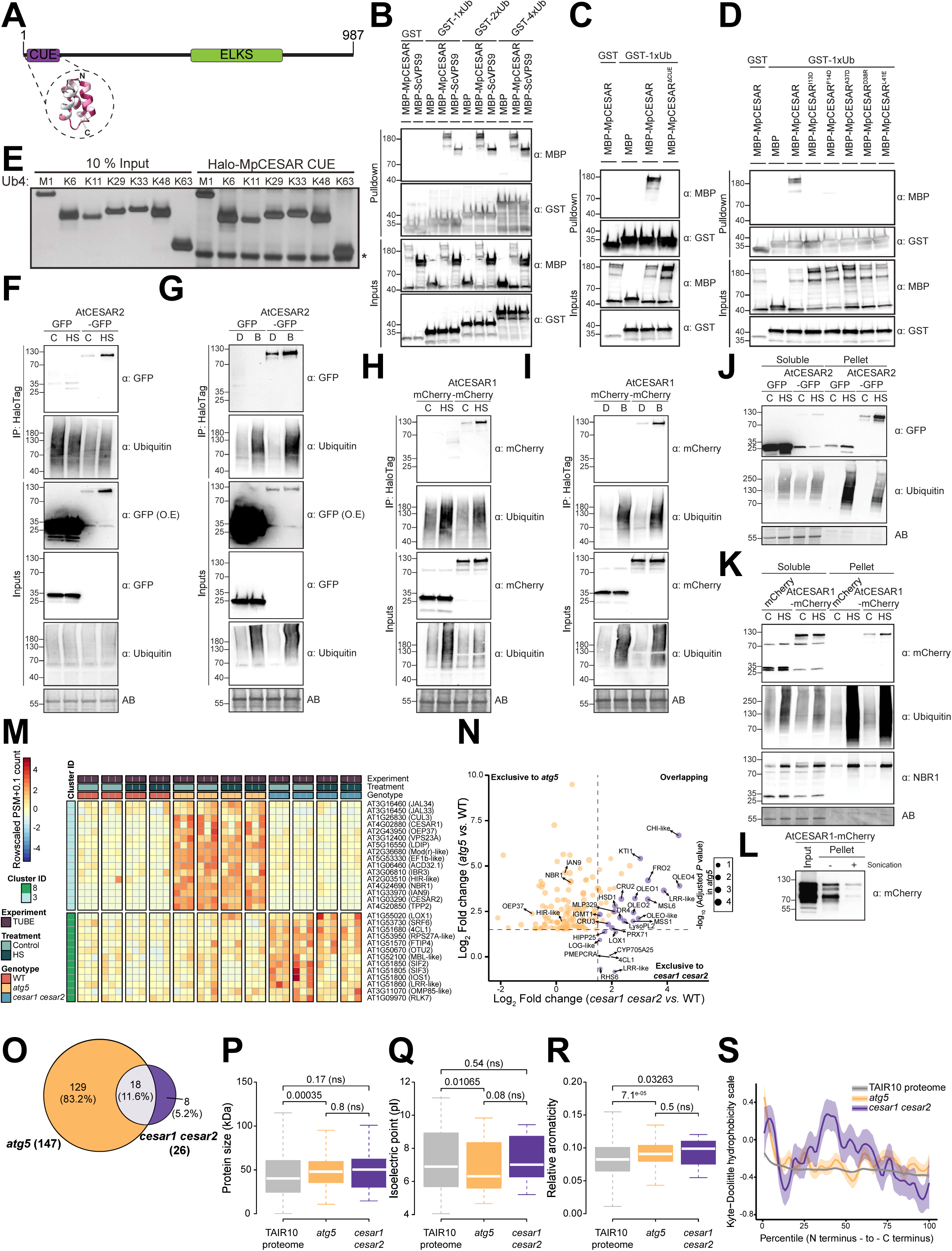
CESAR binds ubiquitin and targets ubiquitinated-proteins for autophagic degradation. **(A) CESAR has a conserved CUE domain.** Protein domain architecture of MpCESAR. The CUE and ELKS-Rab6-interacting/CAST family member 1 (ELKS) domains are highlighted in purple and green, respectively. A schematic representation of the conservation-mapped AF2-predicted model for the CUE domain shown in Fig. S19A is highlighted. **(B) MpCESAR binds ubiquitin *in vitro*.** *S. cerevisiae* (Sc) VPS9 was used as positive control. **(C-D) MpCESAR CUE domain is necessary for ubiquitin binding.** MpCESAR^ΔCUE^=MpCESAR^47–987^. Bacterial lysates containing recombinant protein were mixed and pulled down with glutathione magnetic agarose beads. Input and bound proteins were visualized by immunoblotting with anti-GST or anti-MBP antibodies. **(E) MpCESAR CUE binds different ubiquitin chain linkages with similar affinities.** Halo-tagged MpCESAR CUE coupled to HaloLink resin was incubated with tetra-ubiquitin (Ub4) of the indicated linkage types. The captured materials were separated on 4-12% SDS-PAGE gel and silver stained. The asterisk indicates non-specific bands from Halo-MpCESAR CUE which have a similar electrophoretic mobility as K63-Ub4 chains. **(F-G) Proteotoxic stress enhances AtCESAR2 association with TUBEs.** 5-d-old *A. thaliana* seedlings expressing GFP-EV (GFP) or AtCESAR2-GFP in wild*-type* (Col-0) background were incubated in liquid ½ MS medium with 1% sucrose for 4h at 21°C (Control, C) or 37°C (Heat stress, HS) followed by 4h recovery phase at 21°C (panel F) or for 1h at 21°C in DMSO (D)-supplemented or 5 µM Bortezomib (B)-supplemented media followed by 1h recovery phase in fresh media (panel G) and used for co-immunoprecipitation. Plant lysates were incubated with Magne® HaloTag ® Beads conjugated with HaloTag-TUBE. Input and bound proteins were detected by immunoblotting using the respective antibodies as indicated. Total protein loading control was analysed by Amidoblack (AB) staining. Immunoblotting for bait is shown in Fig. S19E (related to Fig. 4F) and Fig. S19F (related to Fig. 4G). **(H-I) Proteotoxic stress enhances AtCESAR1 association with TUBEs.** 5-d-old *A. thaliana* seedlings expressing mCherry-EV (mCherry) or AtCESAR1-mCherry in wild*-type* (Col-0) background were incubated in liquid ½ MS medium with 1% sucrose for 4h at 21°C (Control, C) or 37°C (Heat stress, HS) followed by 4h recovery phase at 21°C (panel H) or for 1h at 21°C in DMSO (D)-supplemented or 5 µM Bortezomib (B)-supplemented media followed by 1h recovery phase in fresh media (panel I) and used for co-immunoprecipitation. Plant lysates were incubated with Magne® HaloTag ® Beads conjugated with HaloTag-TUBE. Input and bound proteins were detected by immunoblotting using the respective antibodies as indicated. Total protein loading control was analysed by Amidoblack (AB) staining. Immunoblotting for bait is shown in Fig. S19G (related to Fig. 4H) and Fig. S19H (related to Fig. 4I). **(J-K) HS increases CESAR localization to the insoluble fraction.** 5-d-old *A. thaliana* seedlings expressing either GFP-EV (GFP) or AtCESAR2-GFP (panel J), either mCherry-EV (mCherry) or AtCESAR1-mCherry (panel K) in wild*-type* (Col-0) background were incubated in liquid ½ MS medium with 1% sucrose for 4h at 21°C (Control, C) or 37°C (Heat stress, HS) followed by 4h recovery phase at 21°C. Soluble and pellet fractions were separated by centrifugation and normalized before immunoblotting using the respective antibodies as indicated. Total protein loading control was analysed by Amidoblack (AB) staining. **(L) CESAR partitioning to the insoluble fraction is reversible.** 5-d-old *A. thaliana* seedlings expressing AtCESAR1-mCherry in wild*-type* (Col-0) background were incubated in liquid ½ MS medium with 1% sucrose for 4h at 37°C followed by a 4**-**hour recovery phase at 21°C. Soluble (input) and pellet fractions were separated by centrifugation, and pellet fraction was further sonicated in a water bath before immunoblotting using the respective antibodies as indicated. **(M) TUBE interactors are differentially enriched in *atg5* and *cesar1 cesar2*.** Protein abundance pattern represented by a heatmap as (Log2 (PSM+0.1) – mean PSM value per protein) for the 16 and 13 proteins belonging to clusters 8 and 3 in Fig. S20B, respectively. A full caption of the heatmap is shown in Fig. S20C. Rows were clustered using Euclidean distance and resulting dendrograms are omitted from the figures. **(N) Insoluble proteins accumulate in the pellet fraction upon heat stress recovery.** Dot plot representing protein abundance as Log2 (Fold change) for the ratio of two pairwise comparisons, *cesar1 cesar2 vs.* WT (x-axis) and *atg5 vs.* WT (y-axis). Dot size is mapped to reflect significant enrichment in *atg5* represented as – Log10 (Adjusted *P* value). Enriched proteins in *cesar1 cesar2 vs.* WT or *atg5 vs.* WT are colored in purple and yellow, respectively. **(O) Comparison between *atg5* and *cesar1 cesar2* pellet-enriched fractions.** Venn diagram of the two overlapping pairwise comparisons in Fig. 4M comparing proteins enriched in *atg5* (yellow circle) or *cesar1 cesar2* (purple circle) pellet fractions. **(P-R) Analysis of protein features in the pellet-enriched fractions of *atg5* and *cesar1 cesar2*.** Boxplots representing protein size (P), isoelectric point (Q) and relative aromaticity (R) of the indicated genotypes. TAIR10 proteome was used as a reference. Horizontal white lines within the boxes indicate the median and the top and bottom edges of the boxes represent the 3^rd^ and 1^st^ quartile, respectively. Vertical colored dashed lines indicate 1.5x the interquartile range. *P* values derived from pairwise comparisons by Wilcoxon-Mann-Whitney test are shown. ns, not significant. **(S) Enriched proteins in *cesar1 cesar2* pellets are highly hydrophobic.** Line plots representing the hydrophobicity profiles along the protein sequence of pellet-enriched proteins in the indicated genotypes. The solid line represents the mean hydrophobicity and the shaded are represents the standard deviation. TAIR10 proteome was used as a reference. Protein hydrophobicity levels as defined by the Kyle-Doolittle scale is shown on the y-axis and proteins as defined by percentiles of its total sequence length are shown on the x-axis.

Next, we probed whether CESAR associates with ubiquitinated proteins *in vivo*, using the Tandem Ubiquitin Binding Entities (TUBE) technology^31^. For this assay, we sought to compare both heat stress (HS) and Bortezomib as treatment conditions. To this end, we optimized a short-term, low Bortezomib treatment condition that did not lead to acute AtCESAR2-GFP degradation yet stabilized *in vivo* polyubiquitin chains (Figures S19C, S19D). TUBE pull downs showed that both AtCESAR1 and AtCESAR2 associated with soluble ubiquitinated species, and their association increased upon stress treatments (Figures 4F-I, S19E-H). As expected, we could also detect an association between NBR1 and the TUBEs (Figure S19E-H). Altogether, these experiments demonstrate that CESAR interacts with ubiquitin via its CUE domain and is recruited to ubiquitinated proteins upon proteotoxic stress.

Since the inhibition of the proteasome leads to the accumulation of misfolded proteins into protein aggregates^32^, we next determined if CESAR could be associating with insoluble ubiquitinated species. We fractionated soluble and insoluble protein complexes using ultracentrifugation and tested CESAR partitioning using western blots. These assays demonstrated that both AtCESAR paralogs are in the insoluble fraction upon heat stress alongside ubiquitinated proteins and NBR1 (Figures 4J, K). Importantly, this is reversible and indicates an active recruitment of CESAR rather than aggregation (Figure 4L). Altogether, these results suggest CESAR could target insoluble ubiquitinated protein aggregates for autophagic degradation.

To identify potential ubiquitinated proteins that are degraded by CESAR, we generated a double CESAR knockout (*cesar1 cesar2*) in *A*. *thaliana* and independently assessed the soluble and insoluble fractions of Col-0, *atg5* and *cesar1 cesar2* upon proteotoxic stress. To study the soluble ubiquitinome, we performed TUBE AP-MS upon heat stress and proteasome inhibition and identified in total 372 proteins as TUBE interactors (Figures S20A, S21A, S21B, Tables S12, S13). We found NBR1 and both AtCESAR homologs enriched in *atg5*, further supporting our previous observations. However, we did not observe any proteins that were shared between *atg5* and *cesar1 cesar2* (Figures 4M, S20B, S20C, S21C, S21D), suggesting TUBE is more suited to identify soluble ubiquitinated proteins and not appropriate for identification of CESAR cargo. Indeed, TMT-based quantitative proteomics of total protein content showed most of these proteins are enriched in the proteome of *cesar1 cesar2* but not *atg5* (Figure S21E). This prompted us to dissect the proteome of the insoluble fraction. To this end, we performed MS analyses of the insoluble fraction upon heat stress. We identified 26 proteins enriched in *cesar1 cesar2* pellets, of which 18 were also enriched in *atg5* pellets (Figures 4N, 4O, Table S14). Among them, we found several members of the OLEOSIN family proteins, that have been reported to be ubiquitinated^33,34^. To gain knowledge on the properties of CESAR cargo, we studied different protein features such as protein size, isoelectric point (pI) and frequency of aromatic residues (aromaticity) in *cesar1 cesar2* pellet-enriched proteins. There were no significant differences in these properties between *atg5* and *cesar1 cesar2* proteins enriched in the pellet fraction (Figures 4P-R). However, *cesar1 cesar2* insoluble proteins had greater hydrophobicity compared to the whole proteome and *atg5* pellet-enriched proteins (Figure 4S). Altogether, our data suggests that CESAR associates with insoluble aggregates enriched in hydrophobic patches and marked by ubiquitin to mediate their autophagic clearance.

### CESAR is crucial for proteotoxic stress tolerance

Heat stress and erratic weather patterns are defining features of global climate change^35^. Next, we sought to determine the physiological significance of CESAR-mediated autophagy during heat stress. First, we tested the sensitivity of *cesar1 cesar2* to nutrient starvation, since starvation triggers bulk degradation rather than selective autophagy^1^. In contrast to *atg5* mutant where all autophagy pathways are inhibited, *cesar1 cesar2* mutant was not sensitive to either carbon or nitrogen starvation (Figures 5A, S22A). We then tested the sensitivity of *cesar1 cesar2* to two different proteotoxic stress conditions: heat stress and proteasome inhibition. We included *nbr1* and *atg5* lines as positive controls. Already under control conditions, *cesar1 cesar2* plants were smaller than the *wild*-type, suggesting CESAR-mediated autophagy already plays an important homeostatic role (Figures 5B, 5C, left panel). Following heat stress, the rosette area of *cesar1 cesar2* plants was reduced by 3.3-fold compared to control plants (Figures 5B, 5C, right panel). Furthermore, *cesar1 cesar2* plants were more sensitive than *nbr1* (Figures 5B, 5C, right panel). These findings indicate that CESAR is important for heat stress tolerance.

**Figure 5.**
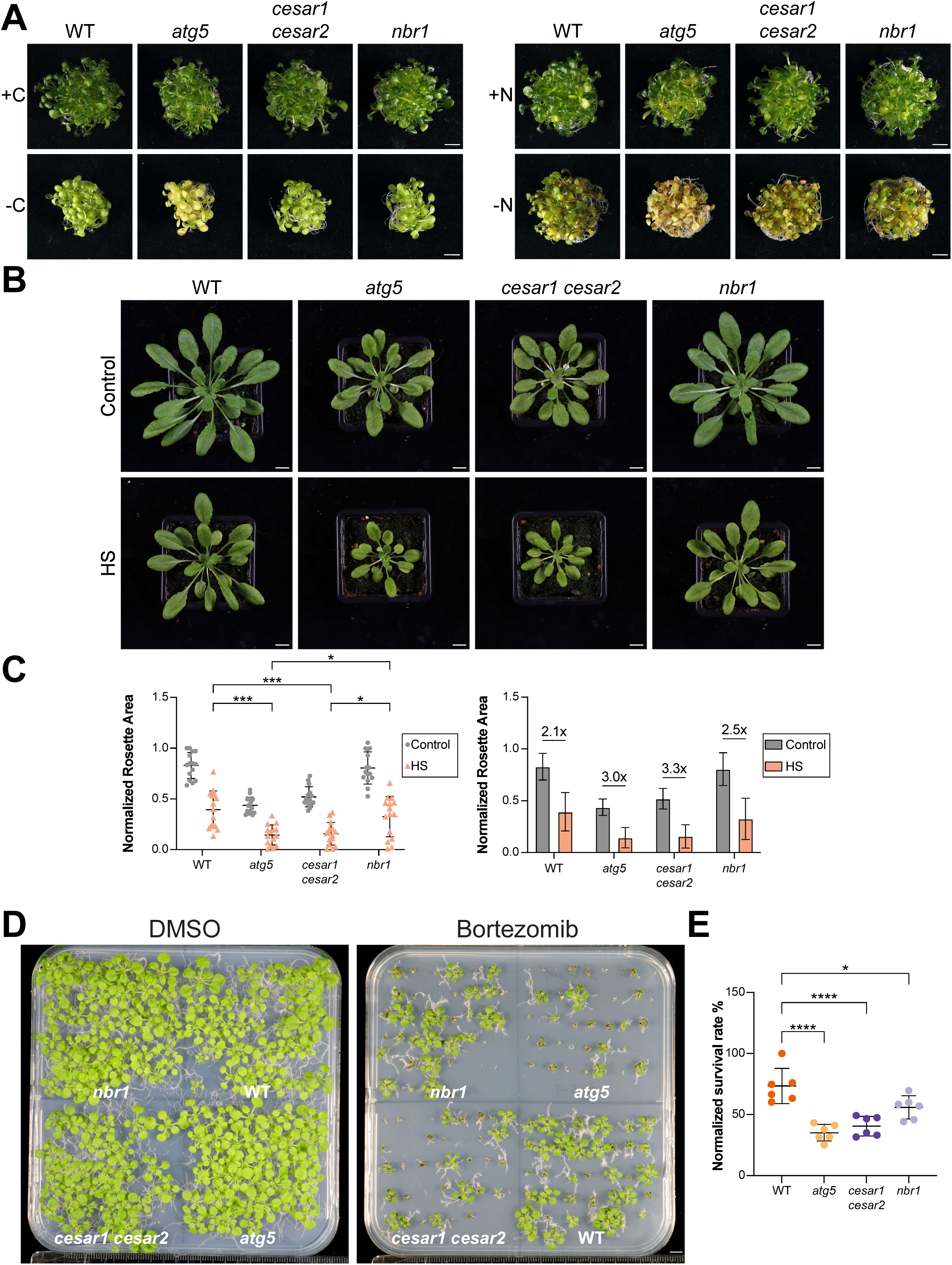
CESAR is essential for proteotoxic stress tolerance. **(A)** *cesar1 cesar2* plants are not hypersensitive to carbon or nitrogen starvation. 9-d-old *A. thaliana* seedlings of the indicated genotypes were grown in ½ MS + MES + 1% sucrose for 9 days, followed by either 4 days of carbon starvation (-C, left) or 6 days of nitrogen starvation (-N, right), respectively. 13-d-old and 15-d-old seedlings are shown in the images. Representative images of 3 independent biological replicates (see Fig. S22A) per genotype are shown. Scale bar, 1 cm. (B) *cesar1 cesar2* plants are sensitive to HS. *A. thaliana* plants of the indicated genotypes were grown on soil under a 8-h light/16-h dark photoperiod at 21°C for 3 weeks before incubation at 21°C (Control) or 37°C (Heat stress, HS) for 3 days without watering, followed by a 18day recovery at 21°C, after which they were imaged. 42-d-old plants are shown in the images. Representative images per genotype are shown. Scale bar, 1 cm. **(C) Rosette area quantification and statistical analysis.** Left panel, rosette areas were measured for each plant and normalized to the maximum rosette area value of wild-type (Col-0) at 21°C. Ordinary-one-way Anova with Tukey’s multiple comparisons test was performed to assess the differences in the normalized Rosette Area between genotypes at 37°C (Heat stress, HS). ***, Adjusted *P* value < 0.001 (0.0003 for WT vs *atg5*; 0.0006 for WT vs *cesar1 cesar2*); *, Adjusted *P* value < 0.05 (0.0121 for *atg5* vs *nbr1*; 0.0219 for *cesar1 cesar2* vs *nbr1*), non-significant differences are not shown. Right panel, size factor difference between the normalized rosette area for the indicated genotypes. (D) *cesar1 cesar2* plants are sensitive to proteasome inhibition. *A. thaliana* seedlings of the indicated genotypes were grown in 1% agar ½ MS + MES + 1% sucrose plates with either DMSO (left plate) or 3.75 µM Bortezomib (right plate) for 18 days before imaging. Representative images of 6 independent biological replicates are shown (n = 40 seeds per genotype per replicate). Scale bar, 1 cm. **(E) Quantification of the survival rate of seedlings grown in Bortezomib-containing plates.** Normalized survival rate for the replicate of Bortezomib plate assays shown in Fig. 5D. Survival rate of each replicate was calculated by dividing the number of seedlings that showed a phenotype of size bigger than 0.3 cm and green colour by the total number of seeds sown per genotype (40) and normalized to the highest survival value for the wild-type (Col-0) background of all the 6 biological replicates (n = 240, each dot represents a replicate with 40 seeds per genotype per plate)). Ordinary One-way Anova with Tukey’s multiple comparisons test was performed to assess the differences between the survival rates of the different phenotypes. ****, Adjusted *P* value < 0.0001; *, Adjusted *P* value < 0.05 (0.0322). Non-significant differences are not shown.

Given the established crosstalk between the Ubiquitin Proteasome System and selective autophagy^36^, we further explored whether the proteasome shows compensatory activation during heat stress in *cesar1 cesar2* plants. To this end, we measured proteasome activity in the *cesar1 cesar2* mutant under control conditions and following heat stress. Our results demonstrated that, compared to the *wild*-type background, *cesar1 cesar2* mutant exhibited constitutively increased trypsin-like proteasome activity (Figure S22B). Altogether, these results suggest that lack of CESAR leads to increased protein aggregate accumulation, which induces the proteasome.

To further corroborate our proteasome activity assays, we examined the sensitivity of *cesar1 cesar2* mutant to proteasome inhibition. The survival rates of *cesar1 cesar2* double mutant and the *atg5* mutant were severely compromised on Bortezomib plates compared to *wild*-type plants. The survival of the *nbr1* mutant was also significantly affected, albeit to a lesser extent (Figures 5D, 5E, S22C, S22D).

Overall, our findings indicate that CESAR-mediated autophagy plays a vital role in plant survival under proteotoxic stress, ensuring proteostasis by orchestrating the timely degradation of ubiquitinated hydrophobic proteins.

## Discussion

### Selective Autophagy: A Dynamic, Modular Pathway Sculpted by Evolution

Eukaryotic cells rely on sophisticated degradation systems to maintain homeostasis, particularly under stress^37^. Among these systems, autophagy plays a pivotal role, with disruptions linked to a range of diseases and aging, highlighting its critical function in cellular adaptation and stress management^38,39^. Autophagy has evolved as a modular pathway, where core machinery drives autophagosome formation, and selective autophagy receptors (SARs) target specific cargo for degradation^23,40^. Despite their central role, the SAR inventory in plants remains incomplete, with only a few SARs characterized, underscoring the need for a more comprehensive exploration^41^. Moreover, no comparative analysis of SAR repertoires across species has been performed, leaving the evolutionary diversity of these critical receptors largely uncharted.

In this study, we address this gap by identifying a diverse suite of SAR candidates across five model organisms, providing a valuable resource for future discoveries. Our data suggest that SARs exhibit both conserved and lineage-specific features, reflecting evolutionary pressures unique to each species (Figure 2). Expanding the known SAR repertoire opens new avenues for understanding how selective autophagy adapts to diverse biological contexts and environmental stressors. Additionally, this work emphasizes the importance of investigating SARs in non-model organisms, particularly extremophiles or species with unique ecological niches. Such studies could reveal untapped autophagy pathways that have remained hidden, shedding light on the full diversity of cellular quality control mechanisms shaped by distinct evolutionary forces.

### CESAR: A Crucial Player in the Plant Proteostasis Network

As a proof of concept, we characterized CESAR, a highly conserved SAR across streptophytes. CESAR interacts with ATG8 in an AIM-dependent manner and undergoes autophagic degradation together with the bound cargo, confirming its role as a canonical SAR (Figure 3). Notably, CESAR contains a CUE domain, which has an affinity for the Isoleucine44 patch on ubiquitin, —a trait it shares with the UBA domain of the archetypal aggrephagy receptor NBR1^28,42^. However, unlike UBA domains, CUE domains can bind to non-ubiquitinated cargo, such as exposed hydrophobic surfaces, allowing CESAR to target a broader spectrum of substrates^43^. Additionally, the CUE domain was shown to undergo autoubiquitination^44^, which would attract further ubiquitin binding SARs to CESAR bound aggregates and increase the bulkiness of the CESAR-bound cargo—an attribute that has been suggested to facilitate efficient autophagic degradation^45^. Consistently, our TUBE pull-down assays have shown that CESAR and NBR1 are recruited to the same cargo, suggesting they may cooperate in clearing protein aggregates (Figure 4).

### Proteotoxic Stress Responses Are Not Uniform: Diverse Stresses Elicit Distinct Autophagic Pathways

Interestingly, CESAR exhibits different autophagic degradation kinetics depending on the type of stress. For instance, under heat stress, CESAR is stabilized, with autophagic degradation only initiated during prolonged recovery periods. In contrast, proteasome inhibition triggers rapid CESAR autophagic flux and reduces the number and size of CESAR puncta (Figure 3 and Figures S16-S17). A key distinction between these stressors is the much higher accumulation of polyubiquitinated proteins following proteasome inhibition.

We propose that CESAR may function as a sensor of aggregate burden within the cell. Under mild proteotoxic stress induced by our heat stress treatments, CESAR could rapidly sequester ubiquitinated aggregates, allowing the proteasome to process them efficiently and mitigate cytotoxicity. However, when the aggregate burden surpasses a critical threshold such as the inhibition of the proteasome, CESAR might undergo conformational changes that induce a rapid autophagic degradation. This dual-sensing mechanism would enable plant cells to coordinate the activity of different SARs targeting ubiquitinated proteins, effectively balancing the proteasome and autophagy pathways to maintain protein homeostasis (Figure 6).

**Figure 6.**
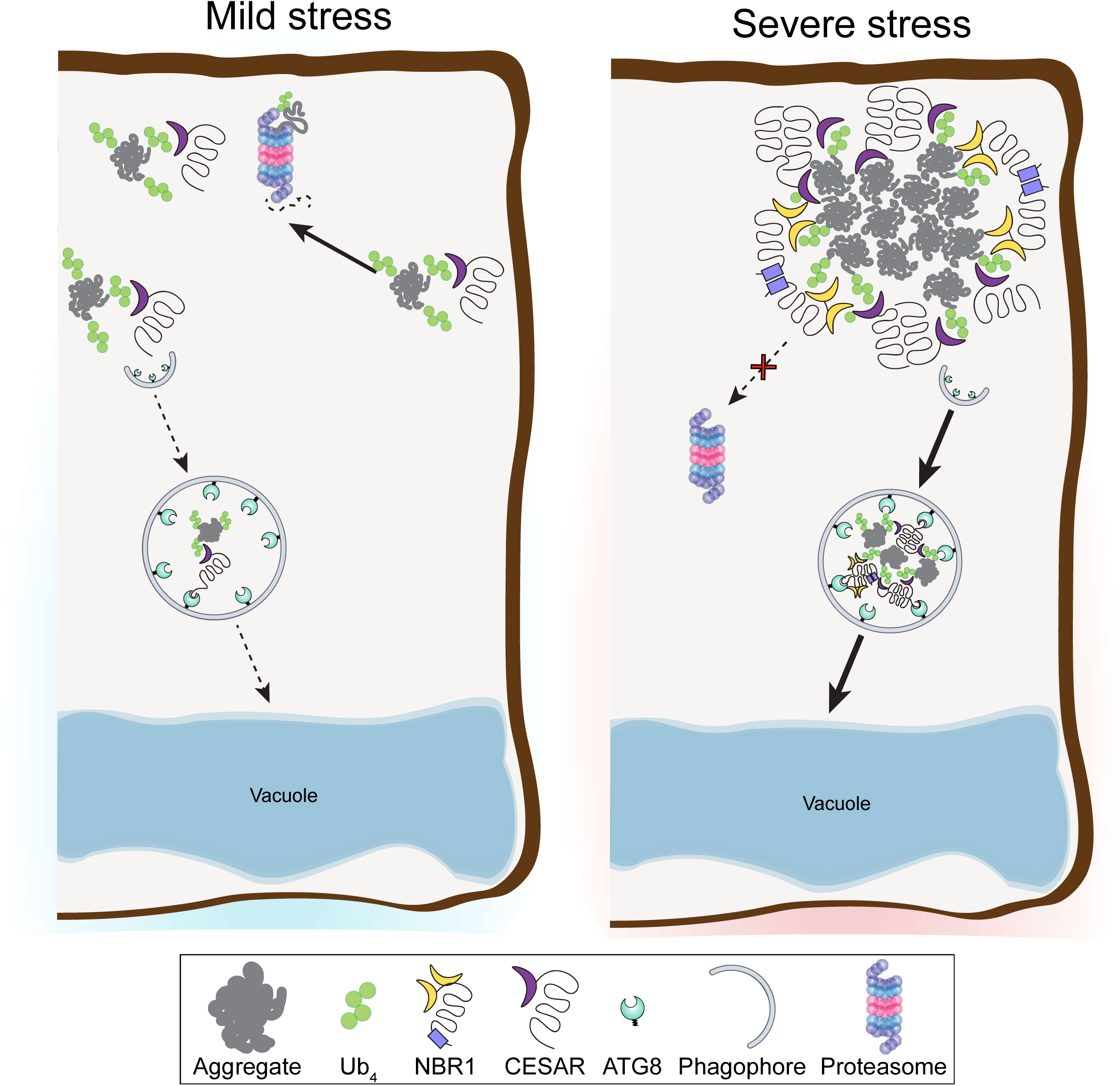
Current working model of CESAR-mediated autophagy. We propose that under mild stress conditions, CESAR binds and stabilizes ubiquitinated protein aggregates, allowing other components of the proteostasis network to process and recycle them (left panel). In response to severe stress conditions, CESAR undergoes conformational changes that activate its role in the autophagic clearance of protein aggregates (right panel). CESAR may also cooperate with NBR1 to facilitate the degradation of these aggregates via autophagy.

## Acknowledgements

We thank the Mass Spectrometry, Peptide Synthesis, BioOptics, Plant Sciences, Protein Technologies of The Vienna BioCenter Core Facilities. We also acknowledge HPC Cluster of Vienna Biocenter and the Crystallography Platform of the Institute of Molecular Pathology. We thank Takashi Ueda for sharing Marchantia *atg7* mutant and Sascha Martens for sharing tetraubiquitin plasmids. We thank the Dagdas lab for fruitful discussions.

## Author contributions

VSM performed molecular cloning, Western based experiments, protein purification, optimisation of crystallization conditions, GST-pulldown, Marchantia CoIPs, affinity purification and sample preparation for Mass Spectrometry, ITC experiments, phylogenetic analyses, analysed protein enrichment in MS/MS and TMT, made figures, wrote the draft and manuscript revisions. MN performed molecular cloning, plant treatments, Western based assays, phenotypic assays and subsequent analysis, confocal microscopy and subsequent analysis, Arabidopsis CoIPs, affinity purification and sample preparation for Mass Spectrometry, generated Arabidopsis transgenic lines, made figures, wrote the draft and manuscript revisions. MC performed molecular cloning of TurboID constructs and generated the plant lines, performed affinity purification and sample preparation for Mass Spectrometry, optimized sample preparation to Mass Spectrometry, analyzed protein enrichment in MS/MS and made figures. RP analyzed AF2 interactome, protein enrichment in MS/MS and general features of proteins identified by MS and contributed critically to the manuscript. VA performed Western based assays, plant treatments, fractionation experiments, sample preparation for Mass Spectrometry, made figures and contributed critically to the manuscript. PG performed root electron microscopy and made figures. JC performed phylogenetic analyses, crystallographic data refinement and made figures. VM performed phylogenetic analyses and made figures. LP performed crystallization optimization and harvested crystals. NG generated Arabidopsis CRISPR lines and performed plant selection. RK performed biotin labelling optimization, affinity purification and sample preparation for Mass Spectrometry. AA performed treatment optimization in Marchantia and generated Marchantia transgenic lines. HD performed confocal microscopy. GA performed protein purification and Ub-chain experiments. NB performed molecular cloning and GST-pulldown. AB performed treatment optimization and sample preparation for Mass Spectrometry. TC performed heat stress experiments. LA performed molecular cloning and generated Marchantia transgenic lines. AM performed treatment optimization in Marchantia and generated Marchantia transgenic lines. MG performed treatment optimization in Marchantia, affinity purification and sample preparation for Mass Spectrometry and generated Marchantia transgenic lines. MM performed molecular cloning and initial characterization of *cesar1* and *cesar2* mutants. TMR performed proteasome activity assays. CLW generated and characterized the Chlamydomonas transgenic strains and prepared the corresponding samples for affinity purification. JK prepared samples for affinity purification. AM performed crystallographic data collection and refinement. ER supervised all MS-related measurements. BH, IB, MP, SU, YK, TC, SR, and YD supervised their respective team members. SR secured funding. YD, secured funding, designed the project, wrote the manuscript. All authors revised the draft and accepted the manuscript submission.

## Data availability

The mass spectrometry proteomics data have been deposited to the ProteomeXchange Consortium via the PRIDE partner repository with the dataset identifiers PXD051966 and PXD055359. All the raw data are available via Zenodo (DOI:10.5281/zenodo.13714560). X-ray diffraction amplitudes and the final model was deposited in the RCSB (PDB:9F2M).

## METHODS

### Phylogenetic analyses

To build a phylogeny of ATG8, genome sequences of ATG8 homologs were collected from NCBI Nucleotide and the joint genome’s institute (JGI) Plant Comparative Genomics portal Phytozome^46^. Initially, protein sequences from *A. thaliana* ATG8 isoforms were used as queries to search predicted proteomes in the NCBI non-redundant (NCBI-nr) database using the blastp algorithm in BLAST^47^. Sequences containing the C-terminal glycine and covering all ATG8-specific α-helices and β-sheets were selected for further analysis, as previously described^48^. In total, 621 non-redundant sequences from 80 different plant and algal species were collected (Table S1). Sequences were aligned using Clustal Omega multiple sequence alignment (MSA) package^49^ and further reconstructed using Gblocks v 0.91.1^50^ to remove blocks of problematic alignment as implemented in http://phylogeny.lirmm.fr/phylo_cgi/. The resulting blocks were used to compute a Maximum-Likelihood (ML) phylogenetic tree using IQ-TREE v2.3.4^51^ and visualized using iTOL v6.9.1^52^.

To identify putative homologs of our autophagy receptor candidates across the tree of life, profile hidden Markov models (HMM) for each of the six proteins (MpCESAR, MpYHBC, MpRING1/2 and MpVWA41/43) were generated using HMMER v3.4 as implemented in http://hmmer.org. A taxonomically balanced set of Archaeplastida genome-predicted proteomes downloaded from UniProt^53^ was curated based on BUSCO scores^54^ as described in De la Concepción et *al.*^55^. Initially, protein sequences from *M. polymorpha* were used as queries to search against the curated proteome dataset. The resulting hits were extracted, aligned using MAFFT v7.525^56^ and trimmed with trimAl v1.4^57^. Multiple sequence alignments were visualized in AliView^58^. Next, ML phylogenetic trees were constructed for each protein using IQ-TREE v2.3.4^51^ and ModelFinder^59^ to infer the Best-fit model. Manual curation followed by reconstruction of phylogenies was carried out iteratively to remove long-branching taxa and paralogous genes. Ultimately, MpVWA41/43 and MpRING1/2 were combined into the same phylogeny as they were found to be recent gene duplications in *M. polymorpha*. Phylogenies were visualized in FigTree as implemented in http://tree.bio.ed.ac.uk/software/figtree/ and iTOL v6.9.1^52^. To classify the domains found within the individual proteins in our alignments, hmmscan was used to search against the Pfam-A HMM database^60^. The domain with the better conditional *E*-value was assigned to the region when multiple HMMs overlapped by more than 50%, and domains mapping with less than 25% HMM coverage were excluded. The domains were visualized in iTOL v6.9.1^52^. Comparison of aminoacid conservation at AIM motifs was performed using WebLogo3^61^.

OrthoFinder v2.5.5 was used to cluster the proteomes of our 5 species of interest (*C. reinhardtii*, *M. polymorpha*, *P. patens*, *N. benthamiana* and *A. thaliana*) into Phylogenetic Hierarchical Orthogroups (HOGs)^62^. UpSet plots were generated in RStudio v2023.12.1 using the complexUpset and ggplot2 libraries.

### Plant and algal material and molecular cloning procedures

Coding sequences from genes of interest were either obtained as synthetic DNA from Twist Biosciences or amplified from cDNA with primers listed in the materials section. The internal BsaI sites were mutated by site-directed-mutagenesis without affecting the amino acid sequence. Constructs for *M. polymorpha* were generated using either the Gateway^63^ or GreenGate^64^ cloning systems. Constructs for *A. thaliana*, *N. benthamiana* and *E. coli* transformation were assembled with the GreenGate cloning system^64^. Constructs for *P. patens* were generated using the polyethylene glycol (Sigma, P7181) transformation^65,66^. Constructs for *C. reinhardtii* were generated using the MoClo^67^ cloning system. All plasmids used in this study, along with the corresponding primers used for their generation, are listed in the materials section.

All *C. reinhardtii* strains used in this study originate from CC124 and are listed in the materials section. Transgenic *C. reinhardtii* 3XFLAG-Venus-ATG8 and 3XFLAG-cytVenus strains were generated by electroporation as previously described^68^. Transformants were isolated based on antibiotic resistance on solid Tris-Acetate-Phosphate (TAP) media containing 1.6% w/v agar and Hutner’s trace elements^69^. The correct expression and localization of 3XFLAG-cytVenus and 3XFLAG-Venus-ATG8 proteins in the transformants were validated via immunoblotting and confocal microscopy.

All *M. polymorpha* lines used in this study originate from Takaragaike-1 (Tak-1) and are listed in the materials section. Transgenic *M. polymorpha* plants were generated by *Agrobacterium*-mediated transformation of regenerated thalli as previously described^70^. Briefly, ∼4cm^2^ *M. polymorpha* thalli extracted from 14-d-old plants were co-incubated with *Agrobacterium* containing the desired constructs for 2 days in transformation media (1.5 g/L Gamborg B5 medium, 2.5 mM MES monohydrate, 20 g/L D-Sucrose, 1g/L Casein hydrolysate, 2 mM L-Glutamine) supplemented with 100 µM Acetosyringone. Transformants were selected based on antibiotic resistance on half-strength Gamborg’s B5 medium containing 1% agar without sucrose supplemented with Ticarcillin.

All *A. thaliana* lines used in this study originate from the Columbia (Col-0) ecotype and are listed in the materials section. The following lines: *atg5-1*^71^*, nbr1-2*^72^, pUBQ::mCherry^73^, pUBQ::mCherry-AtATG8A^16^, pUBQ::mCherry-AtATG8E^74^, pUBQ::mCherry-AtATG8E/*atg5-1*^16^ and pUBQ::GFP-AtATG8A^16^ have been previously described.

Double mutant *cesar1-1/cesar2-1* was obtained by genetic crossing of *cesar1-1* and *cesar2-1* T-DNA insertion lines from SALK (SALK_112206.23.80.x and SALK_022568 lines, respectively). Homozygous plants were obtained by genotyping PCR, primers used for genotyping are listed in the materials section.

Stable transgenic *A. thaliana* lines were generated by *Agrobacterium*-mediated transformation using the floral dip method^75^. The following lines: pRPS5::AtCESAR2-GFP in Col-0, pRPS5::AtCESAR2-GFP in pUBQ::mCherry, pRSP5::AtCESAR2-GFP in pUBQ::mCherry-AtATG8E, pRPS5::AtCESAR2-GFP in pUBQ::mCherry-AtATG8E/*atg5-1* were obtained by transformation of the indicated background genotypes with pGGSun-pRPS5::AtCESAR2-GFP vector. Transformants were selected on sulfadiazine-containing medium and F1 and F3 lines were screened for AtCESAR2-GFP protein expression. The following lines: pRPS5::AtCESAR1-mCherry in Col-0 and pRPS5::AtCESAR1-mCherry in pRPS5::AtCESAR2-GFP were obtained by transformation of the indicated background genotypes with pGGSun-pRPS5::AtCESAR1-mCherry vector. Transformants were selected on either sulfadiazine or hygromycin containing medium, respectively, and F1 and F3 lines were screened for AtCESAR1-mCherry protein expression. Stable lines of pAtATG8E::mCherry-TurboID-AtATG8E, pAtATG8E::mCherry-TurboID-AtATG8E^ADS^, pAtATG8E::mCherry-TurboID-AtATG8E^G118A^ and pAtATG8E:mCherry-TurboID were obtained by transformation of Col-0 plants. Transformants were selected on sulfadiazine-containing medium, F3 lines were screened for AtATG8E expression. pUBQ::GFP-AtATG8A/*atg5-1* was obtained by genetic crossing of the respective lines and the resulting seeds were selected on hygromycin and basta plates.

*A. P. patens* lines used in this study have been previously described^66^. Transient expression in *N. benthamiana* was carried out by *Agrobacterium*-mediated transformation^76^.

### Plant and algal growth conditions and treatments

*A. C. reinhardtii* strains were maintained as previously described^77^. Briefly, *C. reinhardtii* strains were grown in liquid TAP culture under 50 µmol m^-2^ s^-1^ light intensity at 22°C with shaking in the absence of antibiotics. For affinity purification, two cultures per construct (3XFLAG-Venus-ATG8 and 3XFLAG-cytVenus) were diluted to 3·10^6^ cells per ml and placed each under control conditions or in a water bath (GFL Shaking Water Bath 1086) under 50 µmol m^-2^ s^-1^ light intensity constant illumination at 37°C for 4 hours without shaking, followed by a 2-hour recovery under control conditions with shaking. Cultures were pooled together during harvesting by centrifugation at 4,000 x g for 5 min at 4°C.

The male *M. polymorpha* accession Takaragaike-1 (Tak-1) was maintained asexually and cultured through gemmae using half-strength Gamborg’s B5 medium containing 1% agar without sucrose under 50-60 mmol photons m^-2^ s^-1^ continuous white light at 22°C unless stated otherwise^70^. For autophagic flux assays, *M. polymorpha* gemmaes were cultured in liquid half-strength Gamborg’s B5 medium with 1% sucrose for 7 days without shaking. For Torin treatment, gemmalings were cultured on the last day in liquid half-strength Gamborg’s B5 medium with 1% sucrose supplemented with DMSO or 12 µM torin (Santa Cruz Biotechnology) for 5 hours with gentle shaking. For Heat stress (HS) treatment, gemmalings were cultured on the last day for 4 hours at 37°C in a temperature-controlled cabinet (Percival Scientific Series 101) without shaking, followed by a 3-hour recovery phase in liquid half-strength Gamborg’s B5 medium with 1% sucrose under control conditions with gentle shaking. When combined with E64d/pepstatin A treatments, gemmalings were incubated for additional 16 hours in liquid half-strength Gamborg’s B5 medium with 1% sucrose supplemented with 10 µM E64d (Santa Cruz Biotechnology) and 10 µM pepstatin A (Santa Cruz Biotechnology) with gentle shaking. For *M. polymorpha* confocal microscopy, asexual gemmae were incubated in half-strength Gamborg’s B5 medium for 2 days before imaging.

For affinity purification, *M. polymorpha* gemmaes were cultured in liquid half-strength Gamborg’s B5 medium with 1% sucrose for 10 days without shaking. For HS treatment, gemmalings were cultured on the last day for 4 hours at 37°C in a temperature-controlled cabinet (Percival Scientific Series 101) without shaking, followed by a 3-hour recovery phase in fresh liquid media supplemented with DMSO or 50 µM CB5083 under control conditions with gentle shaking.

For *A. thaliana* standard plant growth, seeds were vapor-phase sterilized (90% sodium hypochlorite and 10% HCl) and sown on a water-saturated peat-based substrate (Klasmann Substrat 2, Klasmann-Deilmann GmbH, Geeste, DE). A *Bacillus thuringiensis* solution was used as a preventative control against fungus gnats (Gnatrol, Valent BioSciences, Libertyville, IL, USA). Lighting was provided by a custom-designed, multi-channel LED system developed in collaboration with RHENAC GreenTec AG (Germany), optimized for uniformity. The plants were cultured at 21°C in a 16-hour light/8-hour dark photoperiod under 165 µmol m^-2^ s^-1^ (PPFD) light intensity, using the following LED channels: 400 nm, white 4500K, 660 nm, and 730 nm. This resulted in the following spectral light composition: 22% PPF-Blue (400-500 nm), 31% PPF-Green (500-600 nm), and 47% PPF-Red (600-700 nm). Additionally, far-red light (PPF-NIR; 700-780 nm) was supplemented at 17% of PPFD (red to far-red ratio 1:0.35).

For *in vitro* seedling growth, *A. thaliana* seeds were sterilized in 70% ethanol with 0.05% SDS for 15 minutes, rinsed twice with absolute ethanol, and air-dried on paper under sterile conditions.

All HS assays conducted in plants in this study were performed at 37°C in a temperature-controlled chamber (Percival CU-36L4, SN: 8287.01.06C) under fluorescent lamps with 140 µmol·m⁻²·s⁻¹ light intensity, 65% humidity.

For affinity purification and proximity labelling experiments, around 16 mg of seeds per genotype were sown in 250 mL Erlenmeyer flasks containing 90 mL of liquid ½ MS salts (Duchefa) with 1% sucrose medium and stratified in the dark at 4°C for 48 hours. After stratification, flasks were maintained in a 16-hour light/8-hour dark photoperiod with 50 µmol m^-2^ s^-1^ light intensity under constant gentle shaking for 7 days. For affinity purification, 7-d-old seedlings were incubated at 37°C for 6 hours, followed by a 4-hour recovery in fresh liquid ½ MS salts (Duchefa) with 1% sucrose medium with either 50 µM CB5083 (BIOCAT GmbH; dissolved in DMSO) or an equivalent volume of DMSO under control conditions. For proximity labelling experiments, 50 µM biotin (Sigma Aldrich) was added to the media during the recovery. Control samples were maintained under control conditions for 6 hours, followed by the addition of fresh liquid ½ MS salts (Duchefa) with 1% sucrose medium for additional 4 hours. For proximity labelling experiments, the same treatments as the affinity purification were followed and following treatment, the reaction was stopped with ice-cold water.

For protein extraction, co-immunoprecipitation, and protein content analysis, seeds were sown in 12-well plates containing liquid ½ MS salts (Duchefa) with 1% sucrose medium and stratified in the dark at 4°C for 48 hours. Following stratification, plates were maintained in a 16-hour light/8-hour dark photoperiod with 50 µmol m^-2^ s^-1^ light intensity under constant gentle shaking for 7 days.

For co-immunoprecipitation experiments, 7-d-old seedlings were incubated at 37°C for 6 hours, followed by a 4-hour recovery in fresh liquid ½ MS salts (Duchefa) with 1% sucrose medium with either 50 µM CB5083 (BIOCAT GmbH; dissolved in DMSO) or an equivalent volume of DMSO, under constant gentle shaking. Control samples were maintained under control conditions for 6 hours, followed by the addition of fresh liquid ½ MS salts (Duchefa) with 1% sucrose medium for additional 4 hours.

For protein content analysis upon HS, 7-d-old seedlings were incubated for 1 to 4 hours at 37°C followed by a 1 to 4-hour recovery under control conditions. For autophagic flux experiments upon HS, 7-d-old seedlings were incubated either under control conditions or at 37°C for 4 hours, followed by either a 4-hour recovery in fresh liquid ½ MS medium or a 20-hour recovery in liquid ½ MS medium containing 1 µM Concanamycin A (ConA; CAS 80890-47-7; Santa Cruz) or an equivalent volume of DMSO under control conditions. For TUBE-pulldown experiments, 7-d-old seedlings were incubated either under control conditions or at 37°C for 4 hours, followed by a 4-hour recovery in fresh liquid ½ MS medium under control conditions. For aggregate isolation and fractionation experiments, 7-d-old seedlings were incubated either under control conditions or at 37°C for 4 hours, followed by either a 4-hour or a 20h-recovery in fresh liquid ½ MS medium under control conditions.

For protein content analysis upon Bortezomib treatment, 7-d-old seedlings were incubated in liquid media with either DMSO or 50 µM of Bortezomib (Santa Cruz; CAS 179324-69-7) for 4 hours under control conditions, followed by a 1 to 4-hour recovery in fresh media. For autophagic flux experiments upon Bortezomib treatment, 7-d-old seedlings were incubated with either 50 µM of Bortezomib for 4 hours or an equal volume of DMSO under control conditions, followed by either a 4-hour recovery in fresh liquid ½ MS medium or a 20-hour recovery in fresh liquid ½ MS medium containing 1 µM Concanamycin A (ConA; CAS 80890-47-7; Santa Cruz) or an equal volume of DMSO. For TUBE-pulldown experiments, 7-d-old seedlings were incubated with either 5 µM of Bortezomib or an equal volume of DMSO for 1 hour under control conditions, followed by a 1-hour recovery in fresh liquid ½ MS medium.

For autophagic flux experiments upon CB5083 treatment, 7-d-old seedlings were incubated with either 50 µM of CB5083 (BIOCAT GmbH) for 4 hours or an equal volume of DMSO under control conditions, followed by either a 4-hour recovery in fresh liquid ½ MS medium or a 20-hour recovery in fresh liquid ½ MS medium containing 1 µM Concanamycin A (ConA; CAS 80890-47-7; Santa Cruz) or an equal volume of DMSO.

For electron and confocal microscopy, seeds were sown in a single line on ½ MS medium (Duchefa) containing 1% agar and 1% sucrose, then stratified in the dark at 4°C for 48 hours. Subsequently, seeds were grown under 50 µmol m^-2^ s^-1^ LED lighting with a 16-hour light/8-hour dark photoperiod for 5 days under control conditions. Plates were oriented vertically to facilitate root elongation along the medium surface. To assess co-localization during HS, 5-d-old seedlings were subjected to 37°C for 3 hours, followed by a 3-hour recovery under control conditions prior to imaging. For autophagic flux analysis during HS, seedlings were incubated at 37°C for 3 hours. Post-incubation, individual seedlings were transferred to fresh liquid ½ MS medium containing 1 µM Concanamycin A (dissolved in DMSO) or an equal volume of DMSO and allowed a 3-hour recovery under control conditions before imaging. For autophagic flux upon Bortezomib treatment recovery, 5-d-old seedlings were transferred to fresh liquid ½ MS medium with 50 µM Bortezomib (Santa Cruz; CAS 179324-69-7; dissolved in DMSO) or an equivalent volume of DMSO and incubated for 2 hours under control conditions. This was followed by a 2.5-hour recovery in ½ MS medium containing 1 µM Concanamycin A or an equivalent volume of DMSO, prior to imaging.

For proteasome activity assays, *A. thaliana* seeds of the indicated genotypes were sown on ½ MS medium (Duchefa) containing 0.8%, then stratified in the dark at 4°C for 72 hours. Subsequently, seeds were grown under 110 µmol m^-2^ s^-1^ light intensity in a 16-hour light/8-hour dark period at 22°C/18°C and 70% relative humidity for 14 days. 14-d-old seedlings were incubated under control conditions or at 37°C for 4 hours, followed by a 4-hour recovery phase under control conditions. Following the recovery phase, samples were flash frozen into liquid nitrogen (each biological replicate corresponds to 6 randomly chosen seedlings of the same genotype from identically treated plates) until further use.

*P. patens* (Gransden 2004 strain) was grown as described previously^65,66^. In short, they were cultivated on BCD-AT media (MgSO4·7H2O 250 mg/L, KH2PO4 250 mg/L, KNO3 1010 mg/L, ammonium tartrate 920 mg/L, FeSO4·7H2O 12.5 mg/L, CaCl2·H2O 147 mg/L, trace elements 1 ml/L [H3BO3 61 mg/L, {NH4}6Mo7O24·4H2O 38 mg/L, CuSO4·5H2O 6 mg/L, CoCl2·6H2O 5.1 mg/L, ZnCl2 4.1 mg/L, MnCl2·4H2O 4.1 mg/L; use MiliQ-water], pH 6.5 adjusted with KOH, agar 8 g/L) overlaid with cellophane discs (AA Packaging Ltd special order) when needed. Plants were grown at 22°C with 55 µE m^-2^ s^-1^ in a 16/8-hour light/dark cycle. For affinity purification, *P. patens* plants were grown on solid BCD media (same composition as BCD-AT except for ammonium tartrate) overlaid with cellophane for 13 days and moved to liquid BCD media to acclimate for 24 hours. 14-d-old plants were transferred to 37°C under normal light intensity for 4 hours, followed by a 2-hour recovery phase in fresh liquid BCD media with or without 50 µM CB5083 under control conditions.

*N. benthamiana* seeds were sown on water-saturated peat-based substrate (Klasmann Substrat 2, Klasmann-Deilmann GmbH, Gerste, Germany) containing both fertilizer (Wuxal). A solution containing *Bacillus thuringiensis* was applied as a preventive control against fungus gnat (Gnatrol, Valent BioSciences, Livertyville, Illinois, United States). Plants were grown in a 8/16-hour light/dark cycle under white light with 142 µmol m^-2^ s^-1^. For affinity purification, *N. benthamiana* plants were grown for 21 days prior to *Agrobacterium*-mediated transformation. For HS treatment, plants were transported to a temperature-controlled cabinet (Percival Scientific Series 101) on the third day after transformation and treated for 5 hours at 37°C, followed by a 3-hour recovery phase under control conditions.

### Affinity purification coupled to mass spectrometry (AP-MS) of *C. reinhardtii* samples

About 3-4 grams of harvested algal material was in an equal weight-to-volume ratio (g to ml) of 2XIP Buffer (100 mM HEPES pH 6.8, 100 mM KOAc, 4 mM Mg(OAc)2, 2 mM CaCl2, 400 mM sorbitol, Protease Inhibitor Cocktail tablet, Phosphatase Inhibitor Cocktail tablet). Samples were homogenized by performing freeze-thaw cycles, mixed with an equal volume of 1X IP buffer supplemented with 2% Digitonin (Sigma Aldrich) and allow to solubilize on a carousel for 1 hour at 4°C. Lysates were cleared by centrifugation at 16,000 x g for 15 min at 4°C twice. Protein concentration was measured using Bradford protein assay (Sigma). Normalized extracts were spiked with AIM *wt* or AIM *mut* peptides dissolved in 1XIP buffer at a final concentration of 100μM and incubated with 30μl Anti-Flag M2 Magnetic Beads (Sigma Aldrich) ((previously washed and resuspended in 1X IP buffer) for 3 hours at 4°C. Pellets were washed 5 times with detergent-free 1XIP buffer and boiled for 5 min at 95°C prior to immunoblotting with the respective antibodies.

### Affinity purification coupled to mass spectrometry (AP-MS) of *M. polymorpha* samples

About 1-2 grams of plant material were harvested and homogenized using liquid nitrogen and dissolved in lysis buffer (25 mM Tris-HCl pH 7.5, 150 mM NaCl, 10% glycerol, 1 mM EDTA, 0.25% Nonidet P-40, 2% PVPP, 1 mM DTT, Protease Inhibitor Cocktail tablet). Plant lysates were cleared by centrifugation at 16,000 x g for 15 min at 4°C twice. Protein concentration was measured using Bradford protein assay (Sigma). Beads were equilibrated with pre-chilled IP buffer (25 mM Tris-HCl pH 7.5, 150 mM NaCl, 10% glycerol, 1 mM EDTA, 0.25% Nonidet P-40, 1 mM DTT). Normalized plant extracts were spiked with AIM *wt* or AIM *mut* peptides dissolved in IP buffer at a final concentration of 100μM and incubated with 25μl GFP-Trap Magnetic Agarose beads (ChromoTek) for 1 hour at 4°C. Pellets were washed for 5 times with detergent-free IP buffer and boiled for 5 min at 95°C prior to immunoblotting with the respective antibodies.

### Affinity purification coupled to mass spectrometry (AP-MS) of *A. thaliana* samples

For purification of mCherry, mCherry-AtATG8A and mCherry-AtATG8E, 7-d-old seedlings were incubated as described before and utilized as initial material. Tissue weight was determined and ground in GTEN buffer (10% glycerol, 50 mM Tris-HCl pH 7.5, 150 mM NaCl, 1 mM EDTA, 0.2% Nonidet P-40, 0.5 mM DTT, Protease Inhibitor Cocktail tablet) in a 3:1 v/w ratio. Each sample was first ground to a fine powder with a mortar and pestle in liquid nitrogen. The pulverized tissue in the mortar was then mixed with chilled buffer to form a fine paste. The crude extract was allowed to thaw in the mortar on ice for approximately 15 minutes while other samples were processed similarly. All samples were collected in 15 ml falcon tubes and placed on a carousel for 25 minutes at 4°C. Subsequently, the samples were centrifuged twice in a swinging-bucket rotor for 15 minutes at maximum speed and cleared crude extract was transferred into fresh tubes and kept on ice during total protein quantification. Quantification was performed using the Amidoblack method^78^. For each immunoprecipitation reaction, an equal volume of crude extract was added to LoBind Eppendorf tubes and AIM *wt* or AIM *mut* peptides were added to the protein extracts at a final concentration of 100 µM. RFP-Trap magnetic agarose (Chromotek) bead slurry was equilibrated in 1 ml GTEN buffer on a carousel at 4°C. The volume of slurry was set at 30 µl per reaction and the total volume of slurry for a given experiment was equilibrated in a single tube. Tubes were immobilized on a magnetic stand and the fixed volume of equilibrated beads were added to each tube. The binding reaction was carried out on a carousel (12 rpm) for 1 hour at 4°C. Each immune reaction was then briefly centrifuged in a table-top centrifuge and immobilized on a magnetic stand placed on ice. Crude extract was removed by pipetting and replaced with 0.9 ml GTEN buffer. All tubes were inverted approximately 6 times by hand, spun again in table-top centrifuge, and immobilized. This bead washing procedure was repeated three times with GTEN buffer. Each reaction was further washed in 10% glycerol, 50 mM Tris-HCl pH 7.5, 150 mM NaCl, 1 mM EDTA buffer (without detergent, DTT, or protease inhibitor) three more times. The first wash was done by inverting the tubes by hand, while for the second and third washes, the tubes were rotated 360° on the magnetic rack. Beads were resuspended in 200 µl of the same buffer and transferred to fresh LoBind Eppendorf tubes. 20 µl (1/10th of beads) was removed, placed in new tubes and denatured in 10 µl 2X Laemmli for quality checking each reaction by western blot. The remaining sample was drained of excess buffer and flash-frozen in liquid nitrogen before on-bead trypsin digestion.

### Affinity purification coupled to mass spectrometry (AP-MS) of *P. patens* samples

About 3-4 grams of plant material were harvested and homogenized using liquid nitrogen and dissolved in lysis buffer (50 mM Tris-HCl pH 7.5, 150 mM NaCl, 10% glycerol, 1 mM EDTA, 0.2% Nonidet P-40, 2% PVPP, 1 mM DTT, Protease Inhibitor Cocktail tablet). Plant lysates were cleared by centrifugation at 16,000 x g for 15 min at 4°C twice. Protein concentration was measured using Bradford protein assay (Sigma). Beads were equilibrated with pre-chilled IP buffer (25 mM Tris-HCl pH 7.5, 150 mM NaCl, 10% glycerol, 1 mM EDTA, 0.25% Nonidet P-40, 1 mM DTT). Normalized plant extracts were spiked with AIM *wt* or AIM *mut* peptides dissolved in IP buffer at a final concentration of 100μM and incubated with 25μl GFP-Trap Magnetic Agarose beads (ChromoTek) for 1 hour at 4°C. Pellets were washed for 5 times with detergent-free IP buffer and boiled for 5 min at 95°C prior to immunoblotting with the respective antibodies.

### Affinity purification coupled to mass spectrometry (AP-MS) of *N. benthamiana* samples

About 1-2 grams of plant material were harvested and homogenized using liquid nitrogen and dissolved in lysis buffer (25 mM Tris-HCl pH 7.5, 150 mM NaCl, 10% glycerol, 1 mM EDTA, 0.25% Nonidet P-40, 2% PVPP, 1 mM DTT, Protease Inhibitor Cocktail tablet). Plant lysates were cleared by centrifugation at 16,000 x g for 15 min at 4°C twice. Protein concentration was measured using Bradford protein assay (Sigma). Beads were equilibrated with pre-chilled IP buffer (25 mM Tris-HCl pH 7.5, 150 mM NaCl, 10% glycerol, 1 mM EDTA, 0.25% Nonidet P-40, 1 mM DTT). Normalized plant extracts were spiked with AIM *wt* or AIM *mut* peptides dissolved in IP buffer at a final concentration of 100μM and incubated with 25μl GFP-Trap Magnetic Agarose beads (ChromoTek) for 1 hour at 4°C. Pellets were washed for 5 times with detergent-free buffer and boiled for 5 min at 95°C prior to immunoblotting with the respective antibodies.

### Affinity purification of biotinylated proteins

For the affinity purification with streptavidin beads, plant material was ground mechanically with a pestle and mortar, resuspended in equal volume of the extraction buffer (50 mM Tris pH 7.5, 150 mM NaCl, 0.1% SDS, 1% Triton-X-100, 0.5% Sodium deoxycholate, 1 mM EGTA, 1 mM DTT, 1x Protease Inhibitor Cocktail tablet, 1 mM PMSF, 10 µM Bortezomib) and incubated on a rotor wheel at 4°C for 10 minutes. 1 μl/ml of in-house Benzonase (0,5 mg/ml) was added to digest cell walls and DNA/RNA and the suspension was incubated on the rotor wheel at 4°C for another 15 minutes. The extracts were then distributed into 1.5 ml reaction tubes and sonicated in an ice bath four times for 30 seconds on a high setting using a Bioruptor UCD-200 (Diagenode) with 1.5 minutes breaks. The suspension was centrifuged for 15 minutes at 4°C and 15,000 x g to remove cell debris and the clear supernatant was applied to a PD-10 desalting column (Cytiva) to remove excess free biotin using the gravity protocol according to the manufacturer’s instructions. Briefly, the column was equilibrated with 4 x 5 ml of ice-cold equilibration buffer (extraction buffer without Protease Inhibitor, Bortezomib and PMSF), and 1 x 5 ml extraction buffer. 2.5 ml of the protein extract were loaded, and proteins were eluted with 3.5 ml extraction buffer. The protein concentration of the protein extract was then measured by the Amidoblack method^78^. Bead slurry corresponding to a 1:40 (bead slurry: µg of protein) ratio was prewashed with the extraction buffer and added to the normalized protein extract. The samples were incubated on a rotor wheel at 4°C for 1 hour. After the incubation, the beads were separated from the protein extract on a magnetic rack and washed as described in Branon et *al.*^79^ with 1 ml each of the following solutions: 2x with cold extraction buffer, 1x with cold 1 M KCl, 1x with cold 100 mM Na2CO3, 1x with 2M Urea in 10 mM Tris pH 8.0, 2x with cold extraction buffer without the protease inhibitors and 4x with pre-chilled wash buffer (50 mM Tris pH 7.5, 150 mM NaCl, 1 mM EGTA). Washed beads were transferred into a new LoBind tube (Axygen). 10% of the beads were boiled in 10 μl 4x Laemmli buffer (4% SDS, 20% glycerol, 10% 2-mercaptoethanol, 0.004% bromophenol blue and 0.125 M Tris HCl, pH 6.8) at 95°C for 5 minutes for immunoblots. The rest of the beads were spun down to remove the remaining wash buffer and stored at -20°C until further processing.

### *In vivo* co-immunoprecipitation (CoIP)

For the enrichment of mCherry-AtATG8E, 7-d-old seedlings were incubated as described before and utilized as initial material. Between 150-250 mg of frozen tissue per sample was first ground to a fine powder in a 1.5 ml Safe-Lock Eppendorf tubes with glass beads (Carl Roth; 1.7-2.1 mm) using a Silamat S6 (Ivoclar Vivadent). The pulverized tissue in the Eppendorf was then mixed and homogenized with chilled GTEN buffer (10% glycerol, 50 mM Tris-HCl pH 7.5, 150 mM NaCl, 1 mM EDTA, 0.2% Nonidet P-40, 0.5 mM DTT, Protease Inhibitor Cocktail tablet) in a 3:1 v/w ratio. Tubes were placed at 4°C on a carousel (14 rpm) to allow the lysis to proceed for approximately 30 minutes. Subsequently, the samples were centrifuged twice in a microcentrifuge at 4°C, maximum speed for 15 minutes and cleared crude extract was transferred into fresh tubes and kept on ice. For each immunoprecipitation reaction, an equal volume of crude extract was added to LoBind Eppendorf tubes. Beads were equilibrated in GTEN Buffer, and normalised plant extracts were incubated with 30 µl RFP-Trap magnetic agarose (Chromotek) bead per reaction. The binding reaction was carried out on a carousel (12 rpm) for 1 hour at 4°C. Each immunoprecipitation reaction was then briefly centrifuged in a tabletop centrifuge and immobilized on a magnetic stand placed on ice. Crude extract was removed by pipetting and replaced with 0.9 ml GTEN buffer. All tubes were inverted approximately 6 times by hand, spun again in tabletop centrifuge, and immobilized. This bead washing procedure was repeated three times with GTEN buffer. Purified complexes were eluted and denatured in 20 µl 2X Laemmli Buffer for 10 minutes at 70°C prior to Western blotting.

### Sample preparation for confocal microscopy

For *A. thaliana* confocal microscopy, whole seedlings were mounted on slides with water and the roots were covered with coverslips. The epidermal cells in the root transition zone were imaged for co-localization assessment, and those in the root elongation zone were imaged for autophagic flux analysis.

For *M. polymorpha* confocal microscopy, 2-d-old thalli were placed on a microscope slide with water and covered with a coverslip. The meristematic region was used for image acquisition.

### Confocal microscopy

All *A. thaliana and M. polymorpha* images were acquired by an upright point laser scanning confocal microscope ZEISS LSM800 Axio Imager.Z2 (Carl Zeiss) equipped with high-sensitive GaAsP detectors (Gallium Arsenide), a LD C-Apochromat 40X objective lens (numerical aperture 1.1, water immersion), and ZEN software (blue edition, Carl Zeiss). GFP fluorescence was excited at 488 nm and detected between 410 and 546 nm. mScarlet fluorescence was excited at 561 nm and detected between 571 and 617 nm. mCherry fluorescence was excited at 587 nm and detected between 565-617. For each experiment, all replicate images were acquired using identical parameters. Confocal images were processed with Fiji (version 1.52, Fiji) and exported as .tiff for panel assembly.

### Image processing and quantification

For co-localization and autophagic flux experiments, single snaps were used for quantification. For co-localization experiments, Mander’s co-localization analyses^80^ were performed via Fiji software (version: 1.52 and later 2.14.0/1.54f). M1 and M2 Mander’s coefficient values were calculated via the JACoP plugin. Thresholding for each Channel were adjusted for each snap image according to the puncta signal in original confocal images via JACoP plugin settings.

For the total number of puncta quantification, images were cropped to select the root area. Brightness and contrast for each Channel was homogenized throughout all the images. Threshold was adjusted for each snap image according to the puncta signal in original confocal images and particles were counted by Analyze Particles function with 0.1-9 micron2 in size and 0.00-1.00 circularity values. Number of puncta per 10,000 µm^2^ was normalized by multiplying the number of puncta by 10,000 µm^2^, divided by crop area.

For vacuolar vs cytoplasmic puncta quantification, images were cropped to select the root area. Brightness and contrast for each Channel was homogenized throughout all the images. Cytoplasmic puncta were marked by the “multi-point” function and Region of Interest (ROI) were saved for the selection of cytoplasmic puncta for each of the images. Threshold was adjusted for each snap image according to the puncta signal in original confocal images and particles were counted by “Analyze Particles” function with 0.1-9 micron2 in size and 0.00-1.00 circularity values, images with the composite ROIs were saved for the identified particles. ROIs for cytoplasmic puncta were added to the image with the composite ROIs for the total puncta, to differentiate between the cytoplasmic and vacuolar puncta. Vacuolar puncta were quantified manually by the “multi-point” function. Number of cytoplasmic puncta was confirmed by the subtraction of the vacuolar puncta values to the total number of puncta. Total, cytoplasmic and vacuolar number of puncta per 10,000 µm^2^ were normalized by multiplying the respective puncta numbers by 10,000 µm^2^, divided by crop area. Ratios were quantified by dividing the normalized number of either vacuolar or cytoplasmic puncta by the total number of normalized puncta.

### Transmission electron microscopy (TEM)

High-pressure freezing, freeze-substitution, low-temperature embedding, and ultramicrotomy and transmission electron microscopy imaging were carried out according to a protocol described previously^81^. Briefly, after the respective plant treatments, the roots of 5-d-old seedlings were cryofixed with an HPM100 high-pressure freezer (Leica Microsystems, Austria) and incubated in freeze-substitution medium (anhydrous acetone with 0.25% glutaraldehyde and 0.1% uranyl acetate) for two days at −80°C. After raising the incubation temperature to −50°C (1°C/h), the freeze-substitution medium was washed with anhydrous acetone and the samples were embedded in HM20 resin (Ted Pella, USA) over two days at −50°C. The resin was cured by UV illumination (24 h) at −50°C. An AFS2 machine (Leica Microsystems, Austria) was used for freeze-substitution, resin embedding and polymerization.

For double-immunogold labelling of transgenic *A. thaliana* plants co-expressing AtCESAR2-GFP and mCherry-AtATG8E, α:GFP (Chicken polyclonal antibody; ab13970) and α:mCherry (Rabbit polyclonal antibody; ab167453) were diluted 1:80 and 1:40, respectively. The primary antibodies were visualized with anti-rabbit IgG (10nm; SKU.25109, Electron Microscopy Sciences) and anti-chicken (15 nm; SKU.25591, Electron Microscopy Sciences) secondary antibodies conjugated with gold particles. Preparation of thin sections (90 nm) and immunogold labelling were performed as previously described^81,82^. After post-staining with uranyl acetate and Reynolds lead citrate solutions, the sections were examined with a transmission electron microscope (Hitachi H-7650) operated at 80 kV.

### Plant phenotypic assays

For carbon starvation, approximately 30 seeds were ethanol-sterilized and then cultivated in 12-well plates containing liquid ½ MS salts (Duchefa) with 1% sucrose medium. After stratification, the plates were placed under a 16-hour light/8-hour dark cycle with a light intensity of 50 µmol m^-2^ s^-1^, with gentle constant shaking. For carbon starvation, the medium was discarded after 9 days and replaced with either a control medium ½ MS, MES, 1% sucrose medium (+C) or a carbon-deficient (-C) ½ MS, MES medium. The seedlings were rinsed twice with their respective medium and the plates with carbon-deficient medium (-C) were wrapped in aluminium foil to prevent photosynthesis. Seedlings were grown in their respective media for 6 more days before imaging.

For nitrogen starvation, approximately 30 seeds were ethanol-sterilized and grown in 12-well plates containing liquid ½ MS salts (Caisson Labs) with 1% sucrose medium. Following stratification, plates were kept in a 16-hour light/8-hour dark photoperiod with 50 µmol m^-2^ s^-1^ light intensity under constant gentle shaking. After 9 days, the medium was removed and replaced with either liquid ½ MS salts (Caisson Labs) with 1% sucrose medium (+N) or nitrogen-deficient (-N) ½ MS medium (Murashige and Skoog salt without nitrogen 575 [Caisson Labs] + Gamborg B5 vitamin mixture [Duchefa] supplemented with 0.5 g/L MES and 1% sucrose, pH 5.7). Seedlings were rinsed twice in their respective medium and put back in the same culture conditions for 6 days before pictures were taken.

For heat stress (HS) phenotypic assays, 7 x 7 x 6.5 cm (width, depth, height) pots were filled with 58 grams of peat-based substrate (Klasmann Substrat 2, Klasmann-Deilmann GmbH, Geeste, DE) and left to soak water for 24 hours. *Bacillus thuringiensis* solution was used for fungus gnats. Seeds of the four indicated genotypes (Figure 5), were vapor-phase sterilized (90% sodium hypochlorite and 10% HCl), and around 15 seeds were sown per pot. Genotypes and treatment positions were randomized. Seeds were stratified in the dark at 4°C for 24 hours. Seeds were left to grow for 5 days in a 8-hour light/16-hour dark photoperiod (short-day) with a light intensity of 165 µmol m^-2^ s^-1^ (PPFD), using the following LED channels: 400 nm, white 4500K, 660 nm, and 730 nm, as specified above. 5-d-old extra-seedlings were removed from pots and a single seedling per pot was left to grow for additional 16 days. 21-d-old plants were either incubated under control conditions (Control) or at 37°C (HS) for 3 consecutive days with no watering under an 8-hour light/16-hour dark photoperiod and were subsequently allowed to recover under control conditions for additional 18 days under the same pre-treatment growth conditions. Pictures were taken 18 days post-treatment. For rosette area measurement, individual pot images were cropped in 1720 x 1720 pixels and converted to 8-bit format. Pixel units were converted to centimetres (930 pixels = 7 cm) for all images. The area of each rosette was selected and adjusted manually using the Threshold command. The Paintbrush tool was used to remove regions corresponding to the pot borders and regions not corresponding to the rosette area. The rosette area was then measured with the Analyze Particles command in Fiji with a value of 0 for Circularity and a range of 20-Infinity for size. The values corresponding to the rosette areas were used for the statistical analysis.

For Bortezomib phenotypic assays, 40 seeds/genotype for the four different genotypes specified in Figure 5 were sown in single line arrays on solid ½ MS medium (Duchefa) with 1% sucrose and 1% agar plates supplemented with either 3.75 µM Bortezomib (Santa Cruz; CAS 179324-69-7; dissolved in DMSO) or an equivalent volume of DMSO. Genotypes were sown in random positions within the plates. Seeds were stratified in the dark at 4°C for 72 hours. Subsequently, seeds were grown in horizontally oriented plates under 50 µmol m^-2^ s^-1^ LED light intensity in a 16-hour light/8-hour dark photoperiod at 21°C for 18 days prior to imaging. 18-d-old resistant seedlings were photographed and counted. The number of resistant seedlings was subtracted to the total amount of seedlings sown to obtain the number of sensitive seedlings. Green seedlings larger than 0.3 cm of width were considered resistant.

### Proteasome activity measurements

Proteasome activity measurements upon heat stress in Arabidopsis were performed according to the protocol previously published^83^ using the Proteasome-GloTM Cell-Based Assay from Promega, Walldorf (cat. no. G1180), with only minor adaptations. Briefly, treated seedlings of the indicated genotypes were homogenized in 600 µL ice cold proteasome extraction buffer (50 mM HEPES-KOH pH 7.2, 2 mM DTT, 2 mM ATP, 0.25 M Sucrose) and incubated on ice for 10 minutes. Subsequently, samples were centrifuged at 4°C and 17,000 x g for 10 minutes. During centrifugation, proteasome assay substrates were prepared according to the previously mentioned protocol. After centrifugation, 150 µl of proteasome extract supernatants were transferred to a new microcentrifuge tube and diluted 1:1 with ice cold proteasome extraction buffer. 50 µL of each diluted extract were pipetted into three different wells of a white 96-well plate to be able to measure the three different protease activities (chymotrypsin-like, trypsin-like and caspase-like) individually. After incubating the plate for 15 minutes at room temperature in darkness, 50 µL of respective substrate solutions were added to the proteasome extracts using a multichannel pipette to start the reaction. Luminescence was measured in a microplate reader (Tecan Infinite 200 PRO) over a time course of 30 minutes with data points taken every minute. The results were analysed as previously described^83^ and data show relative luminescence units per minute (RLU min^-1^), representing respective protease activities.

### Plant pictures

All plant pictures were acquired using a Canon EOS 80D DSLR equipped with either 60 mm fixed lenses or 18-135 mm lenses, using manual setting in CR2 format. The camera was mounted on a fixed stand and a dark cloth was used as background. *A. thaliana* images were opened using Fiji, where brightness and contrast was homogenized throughout all the images, and pictures were exported in tiff format to assemble panels.

### Protein extraction and western blotting

Plant material was harvested following the respective treatments and flash-frozen in liquid nitrogen. Frozen tissue was ground to a fine powder in a 1.5 Safe-Lock Eppendorf tubes with glass beads (Carl Roth; 1,7-2,1 mm) using a Silamat S6. Total protein extraction was achieved by adding 350 µl of 2X Laemmli buffer to the ground tissue and mixing in the Silamat S6 for 20 seconds until the ground tissue and the buffer were homogenized. Samples were denatured at 70°C for 10 min and centrifuged for 5 minutes at maximum speed in a microcentrifuge. Protein quantification was performed with Bradford assay (Sigma-Aldrich) following the manufacturer’s instructions or with the Amidoblack method^78^. Briefly, 10 µl of the protein sample was diluted in 190 µl of deonized water and mixed thoroughly with 1 ml of normalized Amidoblack staining solution (90% methanol, 10% acetic acid, 0.05% Naphtol Blue Black). Samples were centrifuged at maximum speed for 10 minutes. Supernatant was discarded and the resulting pellets were washed with 1 ml of washing solution (90% ethanol and 10% acetic acid), centrifuged at maximum speed for 10 minutes and dissolved in 1 ml 0.2 N NaOH. The corresponding optical density (OD) at 630 nm was measured in a plate reader (Synergy HTX Multi-Mode Microplate Reader; BioTek) using NaOH solution as blank. Protein concentration was calculated using the formula C = (OD-b)/10a, where a and b are calibrated by Bovine Serum Albumin (BSA) standard curve of the staining solution.

For Western blotting, the indicated total protein amount was loaded on SDS-PAGE gels (4-20% Mini-PROTEAN TGX precast gel; Bio-Rad) and blotted either on PVDF Immobilon-P (Millipore) membranes using a wet transfer apparatus (Bio-Rad) in cold 25 mM Tris Base, 175 mM Glycine, 20% ethanol for 2h at 170 mA, or on nitrocellulose (Bio-Rad) using the semi-dry Trans-Blot Turbo Transfer System (Bio-Rad).

Membranes were blocked in TBST (10 mM Tris-HCl pH 7.5, 150 mM NaCl, 0.1% Tween 20) + 5% skimmed milk or TBST + 2.5% BSA at room temperature for 1 hour. This was followed by 1 hour incubation at room temperature or an overnight incubation at 4°C with the primary antibody diluted in blocking buffer. After several washes with TBST, membranes were incubated for 45 min with the respective secondary antibody diluted in TBST + 5% skimmed milk and subsequently washed before imaging. All primary and secondary antibodies are provided in the materials section.

The immune-reaction was developed using either Pierce™ ECL Western Blotting Substrate (ThermoFisher) or SuperSignal™ West Pico PLUS Chemiluminescent Substrate (ThermoFisher) and detected with either ChemiDoc Touch Imaging System (Bio-Rad) or iBright Imaging System (Invitrogen). Equal total protein loading on the membrane was corroborated by Amidoblack staining.

### Protein expression and purification for biochemical assays

All recombinant proteins were produced using *E. coli* strain Rosetta2 (DE3) pLysS. Transformed cells were grown in 2XTY media supplemented with 100μg/ml spectinomycin at 37°C to log phase (OD600 0.6–0.8), followed by induction with 300μM isopropyl β-D-1-thiogalactopyranoside (IPTG) and incubation at 18°C overnight. Cells were harvested by centrifugation and resuspended in lysis buffer of 100mM Sodium Phosphate (NaPi) pH 7.0, 300mM NaCl, 20mM imidazole supplemented with cOmplete EDTA-free protease inhibitor (Roche) and Benzonase. Cells were lysed by sonication, and lysate was clarified by centrifugation at 20,000 x g. The clarified lysate was loaded on a HisTrapFF (GE Healthcare) column pre-equilibrated with the lysis buffer. Proteins were washed with lysis buffer for 10 CV and eluted with lysis buffer containing 500mM Imidazole. The eluted fraction was buffer exchanged to 50 mM NaPi pH 7.2, 50 mM NaCl and loaded either on Cation Exchange (Resource S, Cytiva), or Anion Exchange (Resource Q, Cytiva) chromatography columns. Proteins were eluted from 5 to 55 % (v/v) of Ion exchange buffer B (50 mM NaPi pH 7.2, 1 M NaCl by NaCl) gradient in 20 CV. Finally, the proteins were separated by Size Exclusion Chromatography with HiLoad® 16/600 Superdex® 75 pg (GE HealthCare), which were previously equilibrated in 50 mM NaPi pH 7.0, 100 mM NaCl.

Proteins were concentrated using Vivaspin concentrators (5000 Da MWCO). Protein concentration was calculated from the UV absorption at 280 nm by DS-11 FX+ Spectrophotometer (DeNovix).

### Ubiquitin-linkage specificity analysis

10 nmol of Halo-tagged MpCESAR CUE domain was incubated with 100 μl of the HaloLink resin (Promega) in 500 μl of the coupling buffer (50 mM Tris pH 7.5, 150 mM NaCl, 0.05% NP-40 substitute, 0.5 mM TCEP) with constant rotation at 30 rpm for 2 hours at 4°C. The immobilized resin was centrifuged at 800 x g for 2 minutes, supernatant was removed, followed by three times washing with HALO-wash buffer (50 mM Tris pH 7.5, 250 mM NaCl, 0.2% NP-40, 0.5 mM TCEP) and resuspended in 100 μl pre-chilled HALO-pulldown buffer (50 mM Tris pH 7.5, 150 mM NaCl, 0.1% NP-40, 0.5 mM TCEP, 0.5 mg/ml BSA). The ubiquitin-linkage specificity was defined through pulldown analysis by incubating 10 μl of the coupled Halo-MpCESAR CUE resin with 30 pmol of tetra-ubiquitin (Ub4) chains of the indicated linkages (M1, K6, K11, K29, K33, K48, and K63) in 500 μl of pull-down buffer for 1 hour at 4°C. The resins were spun at 800 x g and 4°C for 2 minutes to remove unbound Ub4 chains and washed two times with 500 μl HALO wash buffer and once with 500 μl HALO-coupling buffer. Captured Ub4 chains were eluted by adding 20 μl LDS sample buffer, subjected to SDS-PAGE (4 to 12% Bis-tris gel, Life Technology) and visualized by silver staining using Pierce Silver stain kit (ThermoFischer).

### Isothermal titration calorimetry (ITC)

All experiments were carried out at 25°C in 50mM NaPi pH 7.0, 100mM NaCl buffer using the PEAQ-ITC Automated (Malvern Panalytical Ltd), unless otherwise stated. For protein–peptide interactions, the calorimetric cell was filled with MpATG8 and titrated with either AIM *wt*, AIM *mut* or UIM *wt* peptides with the concentrations indicated. A single injection of 0.4 μl of titrant (not considered) was followed by either 13 injections of 3 μl each or 18 injections of 2 μl each. Injections were made at 150 seconds intervals with a duration of either 6 seconds for 13-injections experiments or 4 seconds for 18-injections experiments and a stirring speed of 750rpm. The reference power was set to 10 μcal/s; the feedback mode was set to high. For protein-protein interactions, the calorimetric cell was filled with AtATG8A and titrated with AtRPN10^201–386^ with the indicated concentrations at 15°C^17^. A single injection of 0.4 μl of titrant (not considered) was followed by 18 injections of 2 μl each. Injections were made at 150 seconds intervals with a duration of 4 seconds and a stirring speed of 500rpm. The reference power was set to 10 μcal/s; the feedback mode was set to high. The raw titration data were integrated with NITPIC^84^, globally fitted with SEDPHAT^85^, and plotted with GUSSI^86^.

### *In vitro* pulldowns

For pulldown experiments, 5 µl of glutathione magnetic agarose beads (Pierce Glutathione Magnetic Agarose Beads, Thermo Scientific) were equilibrated by washing them twice with wash buffer (100 mM NaPi pH 7.2, 300 mM NaCl, 1 mM DTT, 0.01% (v/v) Nonidet P-40). Normalized *E. coli* clarified lysates were mixed, added to the washed beads, and incubated on an end-over-end rotator for 1 hour at 4°C. Beads were washed five times in 1 ml wash buffer. Bound proteins were eluted by adding 50 µl Laemmli buffer. Samples were analyzed by western blotting.

### TUBE pulldown assays

7-d-old *A. thaliana* seedlings treated as previously indicated were utilized as initial material. Around 0.2 grams of tissue was harvested and homogenized using liquid nitrogen and immediately dissolved in grinding buffer (25 mM Tris-HCl, pH 7.5, 150 mM NaCl, 1 mM EDTA, 10% glycerol, 0.25% Nonidet P-40, 1 mM DTT, Protease Inhibitor Cocktail tablet) by vortexing. Plant lysates were cleared by centrifugation at 16,000 x g for 15 min at 4°C. Protein concentration was measured using Bradford protein assay (Sigma-Aldrich). Halo^TM^ Magnetic Beads (Promega) were coupled to recombinant Halo-tagged Tandem Ubiquitin Binding Entities (TUBEs) overnight at 4°C and washed 5 times in GTEN buffer. Normalized plant extracts were incubated with 10 μl TUBE-coupled Halo beads for 4 hours at 4°C. Beads were washed for 5 times with GTEN buffer and boiled for 5 min at 95°C prior to immunoblotting with the respective antibodies.

### Aggregate isolation and co-fractionation assays

*A. thaliana* 7-d-old seedlings of the specified genotype (Col-0, *atg5, cesar1cesar2*) were incubated at 37°C for 4h and recovered either for 4 hours (AtCESAR1-mCherry or AtCESAR2-GFP fractionation assays) or 20h (aggregate isolation) under control conditions. Seedlings were dried and weighted followed by flash-freezing in liquid nitrogen and stored at -80°C. The seedlings were ground in a pre-chilled cryo-mill (Vibrating Mill MM 400, Retsch) for 5 cycles at 30 Hz for 30 seconds with cooling in liquid nitrogen in between cycles. Aggregate fractionation buffer (100 mM HEPES pH 7.4, 1% Triton X-100, 300 mM NaCl) supplemented with Protease Inhibitor Cocktail (Roche) and 25 mM N-ethylmaleimide was added at 3 volumes per unit mass of dried seedling weight. The lysate was filtered through Miracloth and centrifuged at 2000 x g for 5 minutes at 4°C. The clear supernatant was decanted and clarified by filtration through a 0.2 µm PVDF centrifugal filter. The protein concentration of the flowthrough (referred to as whole seedling extract) was quantified using the BCA assay (ThermoFischer Scientific). Subsequently the whole seedling extract was centrifuged at 20,000 x g for 1 hour at 4°C. The supernatant was retained as the soluble fraction and its total volume was used for later normalization steps. The pellet was washed twice by pipette resuspension in aggregate isolation buffer and centrifuged at 20,000 x g for 1 hour at 4°C. The pellet was then boiled in 2X Laemmli buffer and analyzed, together with the soluble fraction, by western blot. For mass spectrometry a sonication step was included (2 sessions on a Diagenode Bioruptor at 4°C at maximum amplitude with a 50% duty cycle for a total of 5 minutes). Subsequently the pellet was boiled and partially resolved on SDS-PAGE followed by in-gel digestion followed up by standard MS sample processing procedure.

### Sample preparation for LC-MS/MS

On-beads samples were resuspended in 50 µl of 100 mM ammonium bicarbonate supplemented with 400 ng of lysyl endopeptidase (Lys-C, Fujifilm Wako Pure Chemical Corporation) and incubated at 37°C for 4 hours on a Thermo-shaker at 1200 rpm. Supernatants were transferred to fresh tubes and reduced with 0.5 mM Tris 2-carboxyethyl phosphine hydrochloride (TCEP, Sigma) for 30 minutes at 60°C and alkylated in 3 mM methyl methanethiosulfonate (MMTS, Fluka) for 30 minutes at room temperature protected from light. Subsequently, the sample was digested with 400 ng trypsin (Trypsin 875 Gold, Promega) at 37°C overnight. The digest was acidified by addition of trifluoroacetic acid (TFA, Pierce) to 1%. A similar aliquot of each sample was analysed by LC-MS/MS.

Coomassie-stained gel samples were cut into 2-3 mm pieces, transferred to 0.6 ml tubes and washed with 200 µl of 100 mM ammonium bicarbonate for 10 minutes at room temperature with shaking. The solution was removed, and pieces were destained by 2 repeated rounds of shrinking in 200 µl of 50% acetonitrile in 50 mM ammonium bicarbonate and reswelling in 200 µl of 100 mM ammonium bicarbonate. Gel pieces were shrunk with 100 µl acetonitrile before being reduced with 100 µl of 6 mM DTT in 100 mM ammonium bicarbonate by incubation at 57°C for 30 minutes and alkylated with 100 µl of 28 mM MMTS in 100 mM ammonium bicarbonate by incubation at room temperature for 30 minutes. Wash steps were repeated as described for de-staining and gel pieces were shortly dried in a speed-vac after the final shrinking step. Gel pieces were soaked in 12.5 ng/µl Trypsin in ammonium bicarbonate for 5 minutes at 4°C. Excess solution was removed, ammonium bicarbonate was added to cover the pieces, and samples were kept overnight at 37°C. The supernatant containing tryptic peptides was transferred to a fresh tube and gel pieces were extracted by addition of 20 µl 5% v/v formic acid and sonication for 10 minutes in a cooled ultrasonic bath. This step was performed twice. All supernatants were then combined. A similar aliquot of each digest was analysed by LCMS/MS.

### NanoLC-MS/MS data acquisition and processing

The nano HPLC system (UltiMate 3000 RSLC nano system or Vanquish Neo UHPLC, Thermo Fisher Scientific) was coupled to an Orbitrap Exploris 480 mass spectrometer equipped with a Nanospray Flex ion source (Thermo Fisher Scientific).

Peptides were loaded onto a trap column (PepMap Acclaim C18, 5 mm × 300 μm ID, 5 μm particles, 100 Å pore size, Thermo Fisher Scientific) at a flow rate of 25 μl/min using 0.1% TFA as mobile phase. After loading, the trap column was switched in line with the analytical column (PepMap Acclaim C18, 500 mm × 75 μm ID, 2 μm, 100 Å, Thermo Fisher Scientific). Peptides were eluted using a flow rate of 230 nl/min, starting with the mobile phases 98% A (0.1% v/v formic acid in water) and 2% B (80% v/v acetonitrile, 0.1% v/v formic acid) and linearly increasing to 35% B over the next 120 minutes. This was followed by a steep gradient to 95%B in 5 minutes, incubation for 5 minutes and ramped down in 2 minutes to the starting conditions of 98% A and 2% B for equilibration at 30°C.

The Orbitrap Exploris 480 mass spectrometer was operated in data-dependent mode ‘Cycle Time’, performing a full scan (m/z range 350-1,200, resolution 60,000, normalized AGC target 100%, minimum intensity set to 25,000 and compensation voltages CV of - 60V and -75V), followed by MS/MS scans of the most abundant ions for a cycle time of 0.9 seconds per CV. MS/MS spectra were acquired using an isolation width of 1.2 m/z, normalized AGC target 100%, HCD collision energy of 30 and orbitrap resolution of 15,000. Precursor ions selected for fragmentation (include charge state 2-6) were excluded for 45 seconds. The monoisotopic precursor selection (MIPS) mode was set to Peptide and the exclude isotopes feature was enabled.

For peptide identification, the RAW-files were loaded into Proteome Discoverer (version 2.5.0.400, Thermo Scientific). All MS/MS spectra were searched using MSAmanda v2.0.0.16129^87^. The peptide mass tolerance was set to ± 10 ppm and fragment mass tolerance to ± 8 ppm, the maximum number of missed cleavages was set to 2, using tryptic enzymatic specificity without proline restriction. Peptide and protein identification was performed in two steps. For an initial search the RAW-files were searched against the following databases for the different species: Chlamydomonas_reinhardtii_CC4532_v6.1_Phytozome.fasta, MpTak_v6.1r2.protein.fasta, Ppatens_870_v6.1protein_noStar_crc64_annot.fasta, Arabidopsis database TAIR10 and Nicotiana NbD dataset^88^ supplemented with common contaminants and sequences of tagged proteins of interest using Iodoacetamide derivative on cysteine as a fixed modification. The result was filtered to 1% FDR on protein level using the Percolator algorithm^89^ integrated in Proteome Discoverer. A sub-database of proteins identified in this search was generated for further processing. For the second search, the RAW-files were searched against the created sub-database using the same settings as above and considering the following additional variable modifications: oxidation on methionine, glutamine to pyro-glutamate conversion at peptide N-terminal glutamine and acetylation on protein N-terminus. The localization of the post-translational modification sites within the peptides was performed with the tool ptmRS, based on the tool phosphoRS^90^. Identifications were filtered again to 1% FDR on protein and PSM level, additionally an Amanda score cut-off of at least 150 was applied. Proteins were filtered to be identified by a minimum of 2 PSMs in at least 1 sample. Protein areas have been computed in IMP-apQuant^91^ by summing up unique and razor peptides. Resulting protein areas were normalised using iBAQ^92^ and sum normalisation was applied between samples. Match-between-runs (MBR) was applied for peptides with high confident peak area that were identified by MS/MS spectra in at least one run. Proteins were filtered to be identified by a minimum of 2 PSMs in at least 1 sample and quantified proteins were filtered to contain at least 3 quantified peptide groups. Statistical significance of differentially expressed proteins was determined using limma^93^.

### Tandem Mass Tag (TMT) sample preparation, data acquisition and data processing

100 µg of peptides (in 100 µl 100 mM HEPES pH 7.6) were labelled with one separate channel of the TMTpro™ 16plex Label Reagent Set (Thermo Fischer Scientific), according to the manufacturer’s description. The labelling efficiency was determined by LC-MS. Samples were mixed in equimolar amounts and equimolarity was again evaluated by LC-MS/MS. The mixed sample was acidified to a pH below 2 with 10% TFA and was desalted using C18 cartridges (Sep-Pak Vac 1cc 50 mg, Waters). Peptides were eluted with 3 x 150 µl 80% Acetonitrile and 0.1% Formic Acid, followed by freeze-drying.

Separation of TMT-labelled peptides was achieved by strong cation exchange chromatography (SCX). The dried sample was dissolved in 70 µl of SCX buffer A (5 mM NaH2PO4, pH 2.7, 15% Acetonitrile) and 200 µg of peptides were loaded at a flow rate of 35 µl/min on a TSKgel SP-2SW column (1 mm ID x 300 mm, particle size 5 µm, TOSOH) on an UltiMate 3000 RSLC nano system (Thermo Fisher Scientific) at a flow rate of 35 µl/min. For the separation, a ternary gradient was used. Starting with 100% buffer A for 10 minutes, followed by a linear increase to 10% buffer B (5 mM NaH2PO4, pH 2.7, 1M NaCl, 15% Acetonitrile) and 50% buffer C (5 mM Na2HPO4, pH 6, 15% Acetonitrile) in 10 minutes, to 25% buffer B and 50% buffer C in 10 minutes, 50% buffer B and 50% buffer C in 5 minutes and an isocratic elution for further 15 minutes. The flow-through was collected as a single fraction, then 60 fractions were collected every minute along the gradient. Fractions with low peptide content were pooled. Acetonitrile was removed by vacuum centrifugation and the samples were acidified with 0.1% TFA and analysed by LC-MS/MS.

The nano HPLC system and settings were the same as described in AP-MS/MS section, except that the gradient was increased linearly to 40% buffer B. The Orbitrap Exploris^TM^ 480 mass spectrometer was operated in data-dependent mode, performing a full scan (m/z range 350-1200, resolution 60.000, target value 100%) at 3 different compensation voltages (CV -40, -55, -70), followed each by MS/MS scans of the most abundant ions for a cycle time of 1 second per CV. MS/MS spectra were acquired using an HCD collision energy of 34%, isolation width of 0.7 m/z, resolution of 45,000, first fixed mass 110, target value at 200%, minimum intensity of 2.5E4 and maximum injection time of 120 miliseconds. Precursor ions selected for fragmentation (include charge state 2-6) were excluded for 45 seconds. The monoisotopic precursor selection (MIPS) mode was set to Peptide, the exclude isotopes and single charge state per precursor feature were enabled.

Peptide and protein identification was performed as described in the AP-MS/MS section but using only one search step and considering the following modifications: Iodoacetamide derivative on cysteine and TMTpro-16plex tandem mass tag on peptide N-Terminus were set as fixed modification, deamidation on asparagine and glutamine, oxidation on methionine, carbamylation on lysine, TMTpro-16plex tandem mass tag on lysine as well as carbamylation on peptide N-Terminus, glutamine to pyro-glutamate conversion at peptide N-Terminus and acetylation on protein N-Terminus were set as variable modifications. Results were filtered as described for IPs including an additional minimum PSM-count of 2 across all fractions. Peptides were quantified based on Reporter Ion intensities extracted by the “Reporter Ions Quantifier”-node implemented in Proteome Discoverer. Proteins were quantified by summing unique and razor peptides and filtered for quantification by at least 3 peptides. Protein-abundance-normalization was done using sum normalization. Statistical significance of differentially expressed proteins was determined using limma^93^.

### Differential protein enrichment analysis

The total number of MS/MS fragmentation spectra was used to quantify each protein (Tables S2, S4, S5, S6, S7, S10, S11). The data matrix of Peptide-Spectrum Match (PSM) was analyzed using the R package IPinquiry4 (https://github.com/hzuber67/IPinquiry4) that calculates Log2 Fold change (LFC) and *P* values using the quasi-likelihood negative binomial generalized loglinear model implemented in the edgeR package^94^. Only proteins identified with at least 2 PSM were considered. Each genotype∼treatment was triplicated per experiment, except the empty vector control IPs which were acquired as duplicates per treatment. For analysis of AIM peptide and ATG8B^ADS^ effect in *M. polymorpha*, two or three levels of filtering were employed: Empty vector control *vs.* bait (Adjusted *P* value < 0.05 & LFC > 0), AIM *wt vs.* AIM *mut* (Adjusted *P* value < 0.05 & LFC > 0) and ATG8B^ADS^ *vs.* ATG8B WT reporter (Adjusted *P* value < 0.05 & LFC > 0). For analysis of AIM peptide effect in *P. patens*, two levels of filtering were employed: Empty vector control *vs.* bait (Adjusted *P* value < 0.01 & LFC > 0) and AIM *wt vs.* AIM *mut* (Adjusted *P* value < 0.05 & LFC > 0). For analysis of AIM peptide effect in *N. benthamiana*, two levels of filtering were employed: Empty vector control *vs.* bait (Adjusted *P* value < 0.05 & LFC > 0) and AIM *wt vs.* AIM *mut* (*P* value < 0.05 & LFC > 0). For analysis of AIM peptide effect in *C. reinhardtii*, two levels of filtering were employed: Empty vector control *vs.* bait (Adjusted *P* value < 0.01 & LFC > 0) and AIM *wt vs.* AIM *mut* (*P* value < 0.05 & LFC > 0). For analysis of AIM peptide effect in *A. thaliana* under control conditions, two levels of filtering were employed: Empty vector control *vs.* bait (Adjusted *P* value < 0.01 & LFC > 0) and AIM *wt vs.* AIM *mut* (Adjusted *P* value < 0.01 & LFC > 0), in which all control reactions were pooled for the pairwise comparison. Annotations were retrieved for each protein detected in any given experiment using TAIR bulk data retrieval gene description tool (https://www.arabidopsis.org/tools/bulk/genes/index.jsp). Overlap analysis was performed using biotools.fr online (https://www.biotools.fr/misc/venny) at the indicated threshold. Protein abundance was plotted in a heatmap as Log2 (Mean PSM+1) and normalized to the mean value per protein using R. Rows were clustered using Euclidean distance and resulting dendrograms were omitted from the figures. Metric MDS plots were obtained by computing Euclidean distance matrixes and using the cmdscale command in R.

For analysis of AIM peptide effect combined with HS and HS + CB5083 (HS+CB) treatment in *A. thaliana*, four levels of filtering were employed: Empty vector control *vs.* bait (Adjusted *P* value < 0.01 & LFC > 0 with all treatments considered), AIM *wt vs.* AIM *mut* (Adjusted *P* value < 0.01 & LFC > 0 with all treatments considered), control condition *vs.* HS (Adjusted *P* value < 0.05 & LFC > 0) and control condition *vs.* HS+CB (Adjusted *P* value < 0.05 & LFC > 0). The same thresholds were used for both mCherry-AtATG8A and mCherry-AtATG8E. Protein abundance was plotted in a heatmap as indicated above, only for proteins that passed filtering. To compare the effect of the AIM peptide with the effect of the HS or HS+CB treatment, the LFC for each pairwise comparison was plotted in x and y axis. Data points were then separated in quadrants by calculating the centroid for both axes.

The analysis of proximity labelling dataset followed a strategy similar to that of classical affinity as indicated above. To identify AIM-dependent and treatment reactive proteins, several levels of filtering were employed: mCherry-TurboID *vs.* mCherry-TurboID-AtATG8E (mixed treatments) (Adjusted *P* value < 0.01 & LFC > 0), mCherry-TurboID-AtATG8E^ADS^ (mixed HS+CB treatments) *vs.* mCherry-TurboID-AtATG8E (mixed HS+CB treatments) (Adjusted *P* value < 0.01 & LFC > 0) to identify AIM-dependent ATG8 proximal proteins, and mCherry-TurboID-AtATG8E^G118A^ (mixed HS+CB treatments) *vs.* mCherry-TurboID-AtATG8E (mixed HS+CB treatments) (Adjusted *P* value < 0.01 & LFC > 0) to identify ATG8 proximal proteins that depend on ATG8-PE adduct formation. To identify proteins that are enriched in the proximity of AtATG8E during proteotoxic stress, the following comparisons were applied: mCherry-TurboID-AtATG8E HS *vs.* control conditions (*P* value < 0.01 & LFC > 0) and mCherry-TurboID-AtATG8E HS+CB *vs.* control conditions (*P* value < 0.01 & LFC > 0).

Overlap analysis was performed using biotools.fr online (https://www.biotools.fr/misc/venny) at the indicated threshold. Protein abundance was plotted in a heatmap as Log2 (Mean PSM+1) and normalized to the mean value per protein using R. Rows were clustered using Euclidean distance and resulting dendrograms were omitted from the figures.

### Analysis of protein features

For analysis of pellet-enriched protein features, protein sequences were extracted from the TAIR10 database. The molecular weight and isoelectric point of proteins were retrieved from the TAIR10 database. The relative aromaticity was calculated as the frequency of aromatic residues (phenylalanine, tryptophan, tyrosine) per protein.

For hydrophobicity analysis, each protein sequence was divided into 5-aminoacid length windows and the average hydrophobicity as defined by the Kyle-Doolittle scale^95^ was calculated. The calculated values per protein were interpolated into a standardized length format with percentiles ranging from 1 (N-terminal) to 100 (C-terminal) to allow comparison of proteins of varying lengths on a consistent scale. Profiles were smoothened using the loess method.

### AlphaFold2 (AF2) multimer screening and model building

The structure prediction of MpCESAR interaction with ubiquitin was generated by AF2 Multimer^21,22^ as implemented by ColabFold^96^. Structural analysis and figure preparation were performed using UCSF ChimeraX^97,98^.

AF2 was used to predict interactions between MpTak_v6.1r2.protein.fasta and MpATG8 isoforms. We employed a custom script to run pairwise predictions on a local CPU and GPU cluster, using MMseqs for local Multiple Sequence Alignment (MSA) creation and ColabFold^96^ for structure prediction with 5 models per prediction and omitting structure relaxation. Predictions with an average iPTM score of > 0.5 were considered putative hits and diagnostic plots (PAE plot, pLDDT plot and sequence coverage) as well as the generated structures were manually inspected.

### Crystallization conditions

MpATG8A-AIM peptide crystals were obtained using the hanging-drop vapor diffusion method at 277 K. First, 2 μl of protein solution (11 mg/ml) was mixed with 1 μl of reservoir solution containing 25% (w/v) PEG 1500, 100 mM PCTP (sodium propionic acid, sodium cacodylate, Bis-Tris propane) pH 5.0. Crystals were cryoprotected upon addition of 20% (v/v) ethylene glycol to the reservoir solution before flash-cooling.

### Crystal structure solution and refinement

Diffraction data were collected using in-house Cu-X-ray source at 100 K. Data were processed and scaled using the XDS package^99^. A total of 5% of reflections were randomly selected for calculation of a Rfree value. The phase problem was solved by molecular replacement with PHASER^100^ using potato ATG8CL (PDB: 5L83) as start model. Phases were improved in iterative cycles of manual model building in COOT^101^ and refinement with PHENIX^102^, which converged with Rwork/Rfree values of 0.195/0.240. Model quality was validated using the tools provided in COOT and MOLPROBITY^103^ with 0% Ramachandran outliers and 100% favored. Structural analysis and figure preparation were performed using UCSF ChimeraX^97,98^.

### Statistical analysis

All statistical analysis was performed using R Statistical Software (version 4.1.2; R Foundation for Statistical Computing, Vienna, Austria) (36). Statistical significance of differences between two experimental groups was assessed with a two-tailed unpaired two-samples t-test if the two groups were normally distributed (Shapiro-Wilk test) and their variances were equal (F-test). If the groups were normally distributed but the variances were not equal a two-samples Welch t-test was performed. If the groups were not normally distributed, an unpaired two-samples Wilcoxon test with continuity correction was performed. Differences between two data sets were considered significant at *P* < 0.05 (*); *P* < 0.01 (**); *P* < 0.001 (***). *P* value > 0.05 (ns, not significant).

All quantification analyses and statistical tests were performed with GraphPad Prism 8 and 10 software. For co-localization microscopy analyses, Two-tailed Mann-Whitney test was performed to analyse the significance of the differences of M1 and M2 values. For puncta quantification, Two-tailed unpaired Student’s t-tests with Welch’s corrections were performed to analyse the significance of the differences between the number of puncta between the treatments and genotypes tested. The significance level of differences between two experimental groups was marked as *, *P* < 0.05; **, *P* < 0.01; ***, *P* < 0.001; ****, *P* < 0.0001; ns, not significant. For rosette area quantification analysis and Bortezomib normalized survival rate in Figure 5, Ordinary-one-way Anova with Tukey’s multiple comparisons test was performed to assess the differences in the normalized Rosette Area and survival rates, respectively, between genotypes. For Bortezomib normalized survival rate in Figure S22, Kruskal-Wallis test followed by a Dunn’s multiple comparisons test was performed to assess the differences in the normalized survival rates between genotypes. The significance level of differences between the different experimental groups based on the multiple comparisons are marked as Adjusted *P* value < 0.05; **, Adjusted *P* value < 0.01; ***, Adjusted *P* value < 0.001; ****, Adjusted *P* value < 0.0001; ns, not significant. For acquired thermotolerance assays, unpaired one-tailed Student’s t-test was performed to assess the differences between the different genotypes at the different ACC times. Significance level marked as *, *P* value < 0.05.

## MATERIALS

### Experimental Model/Cell lines

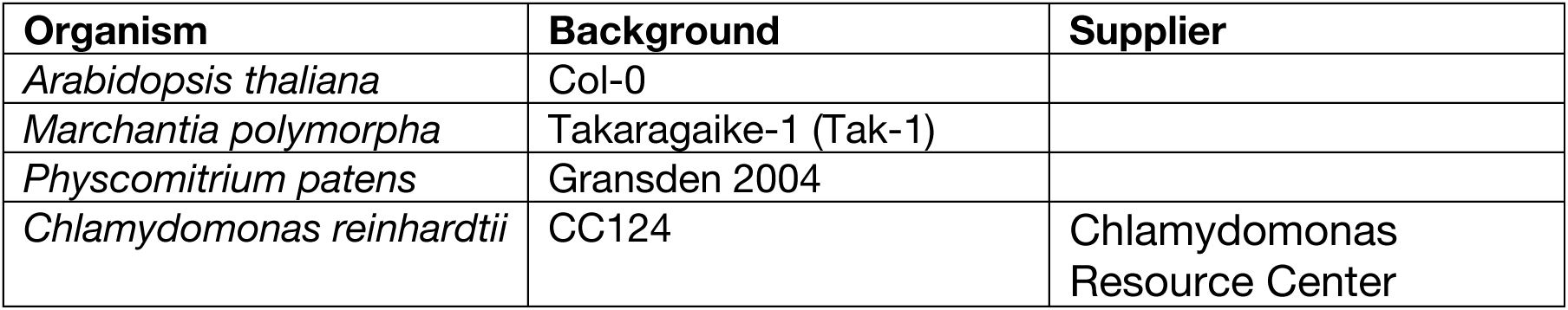

| Organism | Background | Supplier |
| --- | --- | --- |
| *Arabidopsis thaliana* | Col-0 |  |
| *Marchantia polymorpha* | Takaragaike-1 (Tak-1) |  |
| *Physcomitrium patens* | Gransden 2004 |  |
| *Chlamydomonas reinhardtii* | CC124 | Chlamydomonas Resource Center |

### Mutants

| Organism | Genotype | Gene ID | Preparation/T-DNA line | Supplier | Reference |
| --- | --- | --- | --- | --- | --- |
| *M. polymorpha* | *atg7* | Mapoly0015s0071/<br>Mp2g07850 | CRISPR |  | <sup>11</sup> |
| *A. thaliana* | *atg5-1* | AT5G17290 | SAIL_129B07 | NASC (N39993) | <sup>71</sup> |
| *A. thaliana* | *cesar1-1* | AT4G02880 | SALK_112206.2<br>3.80.x | NASC (N612206) | This study |
| *A. thaliana* | *cesar2-1* | AT1G03290 | SALK_022568 | NASC (N522568) | This study |
| *A. thaliana* | *cesar1-1*<br>/*cesar2-1* | AT4G02880<br>AT1G03290 | Crossing |  | This study |
| *A. thaliana* | *nbr1-2* | AT4G24690 | GABI_246H08 |  | <sup>72</sup> |

### Plant Lines

| Name | Accession number | Reference | Plant species |
| --- | --- | --- | --- |
| pEF1α::GFP |  | This study | *M. polymorpha* |
| pEF1α::GFP-MpATG8A | Mapoly0001s0494/<br>Mp1g21590 | This study | *M. polymorpha* |
| pEF1α::GFP-MpATG8B | Mapoly0027s0034/<br>Mp5g05930 | This study | *M. polymorpha* |
| pEF1α::GFP-MpATG8A/*atg7* | Mapoly0001s0494/<br>Mp1g21590 | This study | *M. polymorpha* |
| pEF1α::GFP-MpATG8B/*atg7* | Mapoly0027s0034/<br>Mp5g05930 | This study | *M. polymorpha* |
| pEF1α::MpCESAR-mScarlet | Mapoly0005s0053/<br>Mp1g05540 | This study | *M. polymorpha* |
| pEF1α::GFP-MpATG8A x<br>pEF1α::MpCESAR-mScarlet | Mapoly0001s0494/<br>Mp1g21590;<br>Mapoly0005s0053/<br>Mp1g05540 | This study | *M. polymorpha* |
| pEF1α::GFP-MpATG8B x<br>pEF1α::MpCESAR-mScarlet | Mapoly0027s0034/<br>Mp5g05930;<br>Mapoly0005s0053/<br>Mp1g05540 | This study | *M. polymorpha* |
| pEF1α::GFP x<br>pEF1α::MpCESAR-mScarlet | Mapoly0005s0053/<br>Mp1g05540 | This study | *M. polymorpha* |
| p2x35S::GFP |  | <sup>66</sup> | *P. patens* |
| p2x35S::GFP-PpATG8a | Pp6c17_2570 | <sup>66</sup> | *P. patens* |
| CrRS581 / pTUB2:3XFLAG-Venus-ATG8 | Cre16.g689650 | This study | *C. reinhardtii* |
| CrRS579 / pTUB2:3XFLAG-cytVenus |  | This study | *C. reinhardtii* |
| pAtATG8E::mCherry-TurboID-AtATG8E | AT2G45170 | This study | *A. thaliana* |
| pAtATG8E::mCherry-TurboID-AtATG8E<sup>ADS</sup> | AT2G45170 | This study | *A. thaliana* |
| pAtATG8E::mCherry-TurboID-AtATG8E<sup>G118A</sup> | AT2G45170 | This study | *A. thaliana* |
| pAtATG8E::mCherry-TurboID | AT2G45170 | This study | *A. thaliana* |
| pUBQ::mCherry |  | <sup>73</sup> | *A. thaliana* |
| pUBQ::mCherry-AtATG8A | AT4G21980 | <sup>16</sup> | *A. thaliana* |
| pUBQ::mCherry-AtATG8E | AT2G45170 | <sup>74</sup> | *A. thaliana* |
| pUBQ::mCherry-AtATG8E/atg5-1 | AT2G45170 (ATG8E);<br>AT5G17290 (ATG5) | <sup>16</sup> | *A. thaliana* |
| pRPS5::AtCESAR2-GFP in<br>pUBQ::mCherry | AT1G03290 | This study | *A. thaliana* |
| pRPS5::AtCESAR2-GFP in<br>pUBQ::mCherry-AtATG8E | AT1G03290<br>(CESAR2);<br>AT2G45170 (ATG8E) | This study | *A. thaliana* |
| pRPS5::AtCESAR2-GFP in<br>pUBQ::mCherry-AtATG8E/atg5-1 | AT1G03290<br>(CESAR2);<br>AT2G45170 (ATG8E);<br>AT5G17290 (ATG5) | This study | *A. thaliana* |
| pRPS5::AtCESAR2-GFP | AT1G03290 | This study | *A. thaliana* |
| pRPS5::AtCESAR1-mCherry | AT4G02880 | This study | *A. thaliana* |
| pRPS5::AtCESAR1-mCherry in<br>pRPS5::AtCESAR2-GFP | AT4G02880<br>(CESAR1);<br>AT1G03290<br>(CESAR2) | This study | *A. thaliana* |
| pUBQ::GFP-AtATG8A | AT4G21980 | <sup>16</sup> | *A. thaliana* |
| pUBQ::GFP-AtATG8A/atg5-1 | AT4G21980 (ATG8A);<br>AT5G17290 (ATG5) | This study | *A. thaliana* |
| pUBQ::GFP |  | This study | *A. thaliana* |

### Bacterial Strains

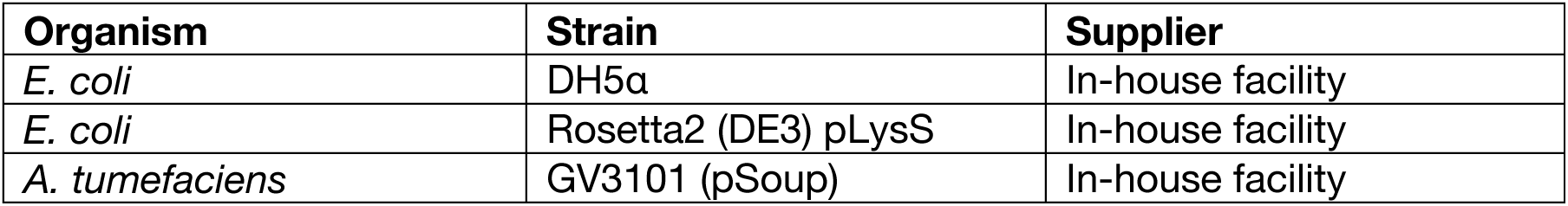

| Organism | Strain | Supplier |
| --- | --- | --- |
| *E. coli* | DH5α | In-house facility |
| *E. coli* | Rosetta2 (DE3) pLysS | In-house facility |
| *A. tumefaciens* | GV3101 (pSoup) | In-house facility |

### Oligonucleotides

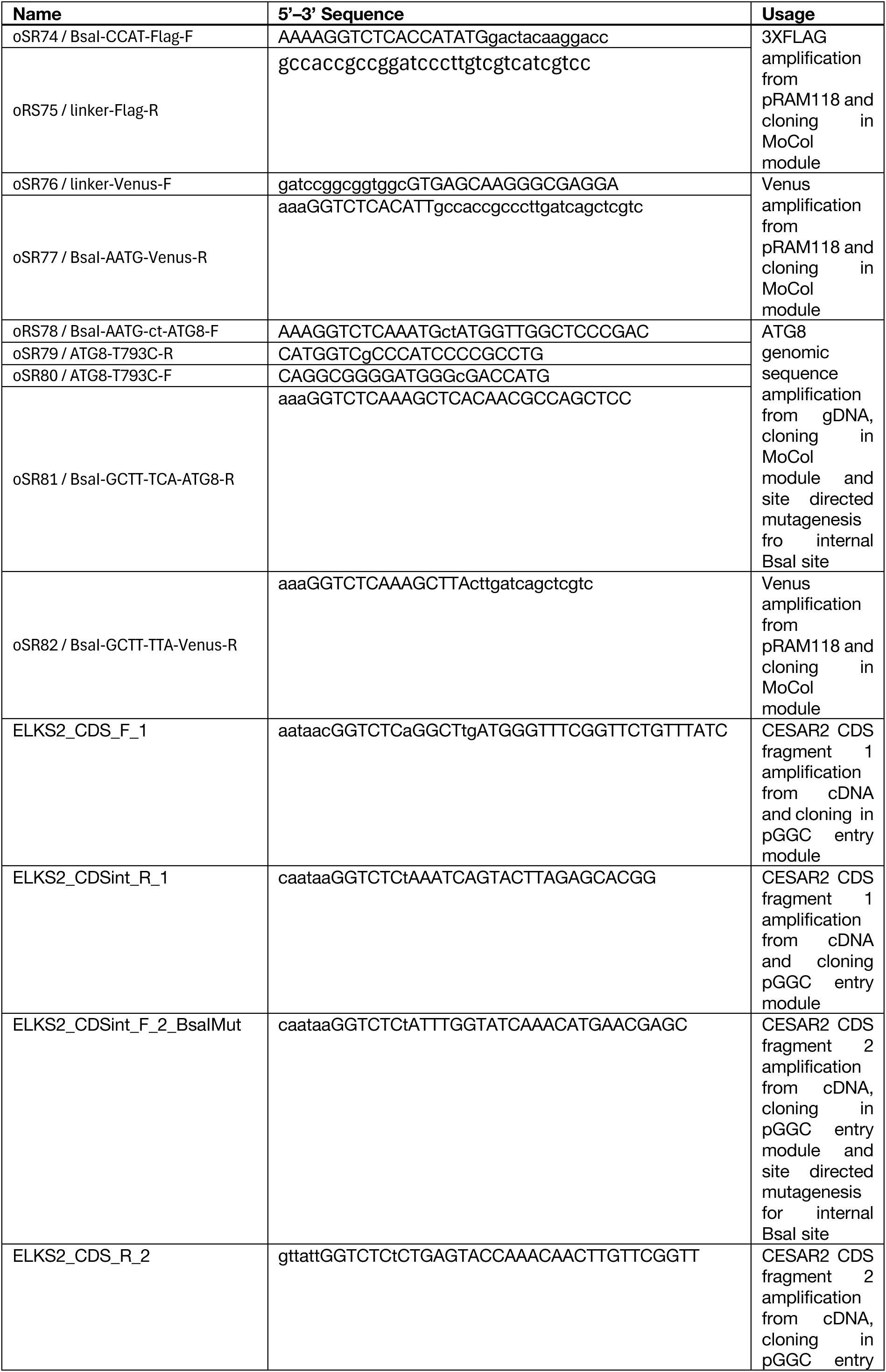
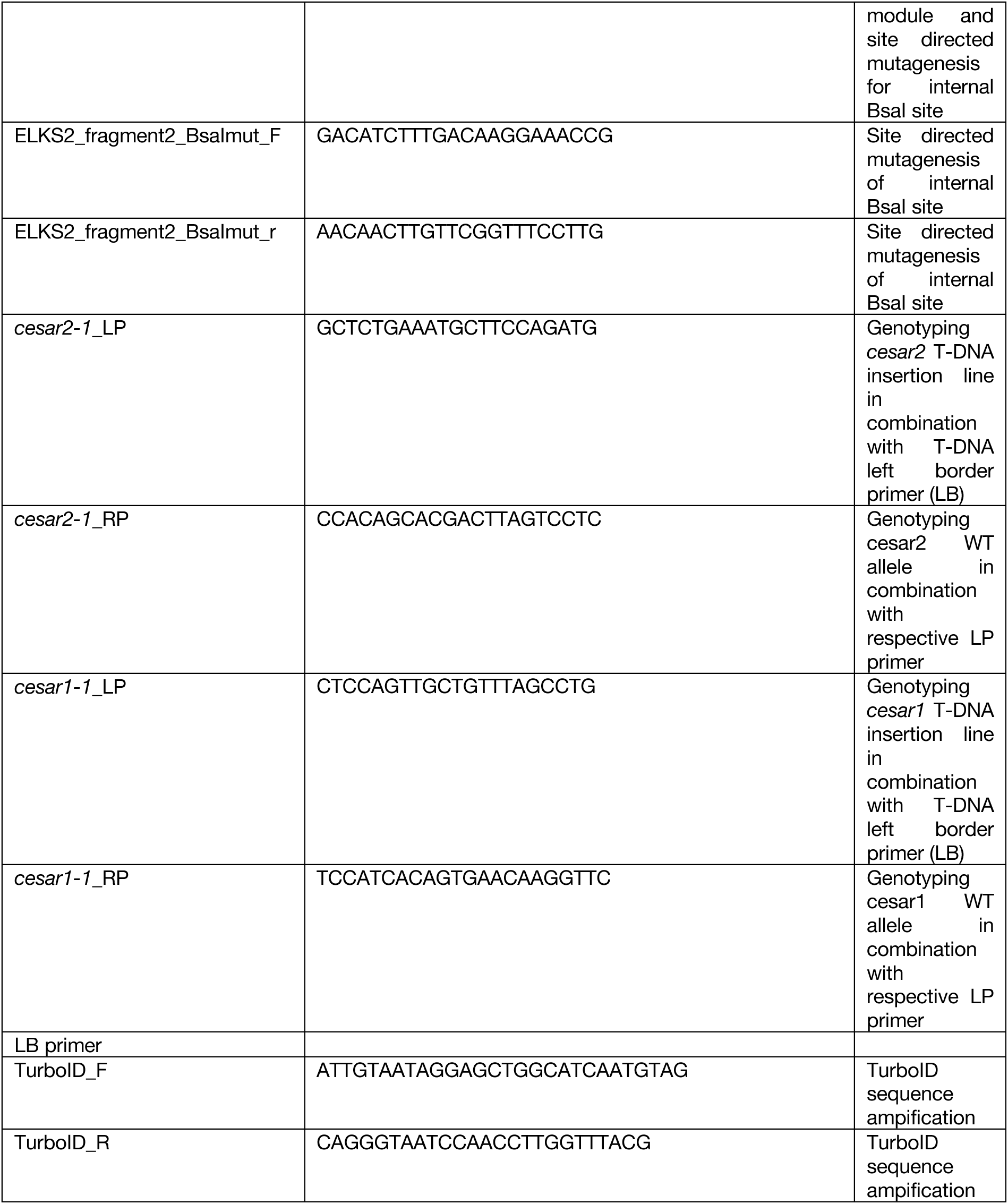

### Peptides

| Name | Source | Sequence |
| --- | --- | --- |
| AIM *wt* | Synthesized *in house* | EPLDFDWEIVLEEEM |
| AIM *mut* | Synthesized *in house* | EPLDFDAEIALEEEM |
| UIM *wt* | Synthesized *in house* | EDSALLDQAIAMSVGDVNMS |
| AIM *wt* (crystal) | Synthesized *in house* | DWEIV |

### Plasmids

| Organism | Name | Reference |
| --- | --- | --- |
| *E. coli* | E. coli destination vector (pET) | 16 |
| *E. coli* | pET-GST | 16 |
| *E. coli* | pET-MBP | 16 |
| *E. coli* | pET-GST-AtATG8A | 16 |
| *E. coli* | pET-GST-AtATG8A<sup>ADS(Y50A,L51A)</sup> | 16 |
| *E. coli* | pET-GST-AtATG8B | 16 |
| *E. coli* | pET-GST-AtATG8C | 16 |
| *E. coli* | pET-GST-AtATG8D | 16 |
| *E. coli* | pET-GST-AtATG8E | 16 |
| *E. coli* | pET-GST-AtATG8F | 16 |
| *E. coli* | pET-GST-AtATG8G | 16 |
| *E. coli* | pET-GST-AtATG8H | 16 |
| *E. coli* | pET-GST-AtATG8I | 16 |
| *E. coli* | pET-GST-MpATG8A | 16 |
| *E. coli* | pET-GST-MpATG8B | 16 |
| *E. coli* | pET-GST-MpATG8A<sup>ADS(Y52A,L53A)</sup> | 16 |
| *E. coli* | pET-GST-MpATG8B<sup>ADS(K51A)</sup> | This study |
| *E. coli* | pGEX-GST-1xUb | 104 |
| *E. coli* | pGEX-GST-2xUb | 104 |
| *E. coli* | pGEX-GST-4xUb | 104 |
| *E. coli* | pET-MBP-MpCESAR | This study |
| *E. coli* | pET-MBP-MpCESAR<sup>ΔCUE (47-987)</sup> | This study |
| *E. coli* | pET-MBP-MpCESAR<sup>I13D</sup> | This study |
| *E. coli* | pET-MBP-MpCESAR<sup>F14D</sup> | This study |
| *E. coli* | pET-MBP-MpCESAR<sup>A37D</sup> | This study |
| *E. coli* | pET-MBP-MpCESAR<sup>D38R</sup> | This study |
| *E. coli* | pET-MBP-MpCESAR<sup>L41E</sup> | This study |
| *E. coli* | pET-MBP-ApCESAR | This study |
| *E. coli* | pET-MBP-PpCESAR2 | This study |
| *E. coli* | pET-MBP-OsCESAR | This study |
| *E. coli* | pET-MBP-ZmCESAR | This study |
| *E. coli* | pET-MBP-NbCESAR | This study |
| *E. coli* | pET-MBP-AtCESAR1 | This study |
| *E. coli* | pET-MBP-AtCESAR2 | This study |
| *E. coli* | pET-MBP-AtCESAR2<sup>2A(W558A, F561A)</sup> | This study |
| *E. coli* | pET-MBP-AtCESAR2<sup>4A(L553A, L554A, W558A, F561A)</sup> | This study |
| *E. coli* | pET-MBP-MpRING1 | This study |
| *E. coli* | pET-MBP-MpRING1<sup>AIM1(W125A)</sup> | This study |
| *E. coli* | pET-MBP-MpRING1<sup>AIM2(F151A)</sup> | This study |
| *E. coli* | pET-MBP-MpRING1<sup>AIM1+2(W125A,F151A)</sup> | This study |
| *E. coli* | pET-MBP-MpRING2 | This study |
| *E. coli* | pET-MBP-MpRING2<sup>AIM1(W139A)</sup> | This study |
| *E. coli* | pET-MBP-MpRING2<sup>AIM2(Y164A)</sup> | This study |
| *E. coli* | pET-MBP-MpRING2<sup>AIM1+2(W139A,Y164A)</sup> | This study |
| *E. coli* | pET-MBP-MpYHBC | This study |
| *E. coli* | pET-MBP-MpYHBC<sup>F450A</sup> | This study |
| *E. coli* | pET-MBP-MpYHBC<sup>F450A,V453A</sup> | This study |
| *E. coli* | pET-MpVWA41-MBP | This study |
| *E. coli* | pET-MBP-MpVWA41<sup>AIM(F494A)</sup> | This study |
| *E. coli* | pET-MBP-MpVWA43 | This study |
| *E. coli* | pET-MBP-MpVWA43<sup>AIM(F485A)</sup> | This study |
| *E. coli* | pET-6xHis-3C-MpATG8A | This study |
| *E. coli* | pET-6xHis-3C-MpATG8B | This study |
| *E. coli* | pET-6xHis-3C-AtRPN10<sup>201-386</sup> | This study |
| *E. coli* | pET-6xHis-3C-AtATG8A | 105 |
| *E. coli* | pET-6xHis-TEV-Halotag-TUBE | 31 |
| *E. coli* | pET-6xHis-TEV-Halotag-MpCESAR<sup>CUE(1-46)</sup> | This study |
| *E. coli* | pGGSun | 106 |
| *M. polymorpha* | pMpGWB303-pEF1α::GFP-MpATG8A | This study |
| *M. polymorpha* | pMpGWB303-pEF1α::GFP-MpATG8B | This study |
| *M. polymorpha* | pMpGWB303-pEF1α::GFP-MpATG8B<sup>ADS(K51A)</sup> | This study |
| *M. polymorpha* | pGGSun-pEF1α::MpCESAR-mScarlet | This study |
| *M. polymorpha* | pMpGWB303-pEF1α::GFP | This study |
| *A. thaliana* | pAtATG8E::mCherry-TurboID-AtATG8E<sup>ADS</sup> | This study |
| *A. thaliana* | pAtATG8E::mCherry-TurboID-AtATG8E<sup>G118A</sup> | This study |
| *A. thaliana* | pAtATG8E::mCherry-TurboID | This study |
| *A. thaliana* | pGGSun-pRPS5::AtCESAR2_GFP | This study |
| *A. thaliana* | pGGSun-pRPS5::AtCESAR1_mCherry | This study |
| *A. thaliana* | pGGSun-pRPS5::AtCESAR2 | This study |
| *A. thaliana* | pGGSun-pRPS5::AtCESAR1 | This study |
| *C. reinhardtii* | pRAM276 / pTUB2:3XFLAG-Venus-ATG8 | This study |
| *C. reinhardtii* | pRAM275 / pTUB2:3XFLAG-cytVenus | This study |
| *P. patens* | pPTM13-p2x35S::GFP-PpATG8a | 66 |
| *S. tuberosum* | pK7WGF2-p35S::GFP-StATG8 2.2 | This study |

### Antibodies

Antibodies:
| Antibody | Host | Supplier | Cat. # | Working concentration |
| --- | --- | --- | --- | --- |
| Anti-Rabbit IgG HRP-Conjugate | Goat | Bio-Rad | 1706515 | 1:5000 |
| Anti-Mouse IgG-HRP Conjugate | Goat | Bio-Rad | 1706516 | 1:5000 |
| Anti-GFP | Mouse | Roche | 11814460001 | 1:2500 and 1:5000 |
| Anti-GFP | Rabbit | Invitrogen | A11122 | 1:5000 |
| Anti-mCherry | Rabbit | Proteintech | 26765-1-AP | 1:5000 |
| Anti-RFP 6G6 | Mouse | Chromotek | AB_2631395 | 1:5000 |
| Anti-GFP polyclonal | Chicken | Abcam | ab13970 | 1:80 |
| Anti-mCherry polyclonal | Rabbit | Abcam | ab167453 | 1:40 |
| Anti-rabbit IgG 10 nm | Goat | Electron Microscopy Sciences | SKU.25109 | 1:20 |
| Anti-chicken IgG 15 nm | Goat | Electron Microscopy Sciences | SKU.25591 | 1:40 |
| Anti-GST HRP-Conjugate | Goat | GE Healthcare | RPN1236 | 1:5000 |
| Anti-MBP | Mouse | Sigma Aldrich | M1321 | 1:5000 |
| Anti-NBR1 | Rabbit | Agrisera | AS14 2805 | 1:10000 |
| Anti-HaloTag | Mouse | Promega | G9211 | 1:5000 |
| Anti-Ubiquitin | Mouse | Santa Cruz Biotechnology | 3936 | 1:5000 |

### Inhibitors and drugs

| Name | Supplier | Cat. # |
| --- | --- | --- |
| Bortezomib | Santa Cruz Biotechnology | sc-217785 |
| CB5083 | BIOCAT GmbH | T6796-50mg-TM |
| Concanamycin A (ConA) | Santa Cruz Biotechnology | sc-202111A |
| E64d | Santa Cruz Biotechnology | sc-201280 |
| Pepstatin A | Santa Cruz Biotechnology | sc-45036 |
| Torin1 (Torin) | Santa Cruz Biotechnology | sc-396760 |

### Media and supplements

| Name | Supplier | Cat. # |
| --- | --- | --- |
| Gamborg B5 vitamin mixture 1000X | Duchefa | G0415.0250 |
| Gamborg B5 medium (microsalt mixture) | Duchefa | M0302.0025 |
| Gamborg B5 medium (including vitamins) | Duchefa | G0210.0010 |
| Gamborg B5 medium (basal salt mixture) | Duchefa | G0209.0050 |
| Murashige & Skoog vitamin mixture 1000X | Duchefa | M0409.0250 |
| Murashige & Skoog micro salt mixture | Duchefa | M0301.0050 |
| Murashige & Skoog macro salt mixture | Duchefa | M0305.0050 |
| Murashige & Skoog Basal salt mixture with MES | Duchefa | M0254.0050 |
| Murashige & Skoog without nitrogen | Caisson labs | MSP07 |
| Murashige & Skoog M51C medium | Duchefa | DU1501 |
| Casamino Acid | Calbiochem | 65072-00-6 |
| MES monohydrate | AppliChem | A1074 |
| D-Sucrose | AppliChem | 131621 |
| L-Glutamine | Sigma Aldrich | G7513 |
| Casein hydrolysate | Duchefa | C1301 |

### Cemicals and reagents

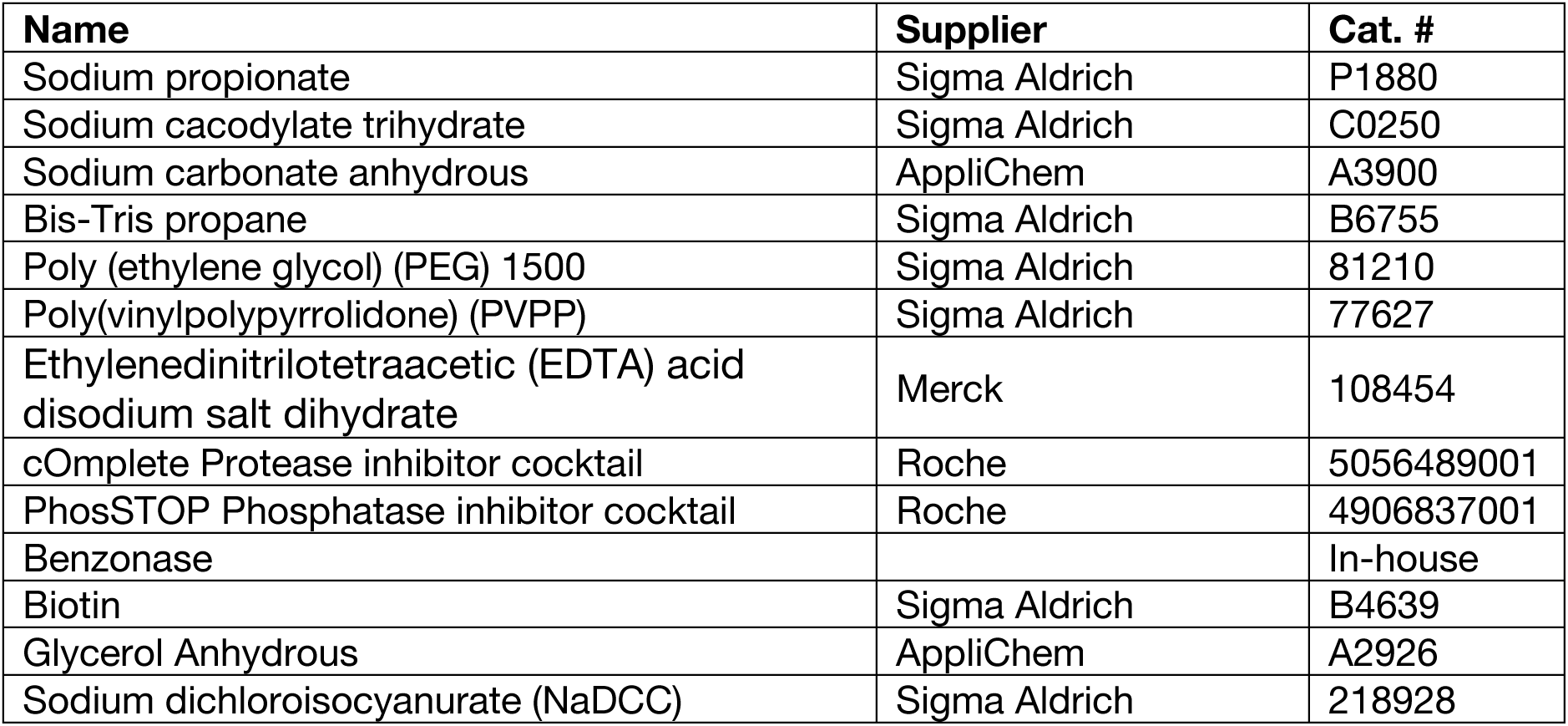
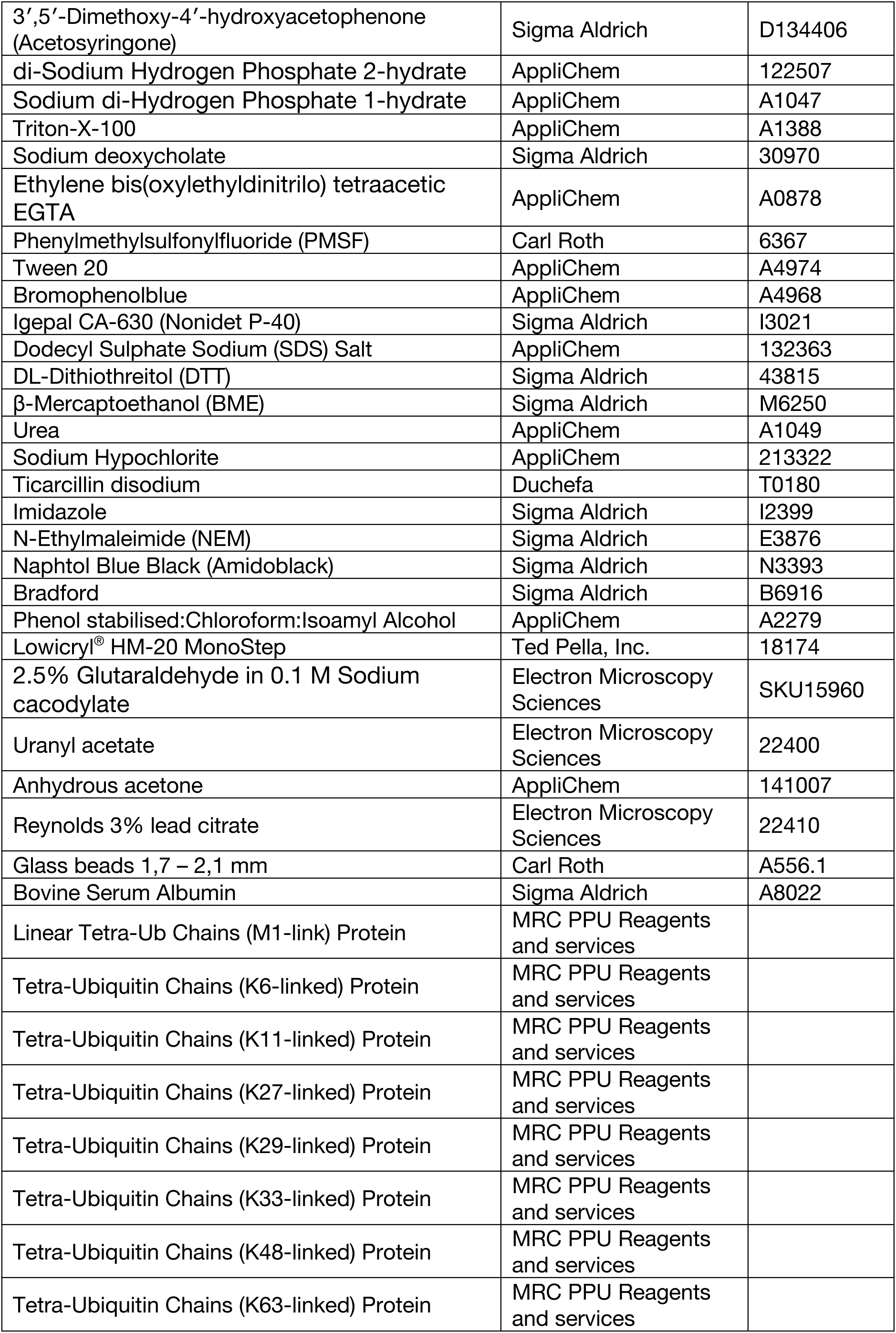

### Affinity matrices for purification and immuno-precipitation

| Name | Supplier | Cat. # |
| --- | --- | --- |
| GFP-Trap Magnetic Agarose | Chromotek | Gtma |
| RFP-Trap Magnetic Agarose | Chromotek | Rtma |
| Magne® HaloTag® Beads | Promega | G7282 |
| Pierce™ Glutathione Magnetic Agarose Beads | Thermo Scientific™ | 78601 |
| Anti-FLAG® M2 Magnetic Beads | Sigma Aldrich | M8823 |
| HisTrap FF 5 ml | GE Healthcare | 17525501 |
| Resource Q 6 ml | GE Healthcare | 17117901 |
| Resource S 6 ml | GE Healthcare | 17118001 |
| HiPrep 26/10 Desalting | GE Healthcare | 17508701 |
| HiLoad 16/600 Superdex 75 pg | GE Healthcare | 28989333 |

### Software

| Name | Company | Application |
| --- | --- | --- |
| CLC main work bench 7 | Qiagen | Cloning |
| Zen Software | Carl Zeiss | Microscopy |
| Image J (Fiji) | NIH | Image Quantification |
| Image Lab | Bio-Rad | Western Blot Analysis |
| iBright analysis software | Invitrogen | Western Blot Analysis |
| Adobe Illustrator 2022 | Adobe Inc. | Graphics editing |
| Adobe Photoshop 2023 | Adobe Inc. | Graphics editing |
| GraphPad Prism 9 | GraphPad Software | Graph plotting, Statistical analysis |
| RStudio 2021.09.2+382 "Ghost Orchid" Release; R version 4.3.0 | RStudio; The R Foundation for Statistical Computing | Graph plotting, Statistical analysis |
| UCSF ChimeraX | Resource for Biocomputing, Visualization, and Informatics (RBVI) | Structural Analysis |
| COOT | MRC Laboratory of Molecular Biology | Structural analysis |
| PHASER v1.2 | Cambridge University | X-ray data curation |
| PHENIX v1.2 | The Phenix Industrial Consortium | X-ray data curation |

### Equipment

| Name | Supplier |
| --- | --- |
| HPM100 high-pressure freezer | Leica Microsystems, Austria |
| AFS2 machine | Leica Microsystems, Austria |
| Hitachi H-7650 electron microscope | Hitachi High-Tech Group |
| Vibrating Mill MM 400 | Retsch |
| Minichiller Bioruptor UCD-200 | Diagenode |
| Silamat S6 | Ivoclar vivadent |
| Percival CU-36L4 | Percival Scientific, inc. |
| Synergy HTX Multi-Mode Microplate Reader | BioTek |
| ChemiDoc Touch Imaging System | Bio-Rad |
| iBright Imaging System | Invitrogen |

---

**Figure S1.**
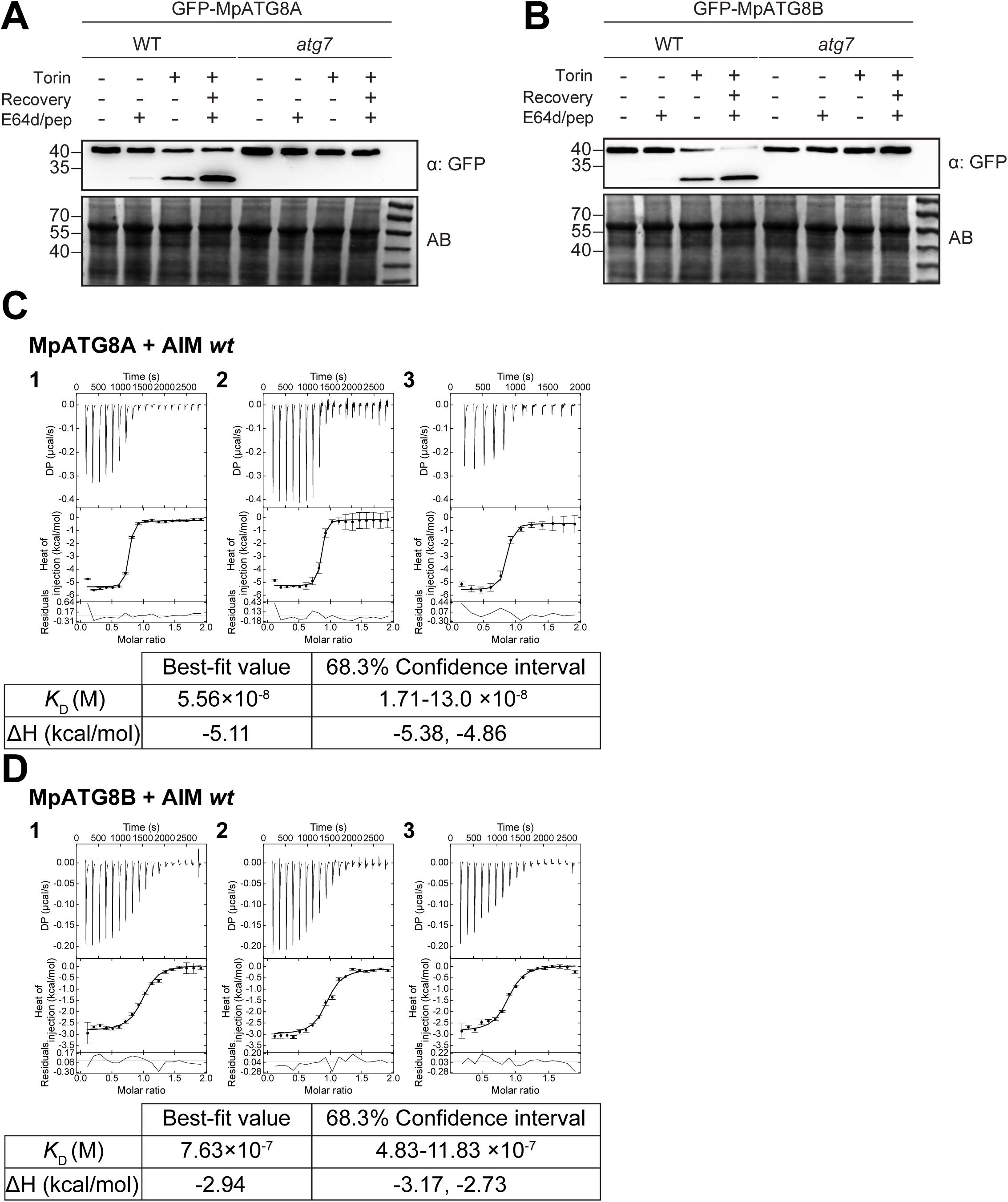
Characterization of canonical MpATG8 function. **(A-B) MpATG8 undergoes autophagic flux.** A second independent replicate of autophagic flux assays shown in Fig. 1B,C. Protein extracts were analysed by immunoblotting with anti-GFP. Total protein loading control was analysed by Amidoblack (AB) staining. **(C-D) MpATG8 binds AIM *wt*.** Isothermal titration calorimetry (ITC) experiments showing three independent titrations of AIM *wt* peptide to MpATG8A (C) or MpATG8B (D). The concentration of reactants in (C) are 25 µM (2) or 50 µM (1,3) for MpATG8A (in cell) and 250 µM (2) or 500 µM (1,3) AIM *wt* peptide (in syringe), while in (D) are 50 µM (1,2,3) for MpATG8B (in cell) and 500 µM (1,2,3) AIM *wt* peptide (in syringe). Global analysis was performed using a hetero-association model A + B. The top panels show the SVD-reconstructed thermograms, the middle panels show the isotherms as integrated heat of injection (dots) and best fit (solid line), and the bottom panels show the residuals (difference between the fit to a single hetero-association model and the experimental data; closer to zero indicates stronger agreement between data and the fit). Extracted global parameters and their 68.3% confidence interval are reported for titrations of AIM *wt* peptide with either of MpATG8 isoform. Thermograms were reconstructed with NITPIC, global analysis was done in SEDPHAT, and data visualization was plotted in GUSSI. The dissociation constant (KD) is reported in molar (M) units, while the enthalpy (ΔH) is reported in kcal/mol units. Note that replicate 1 for AIM *wt* peptide titration to MpATG8A is shown in Fig. 1E.

**Figure S2.**
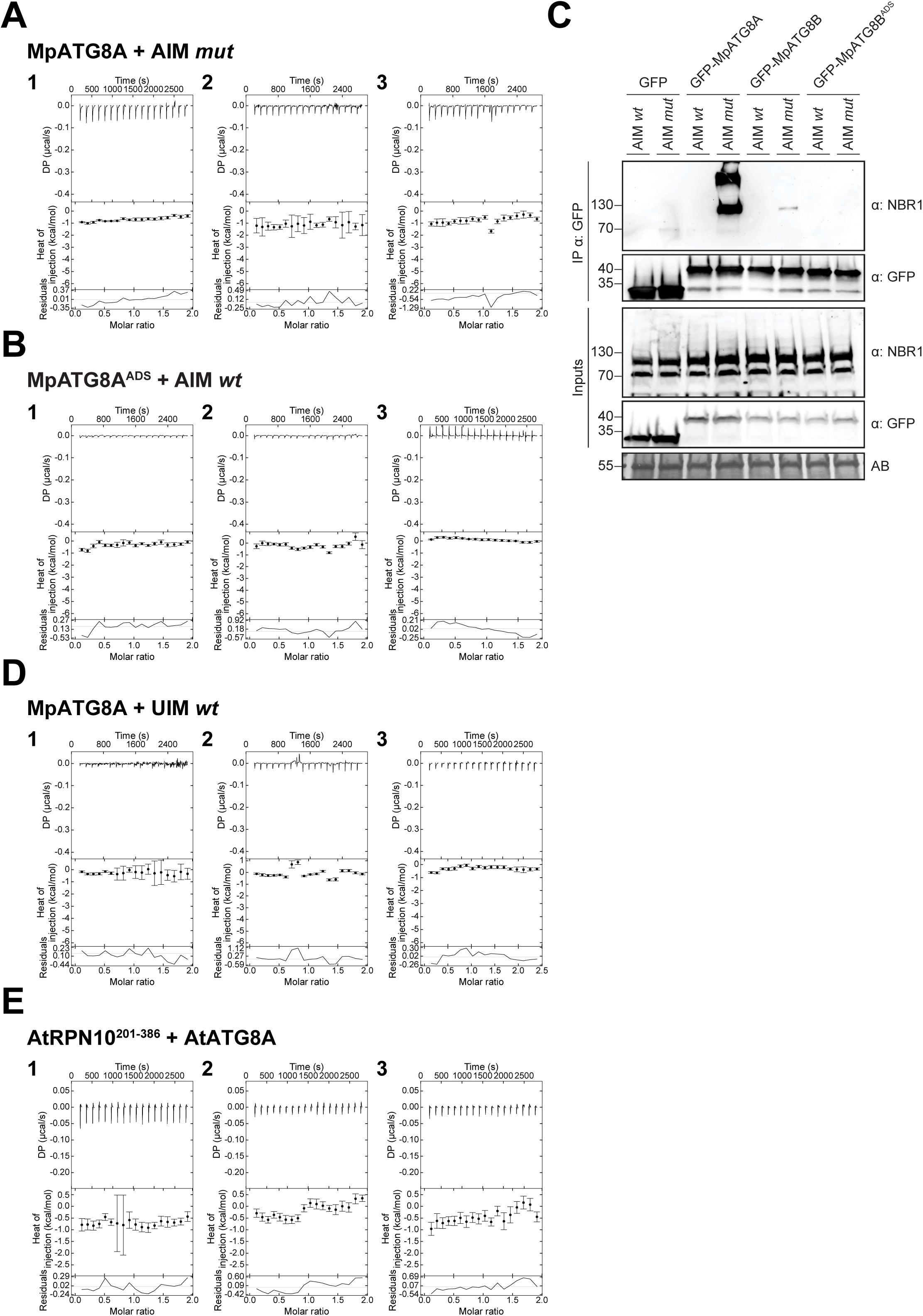
MpATG8^ADS^ mutagenesis supports canonical AIM binding. **(A) MpATG8A does not bind AIM *mut*.** Isothermal titration calorimetry (ITC) experiments showing three independent titrations of AIM *mut* peptide to MpATG8A. The concentration of reactants are 25 µM (2) or 50 µM (1,3) for MpATG8A (in cell) and 250 µM (2) or 500 µM (1,3) AIM *mut* peptide (in syringe). Note that replicate 1 for AIM *mut* peptide titration to MpATG8A is shown in Fig. 1F. **(B) MpATG8A^ADS^ does not bind AIM *wt*.** Isothermal titration calorimetry (ITC) experiments showing three independent titrations of AIM *wt* peptide to MpATG8A^ADS^. The concentration of reactants is 25 µM (1,2) or 50 µM (3) for MpATG8A^ADS^ (in cell) and 250 µM (1,2) or 500 µM (3) AIM *wt* peptide (in syringe). Note that replicate 1 for AIM *wt* peptide titration to MpATG8A^ADS^ is shown in Fig. 1G. **(C) MpATG8^ADS^ does not bind NBR1 *in vivo*.** Co-immunoprecipitation experiment of *M. polymorpha* transgenic plants expressing GFP-EV (GFP) or GFP-MpATG8 isoforms in wild*-type* (Tak-1) background. 7-day-old *M. polymorpha* gemmae were incubated at 37°C for 4 hours and recovered for additional 3 hours with fresh media at 21°C before harvesting. Lysates were incubated with GFP-Trap Magnetic Agarose beads. Input and bound proteins were analysed by immunoblotting using the indicated antibodies. AIM *wt* and AIM *mut* peptides were added to a final concentration of 200 µM. Total protein loading control was analysed by Amidoblack (AB) staining. **(D) MpATG8A does not bind UIM *wt*.** ITC experiments showing three independent titrations of UIM *wt* peptide to MpATG8A. The concentration of reactants in (A) are 50 µM (1,2,3) for MpATG8A (in cell) and 500 µM (1,2,3) UIM *wt* peptide (in syringe). Note that replicate 3 for UIM *wt* peptide titration to MpATG8A is shown in Fig. 1H. **(E) AtATG8A does not bind the UIM-containing AtRPN10 C-terminal portion (AtRPN10^201-386^**). ITC experiments showing three independent titrations of AtRPN10^201-386^ to AtATG8A. The concentration of reactants is 10 µM (1,2,3) for AtATG8A (in cell) and 100 µM (1,2,3) AtRPN10^201-386^ (in syringe). Global analysis was performed using a hetero-association model A + B. The top panels show the SVD-reconstructed thermograms, the middle panels show the isotherms as integrated heat of injection (dots) and best fit (solid line), and the bottom panels show the residuals (difference between the fit to a single hetero-association model and the experimental data; closer to zero indicates stronger agreement between data and the fit). Thermograms were reconstructed with NITPIC, global analysis was done in SEDPHAT, and data visualization was plotted in GUSSI.

**Figure S3.**
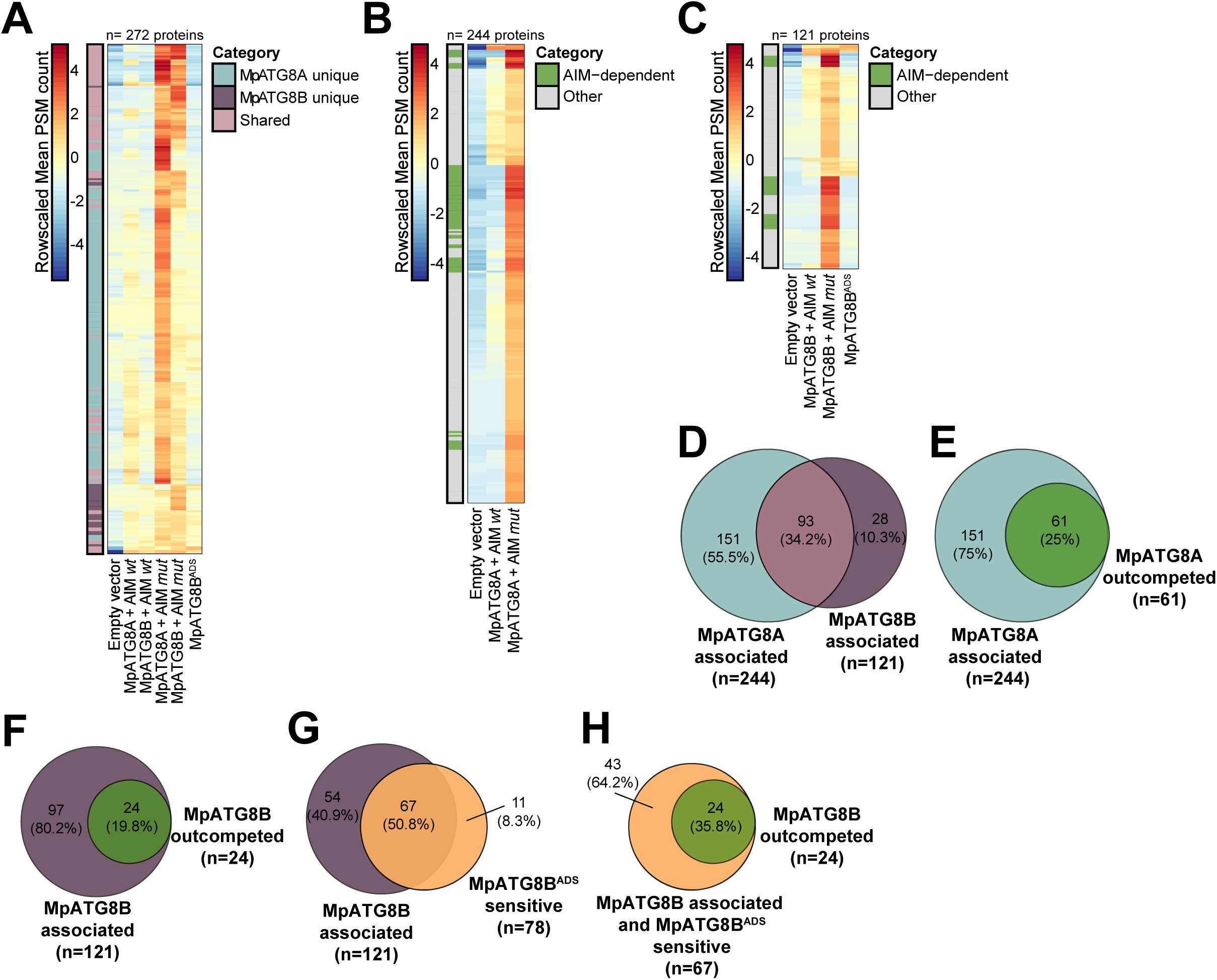
MpATG8 interactome analysis using peptide competition coupled AP-MS. **(A) Total MpATG8 interactome as defined by peptide competition.** Protein abundance pattern represented by a heatmap (Log2 (Mean PSM+1) – Mean PSM per protein) for the 272 MpATG8-associated proteins. Proteins are annotated based on their enrichment in one (MpATG8A/B unique) or multiple (shared) samples. Each column represents the row-scaled mean PSM count of 3 independent biological replicates for empty vector and MpATG8B^ADS^ and 2 independent biological replicates for AIM *wt* and AIM *mut* samples. **(B) MpATG8A interactome as defined by peptide competition.** Protein abundance pattern represented by a heatmap (Log2 (Mean PSM+1) – Mean PSM per protein) for the 242 MpATG8A**-**associated proteins. Proteins retained in MpATG8A+AIM *mut vs.* MpATG8A+AIM *wt* are annotated as AIM-dependent. Each column represents the row-scaled mean PSM count of 3 independent biological replicates for empty vector and 2 independent biological replicates for AIM *wt* and AIM *mut* samples. **(C) MpATG8B interactome as defined by peptide competition.** Protein abundance pattern represented by a heatmap (Log2 (Mean PSM+1) – Mean PSM per protein) for the 121 MpATG8B**-**associated proteins. Proteins retained in MpATG8B+AIM *mut vs.* MpATG8B+AIM *wt* are annotated as AIM-dependent. Each column represents the row-scaled mean PSM count of 3 independent biological replicates for empty vector and MpATG8B^ADS^ and 2 independent biological replicates for AIM *wt* and AIM *mut* samples. **(D) Comparison between the interactome of both MpATG8 isoforms.** Venn diagram of two overlapping pairwise comparisons. MpATG8A**-**associated proteins (aquamarine circle) are defined as in Fig. S3B. MpATG8B**-**associated proteins (purple circle) are defined as in Fig. S3C. **(E) Peptide competition reduces the MpATG8A interactome 4-fold.** Venn diagram of two overlapping pairwise comparisons. MpATG8A-associated proteins (aquamarine circle) are defined as in Fig. S3B. MpATG8A outcompeted proteins (green circle) are defined as AIM-dependent in Fig. S3B. **(F) Peptide competition reduces the MpATG8B interactome 5-fold.** Venn diagram of two overlapping pairwise comparisons. MpATG8B-associated proteins (purple circle) are defined as in Fig. S3C. MpATG8B outcompeted proteins (green circle) are defined as AIM-dependent in Fig. S3C. **(G) ADS mutation reduces the MpATG8B interactome.** Venn diagram of two overlapping pairwise comparisons. MpATG8B-associated proteins (purple circle) are defined as in Fig. S3C. MpATG8B^ADS^ sensitive proteins (orange circle) are defined as significantly depleted proteins in MpATG8B^ADS^. **(H) Outcompeted proteins are sensitive to ADS mutation.** Venn diagram of two overlapping pairwise comparisons. MpATG8B-associated and MpATG8B^ADS^ sensitive proteins (orange circle) are defined as the intersection in Fig. S3G (see also Fig. 1N). MpATG8B outcompeted proteins (green circle) are defined as AIM-dependent in Fig. S3C.

**Figure S4.**
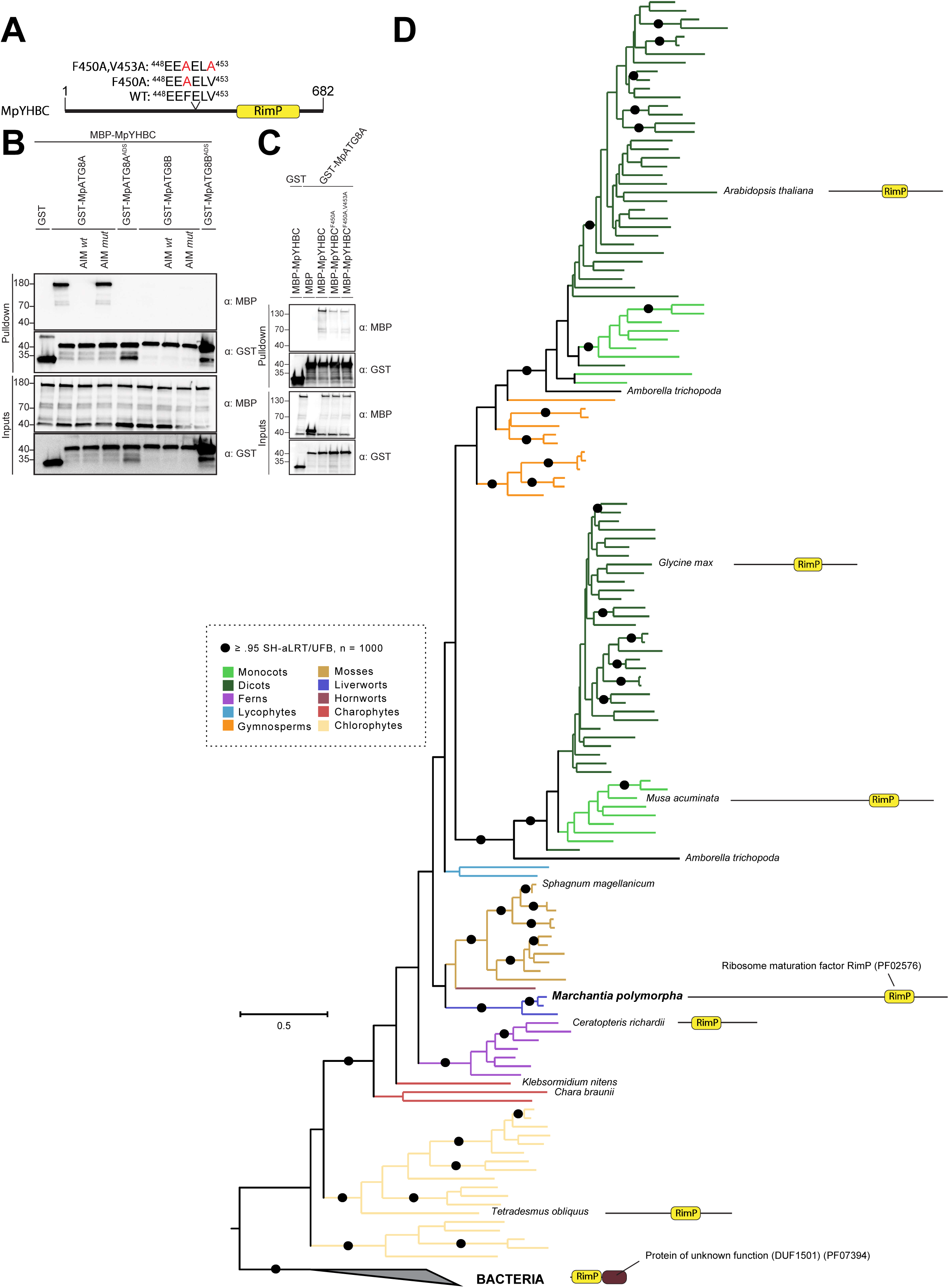
MpYHBC is a prokaryote-derived MpATG8 interactor. **(A) MpYHBC has a single AIM.** Protein domain architecture of MpYHBC. The ribosome maturation factor (RimP) domain is highlighted in yellow, AIM residues, their positions and mutagenesis are shown. **(B) MpYHBC binds MpATG8 in an AIM-dependent manner *in vitro*.** MpATG8A^ADS^=MpATG8A^(Y52A,L53A)^, MpATG8B^ADS^=MpATG8B^(K51A)^. AIM *wt* and AIM *mut* peptides were added to a final concentration of 200 µM. **(C) AIM mutagenesis disrupts MpATG8 binding.** Bacterial lysates containing recombinant protein were mixed and pulled down with glutathione magnetic agarose beads. Input and bound proteins were visualized by immunoblotting with anti-GST and anti-MBP antibodies. **(D) YHBC evolved from a prokaryotic ancestor.** Maximum likelihood phylogeny of 162 non-redundant YHBC homologs from 95 different plant and algal species and 16 different bacteria species constructed using the LG4M+R6 substitution model in IQ-TREE. Bacterial sequences are used for outgroup-based tree-rooting. Branches are colored for sequences belonging to the indicated clade/family, except *Amborella trichopoda* (black). The protein domain architecture of representative YHBC homologs is shown as represented in Fig. S4A, the bacterial domain of unknown function (DUF) is colored in brown. Shimodaira-Hasewaga with approximate likelihood-ratio test (SH-aLRT) and ultrafast bootstrap support (UFB) values (n = 1,000) greater than 95% are shown on the branches as black circles.

**Figure S5.**
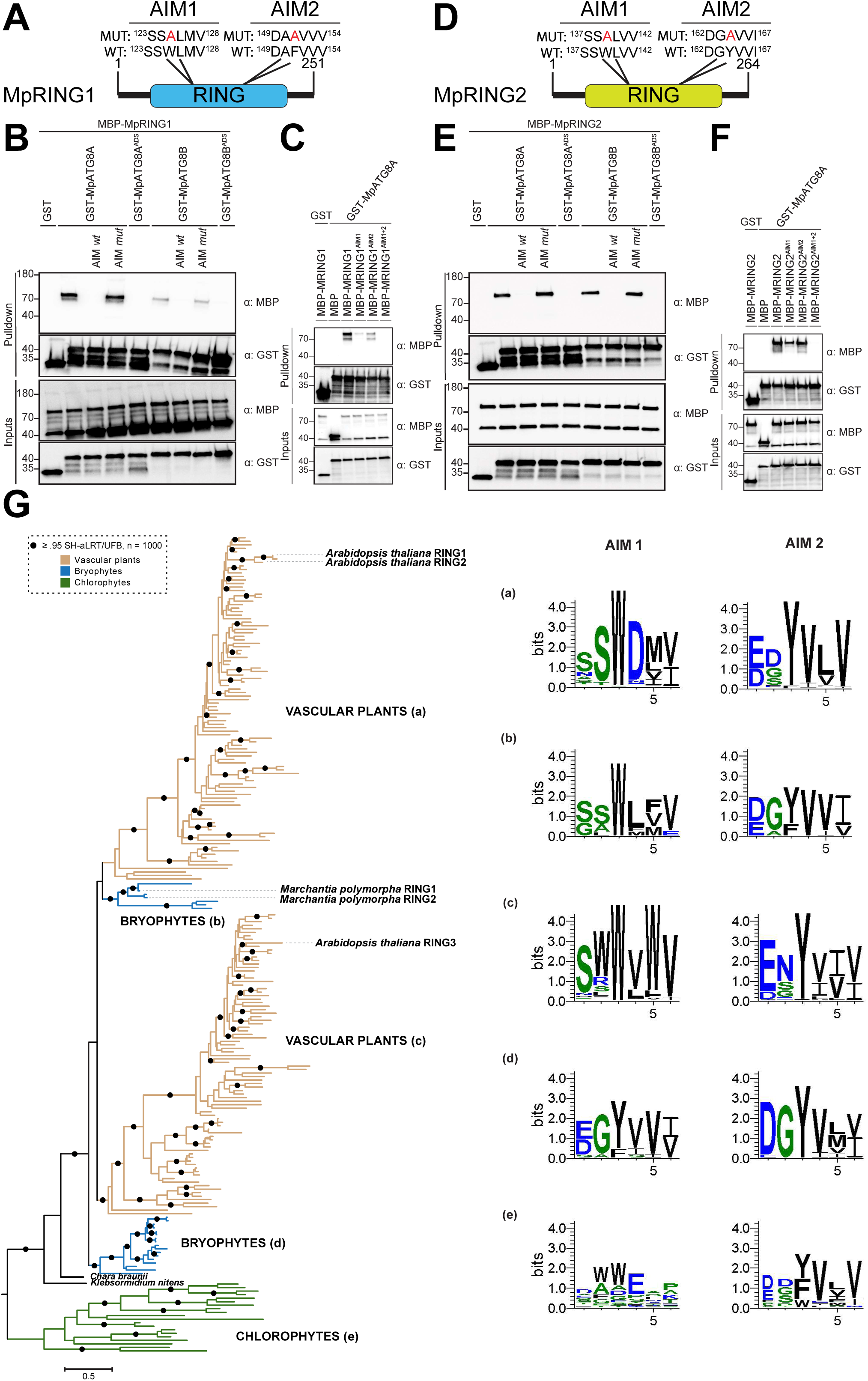
MpRING paralogs are MpATG8 interactors that evolved in the green lineage. **(A) MpRING1 has two AIMs in its RING domain.** Protein domain architecture of MpRING1. The RING domain is highlighted in blue, AIM residues, their positions and mutagenesis are shown. **(B) MpRING1 binds MpATG8 in an AIM-dependent manner *in vitro*.** MpATG8A^ADS^=MpATG8A^(Y52A,L53A)^, MpATG8B^ADS^=MpATG8B^(K51A)^. AIM *wt* and AIM *mut* peptides were added to a final concentration of 200 µM. **(C) AIM mutagenesis abolishes MpATG8 binding.** MpRING1^AIM1^=MpRING1^(W125A)^, MpRING1^AIM2^=MpRING1^(F151A)^, MpRING1^AIM1+2^=MpRING1^(W125A,F151A)^. Bacterial lysates containing recombinant protein were mixed and pulled down with glutathione magnetic agarose beads. Input and bound proteins were visualized by immunoblotting with anti-GST and anti-MBP antibodies. **(D) MpRING2 has two AIMs in its RING domain.** Protein domain architecture of MpRING2. The RING domain is highlighted in yellow, AIM residues and their positions are shown. **(E) MpRING2 binds MpATG8 in an AIM-dependent manner *in vitro*.** MpATG8A^ADS^=MpATG8A^(Y52A,L53A)^, MpATG8B^ADS^=MpATG8B^(K51A)^. AIM *wt* and AIM *mut* peptides were added to a final concentration of 200 µM. **(F) AIM mutagenesis abolishes MpATG8 binding.** MpRING2^AIM1^=MpRING2^(W139A)^,MpRING2^AIM2^=MpRING2^(Y164A)^,MpRING2^AIM1+2^=MpRING2^(W139A,Y164A)^. Bacterial lysates containing recombinant protein were mixed and pulled down with glutathione magnetic agarose beads. Input and bound proteins were visualized by immunoblotting with anti-GST and anti-MBP antibodies. **(G) RING paralogs evolved from a green algal progenitor.** Maximum likelihood phylogeny of 237 non-redundant RING domain-containing protein homologs from 101 different plant and algal species constructed using the LG4M+R7 substitution model in IQ-TREE (left). Chlorophytes are used for outgroup-based tree-rooting. Branches are colored for sequences belonging to the indicated clade/family, except charophytes *Chara braunii* and *Klebsormidium nitens* (black). *M. polymorpha* and *A. thaliana* RING protein homologs are highlighted. Shimodaira-Hasewaga with approximate likelihood-ratio test (SH-aLRT) and ultrafast bootstrap support (UFB) values (n = 1,000) greater than 95% are shown on the branches as black circles. Sequence logos (right) show aminoacid conservation for AIM1 and AIM2 across the RING protein homologs in the different clades (a-e).

**Figure S6.**
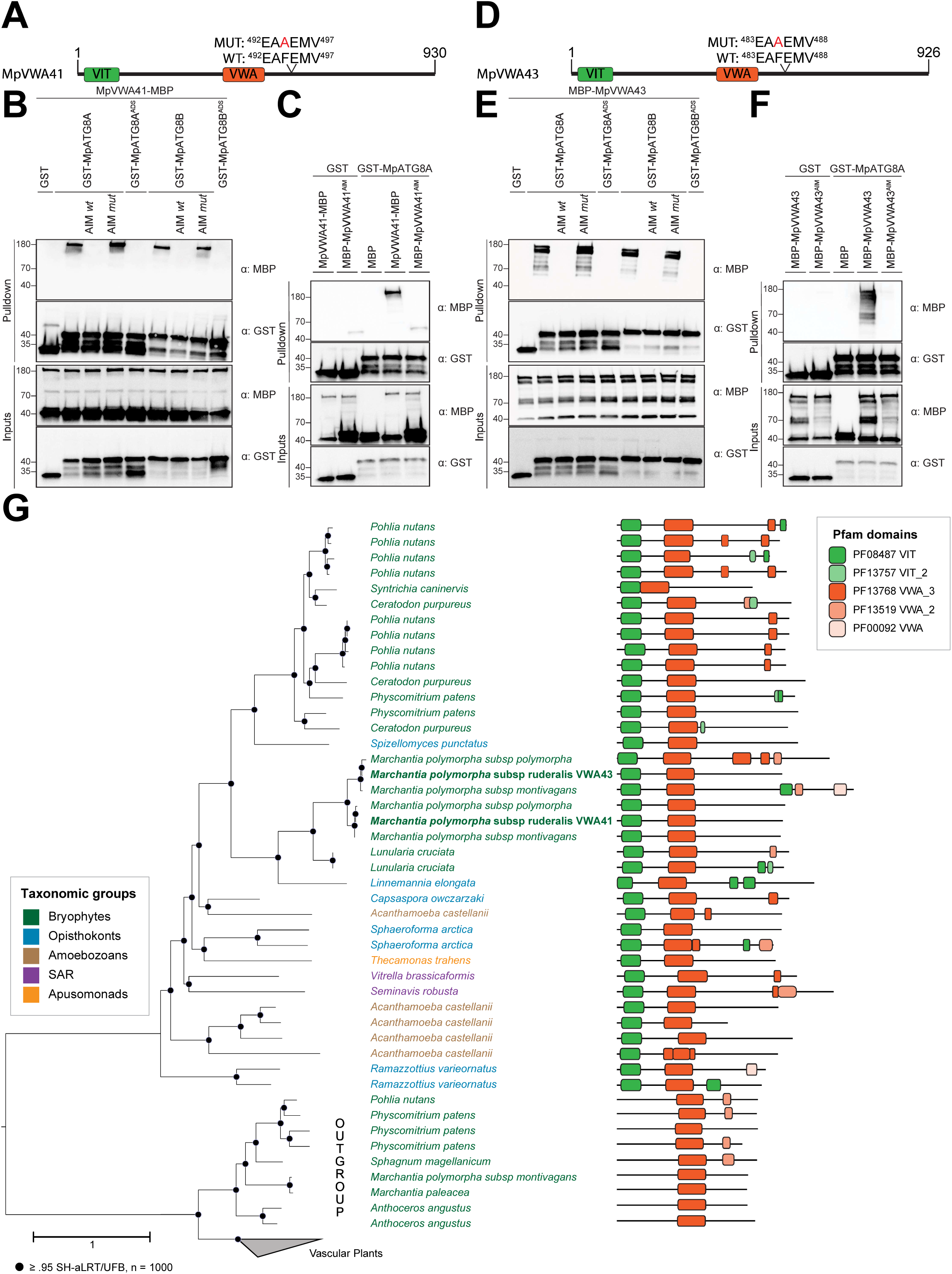
MpVWA paralogs are bryophyte-specific MpATG8 interactors. **(A) MpVWA41 has a single AIM.** Protein domain architecture of MpVWA41. The **V**ault-**I**nter **T**rypsin (VIT) and **v**on **W**illebrand factor (VWA) domains are highlighted in green and red, respectively. AIM residues, their positions and mutagenesis are shown. **(B) MpVWA1 binds MpATG8 in an AIM-dependent manner *in vitro*.** MpATG8A^ADS^=MpATG8A^(Y52A,L53A)^, MpATG8B^ADS^=MpATG8B^(K51A)^. AIM *wt* and AIM *mut* peptides were added to a final concentration of 200 µM. **(C) AIM mutagenesis abolishes MpATG8 binding.** MpVWA41^AIM^=MpVWA41^(F494A)^. Bacterial lysates containing recombinant protein were mixed and pulled down with glutathione magnetic agarose beads. Input and bound proteins were visualized by immunoblotting with anti-GST and anti-MBP antibodies. **(D) MpVWA43 has a single AIM.** Protein domain architecture of MpVWA43. The VIT and VWA domains are highlighted in green and red, respectively. AIM residues and their positions are shown. **(E) MpVWA43 binds MpATG8 in an AIM-dependent manner *in vitro*.** MpATG8A^ADS^=MpATG8A^(Y52A,L53A)^, MpATG8B^ADS^=MpATG8B^(K51A)^. AIM *wt* and AIM *mut* peptides were added to a final concentration of 200 µM. **(F) AIM mutagenesis abolishes MpATG8 binding.** MpVWA43^AIM^=MpVWA43^(F485A)^. Bacterial lysates containing recombinant protein were mixed and pulled down with glutathione magnetic agarose beads. Input and bound proteins were visualized by immunoblotting with anti-GST and anti-MBP antibodies. **(G) VWA paralogs evolved independently across the eukaryotic tree of life.** Maximum likelihood phylogeny of 84 non-redundant VWA protein homologs from 46 different eukaryotic species constructed using the LG4M+I+R6 substitution model in IQ-TREE. A clade of an unrelated VWA-containing protein conserved in land plants is used for outgroup-based tree-rooting. Branches are colored for sequences belonging to the indicated taxonomic group. The protein domain architecture of representative VWA homologs is shown as represented in Fig. S6A, other annotated Pfam domains are also shown. Shimodaira-Hasewaga with approximate likelihood-ratio test (SH-aLRT) and ultrafast bootstrap support (UFB) values (n = 1,000) greater than 95% are shown on the branches as black circles.

**Figure S7.**
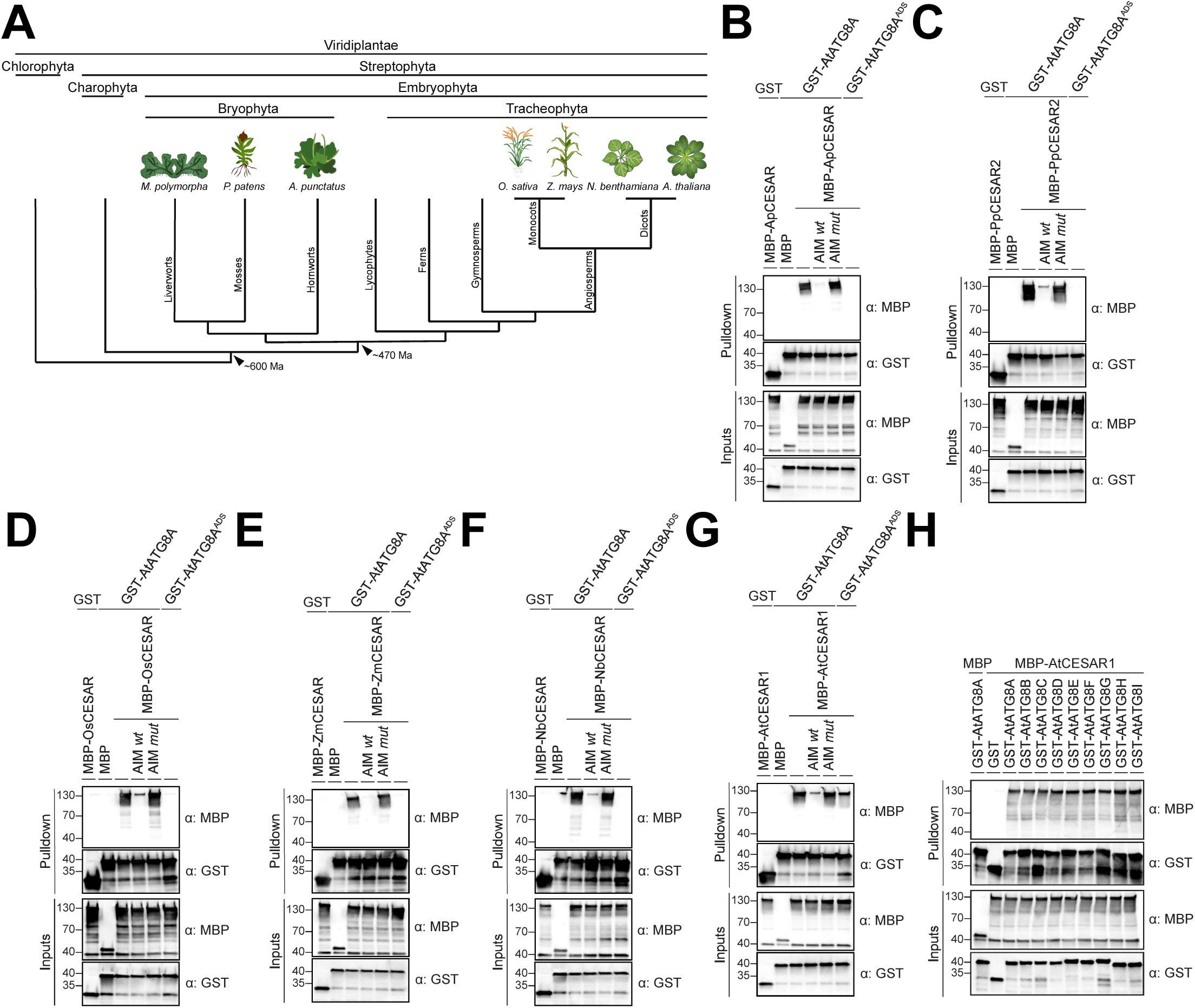
CESAR is an evolutionary conserved ATG8 interactor. **(A) Phylogenetic relationships between plant species tested.** Schematic phylogram of the Viridiplantae kingdom and its phylum. Species used for testing evolutionary conservation of CESAR interaction with ATG8 are represented. **(B-G) CESAR orthologs bind to AtATG8A in an AIM-dependent manner *in vitro*.** AtATG8a^ADS^=AtATG8a^(Y50A,L51A)^. AIM *wt* and AIM *mut* peptides were added to a final concentration of 200 µM. **(H) AtCESAR1 binds all AtATG8 isoforms *in vitro*.** Bacterial lysates containing recombinant protein were mixed and pulled down with glutathione magnetic agarose beads. Input and bound proteins were visualized by immunoblotting with anti-GST and anti-MBP antibodies.

**Figure S8.**
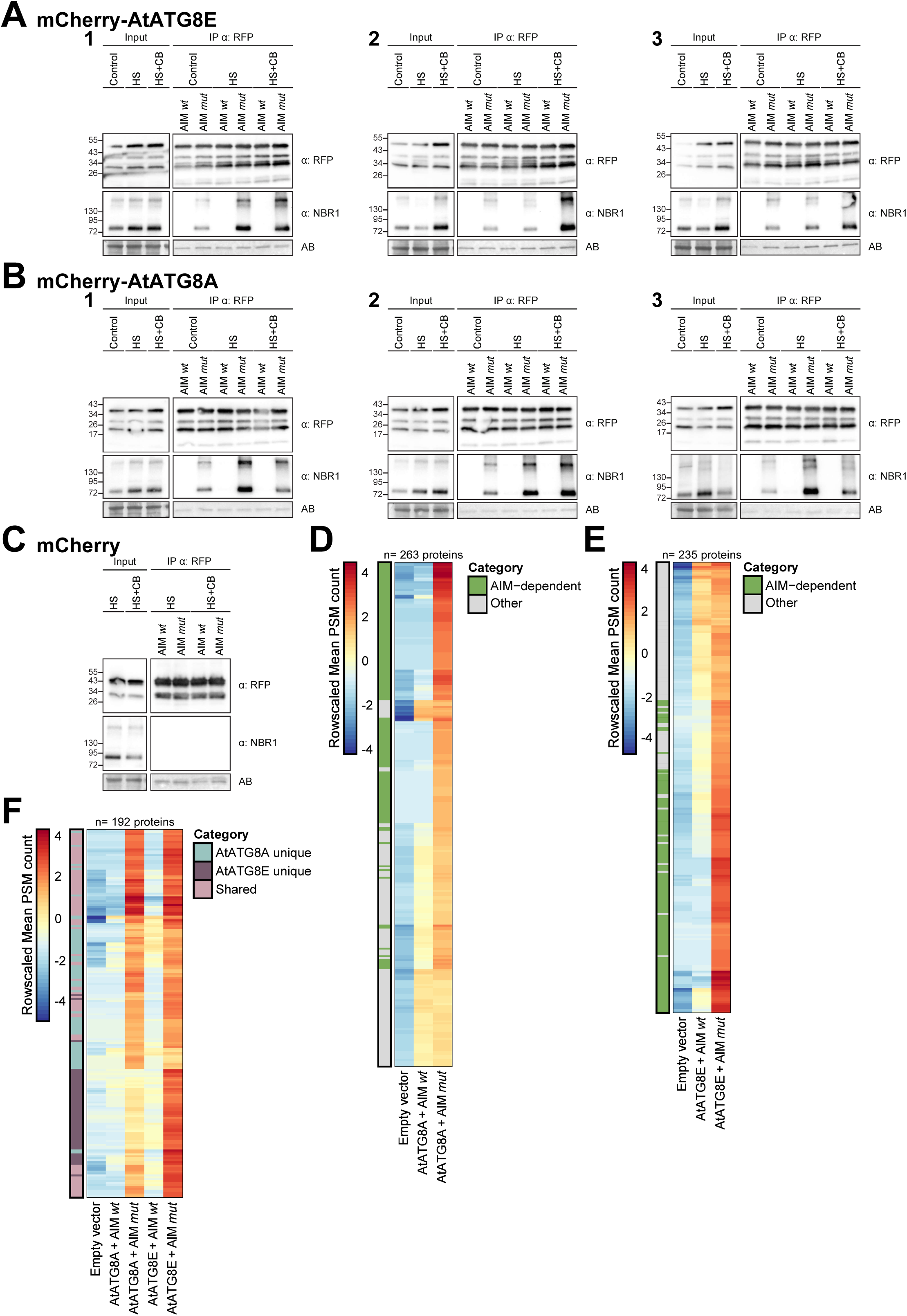
AtATG8 interactome analysis using peptide competition coupled with AP-MS. **(A-C) AtATG8 immunoprecipitation upon HS/HS+CB treatment.** 7-d-old *A. thaliana* seedlings expressing either mCherry-AtATG8E (A), mCherry-AtATG8A (B) or mCherry-EV (C) were incubated at 21°C (Control) or 37°C (Heat stress, HS) conditions for 6h followed by a 4h recovery at 21°C in fresh ½ MS + MES + 1% sucrose media supplemented with either DMSO or 50 µM CB5083 (HS+CB). AIM *wt* and AIM *mut* peptides were added to a final concentration of 100 μM. Protein extracts were immunoblotted with anti-RFP and anti-NBR1 antibodies. Reference protein sizes are indicated on the left side of the blots (unit: kDa). Total protein loading control was analysed by Amidoblack (AB) staining. **(D) Peptide competition enriches for direct AtATG8A interactors.** Protein abundance pattern represented by a heatmap (Log2 (Mean PSM+1) – Mean PSM per protein) for the 263 AtATG8A-associated proteins. Proteins retained in the AIM *mut vs*. AIM *wt* comparison are annotated as AIM-dependent. Each column represents the row-scaled mean PSM count of 3 independent biological replicates. **(E) Peptide competition enriches for direct AtATG8E interactors.** Protein abundance pattern represented by a heatmap (Log2 (Mean PSM+1) – Mean PSM per protein) for the 235 AtATG8E-associated proteins. Proteins retained in the AIM *mut vs*. AIM *wt* comparison are annotated as AIM-dependent. Each column represents the row-scaled mean PSM count of 3 independent biological replicates. **(F) Comparison between AIM-dependent interactors of AtATG8A and AtATG8E.** Protein abundance pattern represented by a heatmap (Log2 (Mean PSM+1) – Mean PSM per protein) for the 192 outcompeted proteins identified for either of both AtATG8 isoforms, defined as AIM-dependent in Fig. S8D,E. Proteins are annotated based on their enrichment in one (AtATG8A/E unique) or multiple (shared) samples. Each column represents the row-scaled mean PSM count of 3 independent biological replicates.

**Figure S9.**
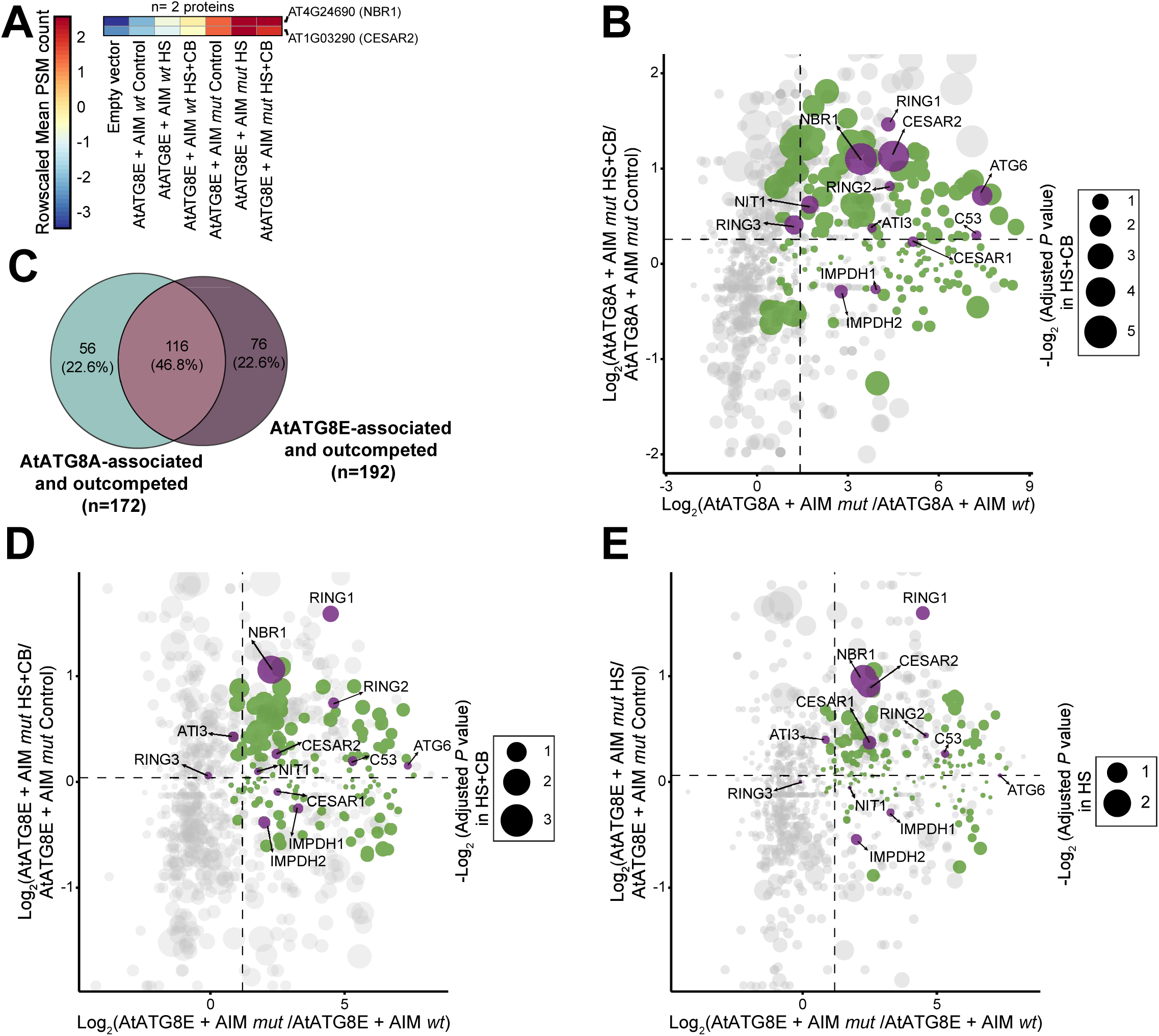
HS and HS+CB treatment affect the interactome of AtATG8A and AtATG8E isoforms. **(A) HS+CB treatment induces AIM-dependent interactions of AtCESAR2 and AtNBR1 with AtATG8E.** Protein abundance pattern represented by a heatmap (Log2 (Mean PSM+1) – Mean PSM per protein) for the 2 AtATG8E-associated and outcompeted proteins identified as induced upon HS+CB. Each column represents the row-scaled mean PSM count of 3 independent biological replicates. **(B) HS+CB treatment induces AIM-dependent interactions of AtCESAR2 with AtATG8A.** Dot plot representing protein abundance as Log2 (Fold change) for the ratio of two pairwise comparisons comparing the peptide effect (AtATG8A+AIM *mut vs.* AtATG8A+AIM *wt*) on the x-axis and the treatment effect (AtATG8A+AIM *mut* HS+CB *vs.* AtATG8A+AIM *mut* Control) on the y-axis. The centroid for x-axis and y-axis is represented as dashed lines. Dot size is mapped to reflect significant induction in HS+CB-treated samples represented as – Log2 (Adjusted *P* value). The 172 AtATG8A-associated and outcompeted proteins (annotated as AIM-dependent in Fig. S8D) are colored in green, proteins of interest are colored in purple. **(C) Comparison between AtATG8A and AtATG8E interactomes.** Venn diagram of two overlapping pairwise comparisons. AtATG8A-associated and outcompeted proteins (yellow circle) are defined as AIM-dependent in Fig. S8D. AtATG8E-associated and outcompeted proteins (pink circle) are defined as AIM-dependent in Fig. S8E. **(D-E) HS+CB induces AtCESAR2-AtATG8E interaction.** Dot plot representing protein abundance as Log2 (Fold change) for the ratio of two pairwise comparisons comparing the peptide effect (AtATG8E+AIM *mut vs.* AtATG8E+AIM *wt*) on the x-axis and the treatment effect (D: AtATG8E+AIM *mut* HS+CB / E: AtATG8E+AIM *mut* HS *vs.* AtATG8E+AIM *mut* Control) on the y-axis. The centroid for x-axis and y-axis is represented as dashed lines. Dot size is mapped to reflect significant induction in HS+CB (D) or HS (E)-treated samples represented as – Log2 (Adjusted *P* value). The 192 AtATG8E-associated and outcompeted proteins (annotated as AIM-dependent in Fig. S8E) are colored in green, proteins of interest are colored in purple.

**Figure S10.**
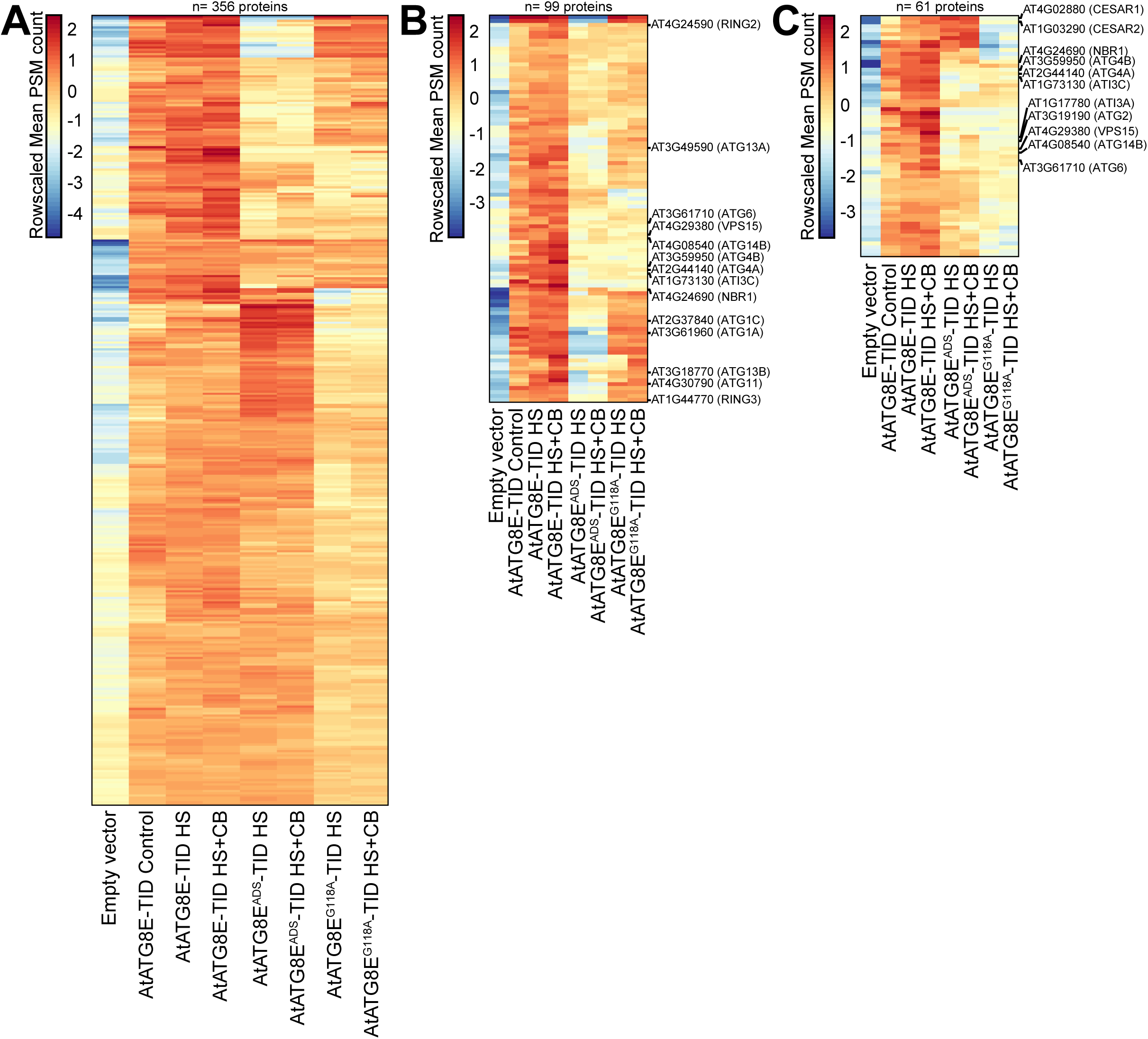
Turbo-ID-based proximity-labelling identifies the AtATG8E proxitome. **(A) AtATG8E proxitome as defined by Turbo-ID-based proximity-labelling.** Protein abundance pattern represented by a heatmap (Log2 (Mean PSM+1) – Mean PSM per protein) for the 356 AtATG8E-TID proximal proteins. Each column represents the row-scaled mean PSM count of 3 independent biological replicates. **(B) ADS mutation reduces the AtATG8E proxitome.** Protein abundance pattern represented by a heatmap (Log2 (Mean PSM+1) – Mean PSM per protein) for the 99 AtATG8E-TID proximal proteins identified as depleted in AtATG8E^ADS^-TID. Each column represents the row-scaled mean PSM count of 3 independent biological replicates. **(C) AtATG8E C-terminal glycine mutation highlights conjugated ATG8 sensitive neighbour proteins.** Protein abundance pattern represented by a heatmap (Log2 (Mean PSM+1) – Mean PSM per protein) for the 61 AtATG8E-TID proximal proteins identified as depleted in AtATG8E^G118A^-TID. Each column represents the row-scaled mean PSM count of 3 independent biological replicates.

**Figure S11.**
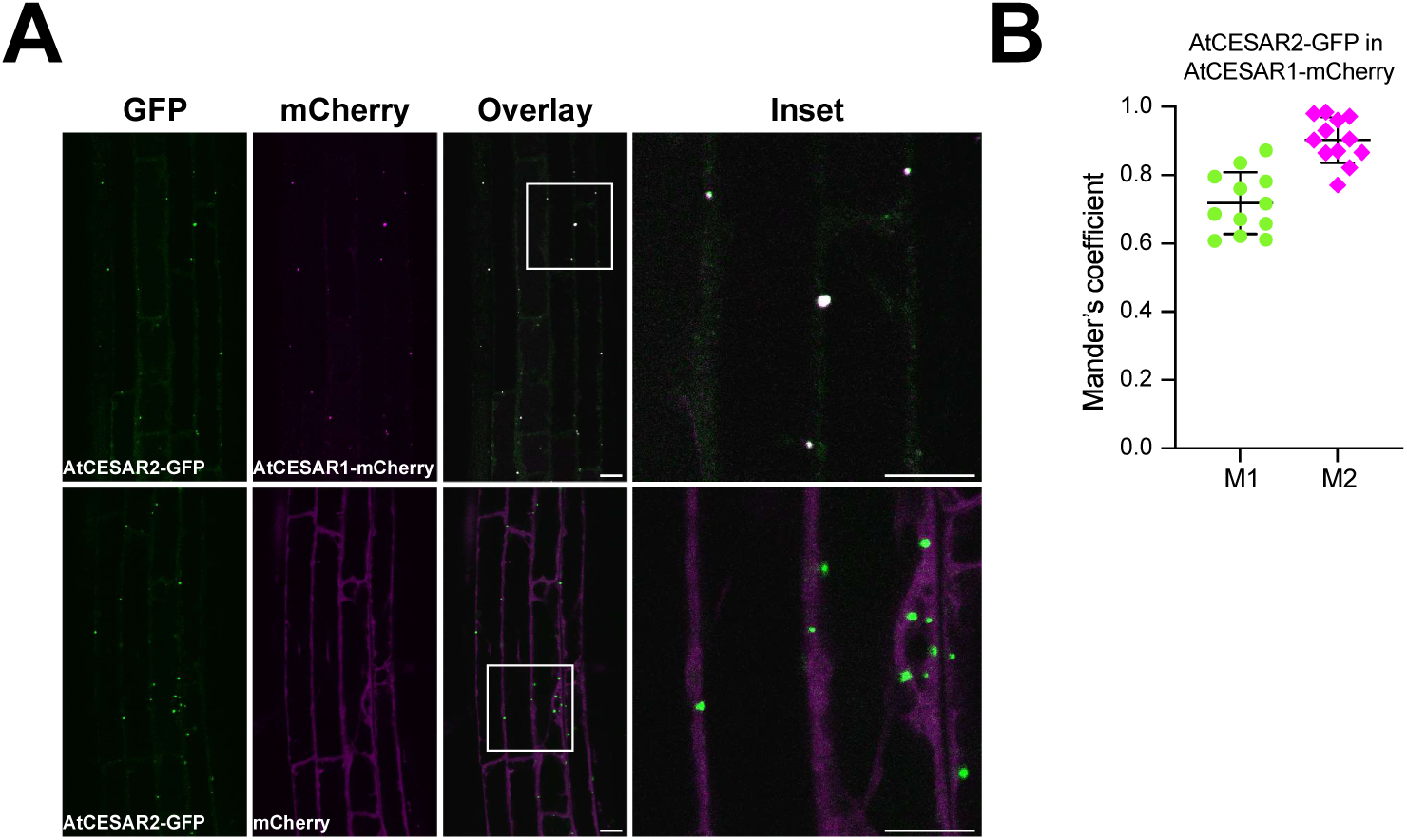
AtCESAR paralogs colocalize with each other. **(A) AtCESAR1 and AtCESAR2 co-localize**. Confocal microscopy images of *A. thaliana* root epidermal cells co-expressing AtCESAR2-GFP with either AtCESAR1-mCherry or mCherry-EV (mCherry). 5-d-old *A. thaliana* seedlings were grown in 1% agar ½ MS + MES + 1% Sucrose plates under control (21°C) conditions prior to imaging. Area highlighted in the white-boxed region in the merge panel was further enlarged and presented in the inset panel. Scale bars, 10 μm. Inset scale bars, 10 μm. **(B)** Quantification of confocal experiments in Fig. S11A as assessed by the Mander’s colocalization coefficients between AtCESAR2-GFP and AtCESAR1-mCherry. M1, fraction of AtCESAR2-GFP signal that overlaps with AtCESAR1-mCherry signal. M2, fraction of AtCESAR1-mCherry signal that overlaps with AtCESAR2-GFP signal. Bars indicate the mean ± SD of 12 biological replicates.

**Figure S12.**
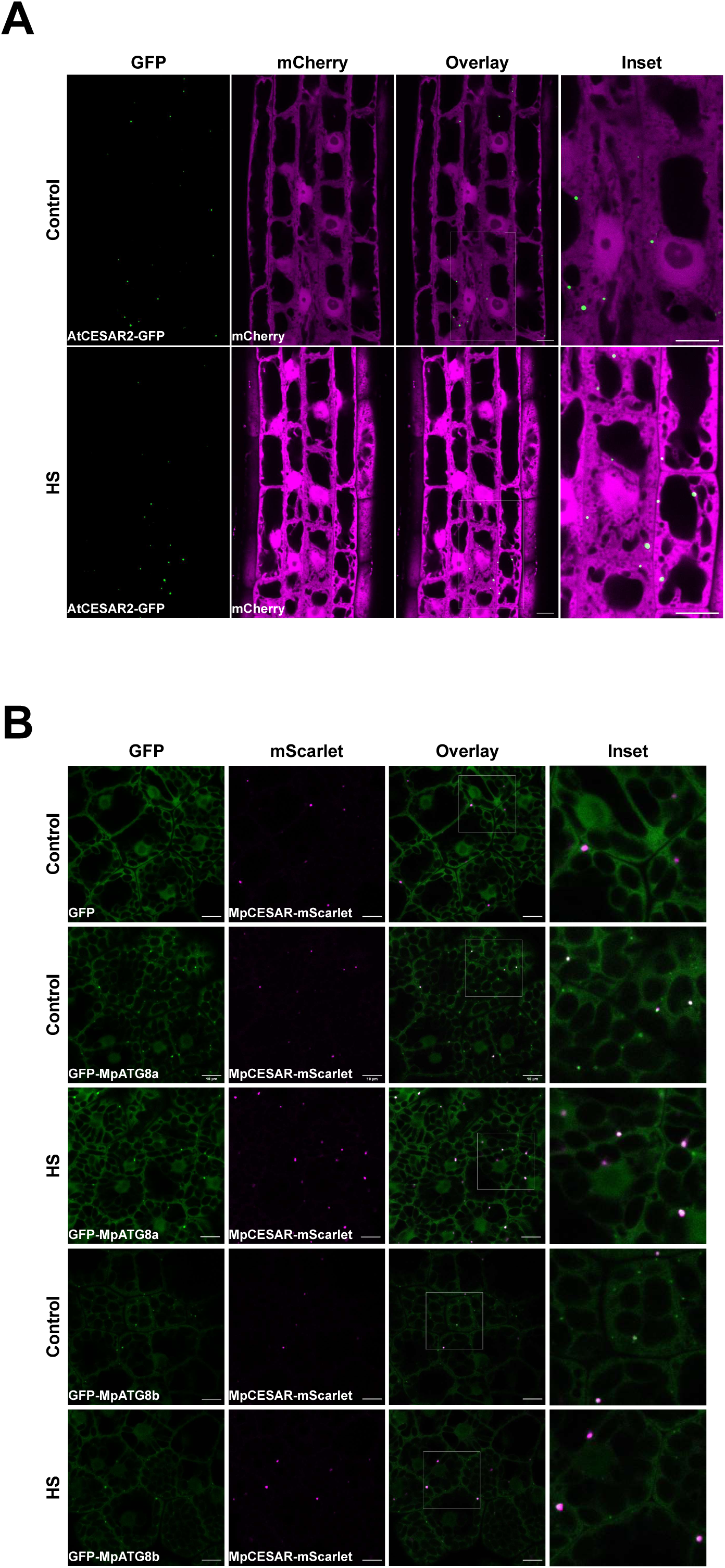
CESAR localizes to autophagosomes. **(A) AtCESAR2 does not co-localize with mCherry-EV control.** Confocal microscopy images of *A. thaliana* root epidermal cells at the transition zone co-expressing AtCESAR2-GFP with mCherry-EV. 5-d-old *A. thaliana* seedlings were grown in 1% agar ½ MS + MES + 1% sucrose plates and incubated at 21°C (Control, C) or 37°C (Heat stress, HS) for 3h followed by a recovery of 3h at 21°C prior to imaging. Representative images of 3 biological replicates are shown. Area highlighted in the white-boxed region in the merged panel was further enlarged and presented in the inset panel. Scale bars, 10 μm. Inset scale bars, 10 μm. **(B) MpCESAR co-localizes with MpATG8-decorated autophagosomes**. Confocal microscopy images of *M. polymorpha* thallus cells co-expressing MpCESAR-mScarlet with either GFP-EV, GFP-MpATG8A or GFP-MpATG8B. 2-d-old thalli were incubated in ½ Gamborg B5 media for 1 hour at 21°C (Control) or 37°C (Heat stress, HS) followed by a 1-hour recovery phase at 21°C before imaging. Representative images of 10 biological replicates are shown. Inset panels show enlarged sections of the white-boxed areas highlighted in the merge panel. Scale bar,10 µm.

**Figure S13.**
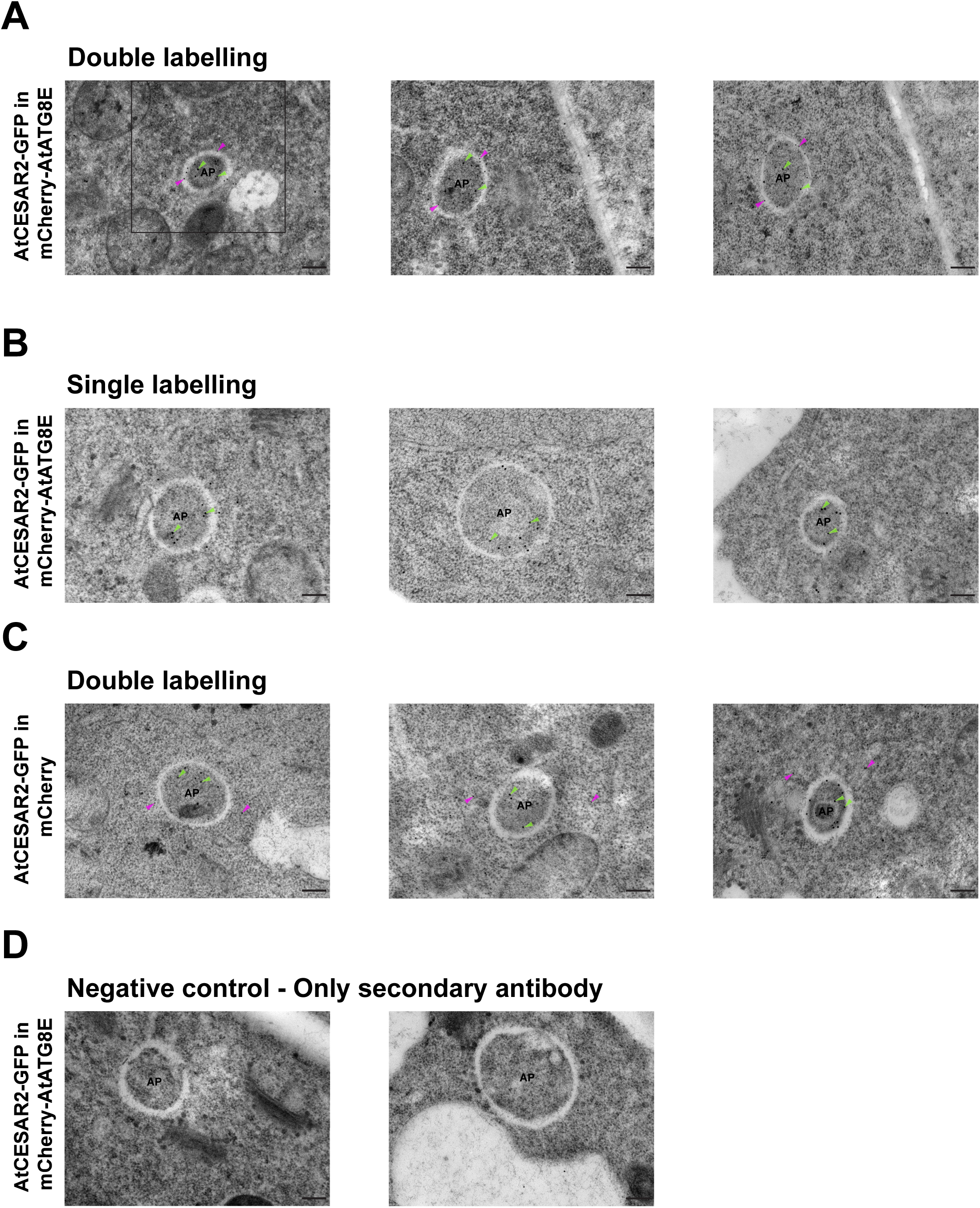
Transmission electron microscopy illuminates CESAR localization inside autophagosomes. **(A-D) CESAR localizes to the autophagosome lumen.** Transmission electron microscopy (TEM) micrographs showing either immune-gold labelled AtCESAR2-GFP (15 nm gold particles, green arrows) and mCherry-AtATG8E or mCherry-EV (10 nm gold particles, magenta arrows), in the cytoplasm of *A. thaliana* root cells. 5-d-old *A. thaliana* seedlings were grown at 21°C in 1% agar ½ MS + MES + 1% sucrose plates before cryofixation. Sections were labelled with anti-GFP and anti-mCherry primary antibodies and secondary antibodies conjugated with either 15 nm gold particles (for GFP, green arrows) or 10 nm gold particles (for mCherry, magenta arrows). Area highlighted in the first panel was used as a representative image in the main figure. Scale bars, 200 nm. AP, autophagosome.

**Figure S14.**
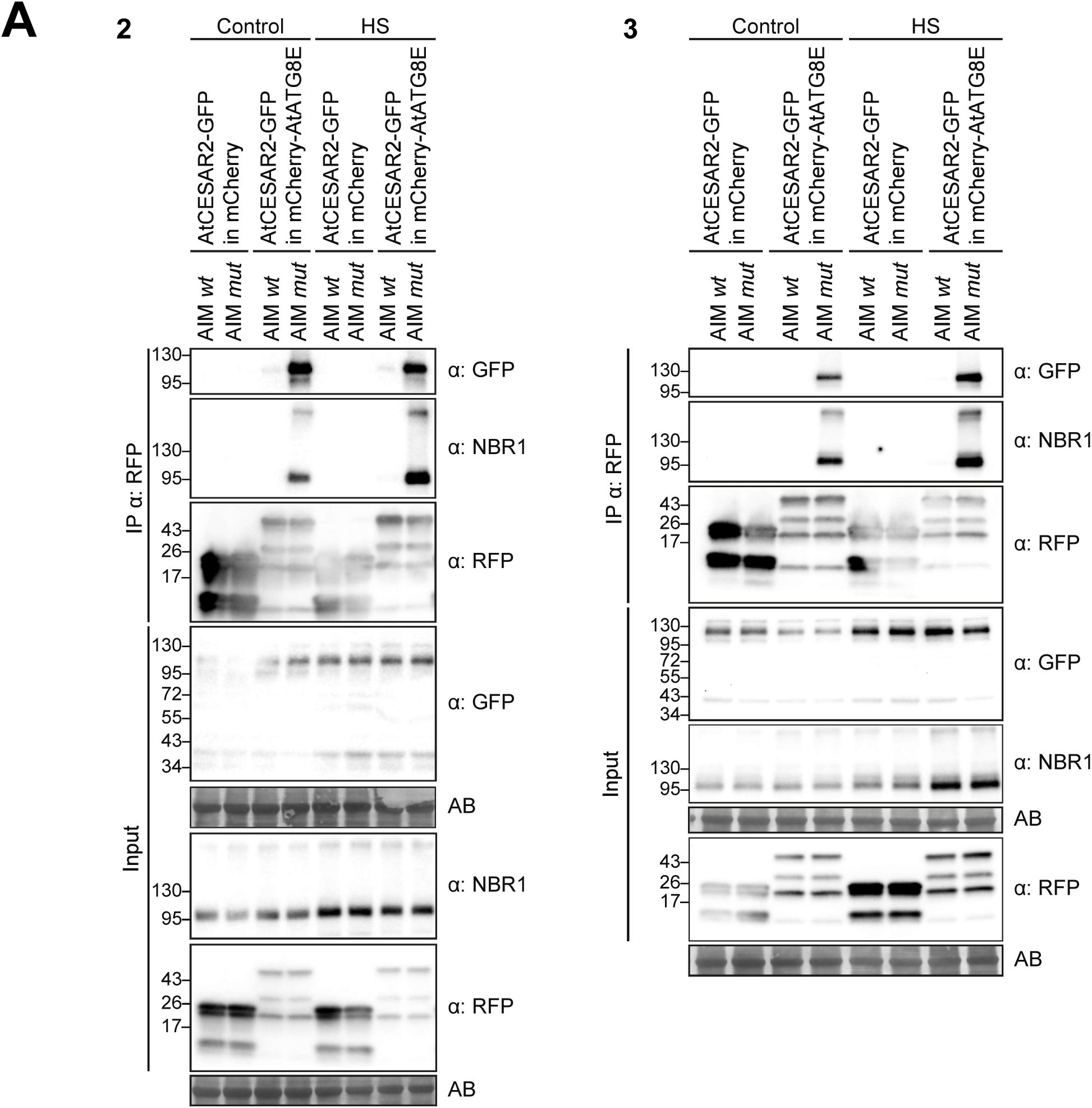
AtCESAR2 co-immunoprecipitates with AtATG8E *in vivo*. Two independent biological replicates (2 and 3) of RFP-Trap co-immunoprecipitation coupled to peptide competition of whole-seedling extracts of 7-d-old *A. thaliana* seedlings co-expressing AtCESAR2-GFP with either mCherry-AtATG8E or mCherry-EV. Replicate 1 is presented in Figure 3H. Seedlings were incubated either at 21°C (Control) or 37°C (Heat stress, HS) for 6 hours followed by a 4-h recovery at 21°C in fresh ½ MS + MES + 1% Sucrose media. AIM *wt* and AIM *mut* peptides were added to a final concentration of 100 μM. Protein extracts were immunoblotted with the indicated antibodies. Reference protein sizes are indicated on the left side of the blots (unit: kDa). Total protein loading control was analysed by Amidoblack (AB) staining.

**Figure S15.**
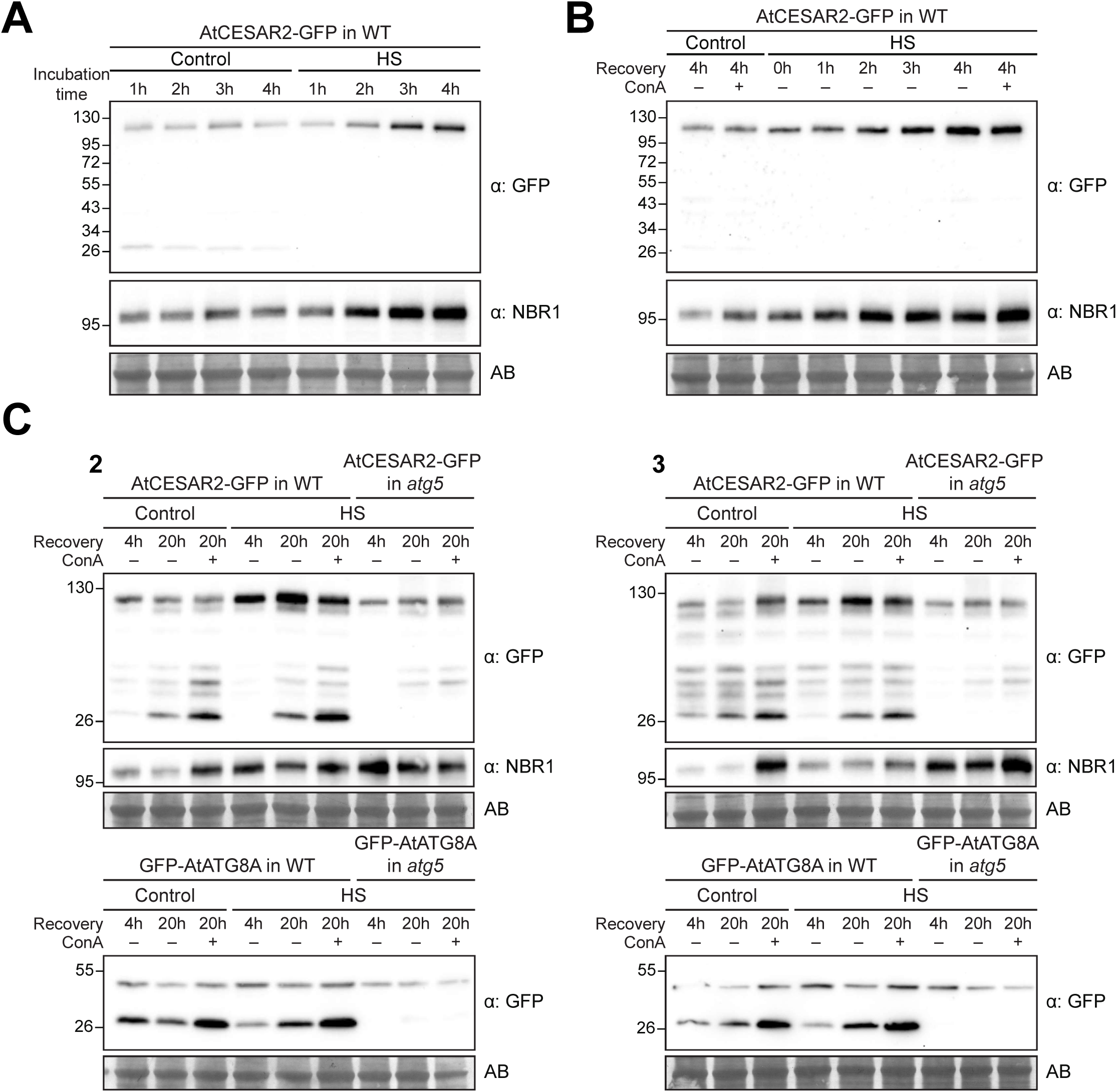
AtCESAR2 protein levels are sensitive to HS. **(A) HS stabilizes AtCESAR2 levels.** Western blot showing AtCESAR2-GFP protein levels, and endogenous NBR1 levels. *A. thaliana* seedlings were grown in ½ MS + MES + 1% sucrose media at 21°C media for 7 days and incubated either at 21°C (Control) or 37°C (Heat stress, HS) for the indicated time. 20 μg of total protein extract was loaded and immunoblotted with anti-GFP and anti-NBR1 antibodies. Reference protein sizes are indicated on the left side of the blots (unit: kDa). Total protein loading control was analysed by Amidoblack (AB) staining. **(B) HS followed by recovery stabilizes AtCESAR2 levels.** Western blot showing AtCESAR2-GFP and endogenous NBR1 protein levels. levels. *A. thaliana* seedlings were grown in ½ MS + MES + 1% sucrose media for 7 days and incubated either at 21°C (Control) or 37°C (Heat stress, HS) conditions for 4h, followed by the indicated recovery times at 21°C in fresh media ± 1 μM Concanamycin A (ConA). 20 μg of total protein extract was loaded and immunoblotted with anti-GFP and anti-NBR1 antibodies. Reference protein sizes are indicated on the left side of the blots (unit: kDa). Total protein loading control was analysed by Amidoblack (AB) staining. **(C) AtCESAR2 undergoes autophagic degradation during prolonged recovery post-heat stress.** Two independent biological replicates (2 and 3) from the experiment shown in Fig. 3K. *A. thaliana* seedlings were grown in 1/2 MS media for 7 days and incubated either at 21°C (Control) or 37°C (Heat stress, HS) for 4h, followed either by 4h or 20h recovery in fresh ½ MS + MES +1% sucrose media ± 1 μM ConA at 21°C. 20 μg of total protein extract was loaded and immunoblotted with the indicated antibodies. Reference protein sizes are indicated on the left side of the blots (unit: kDa). Total protein loading control was analysed by Amidoblack (AB) staining.

**Figure S16.**
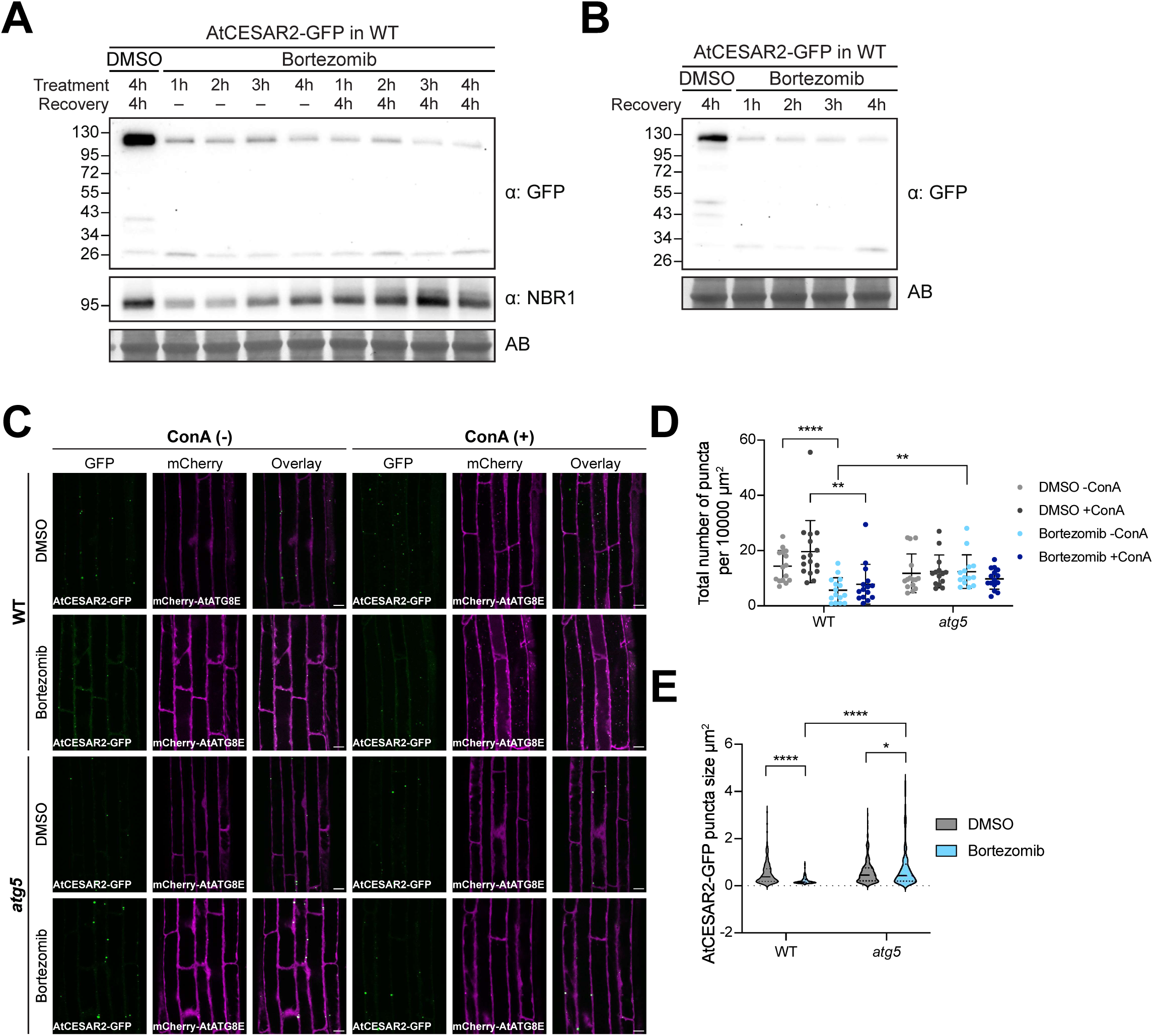
AtCESAR2 protein levels, puncta number and size decrease upon proteasome inhibition in an autophagy dependent manner. **(A) Proteasome inhibition induces AtCESAR2 degradation.** Western blot showing AtCESAR2-GFP protein levels, and endogenous NBR1 levels upon bortezomib treatment. *A. thaliana* seedlings were grown in ½ MS + MES +1% sucrose media for 7 days before incubation at 21°C in media supplemented with either DMSO (Control) for 4h followed by a 4h recovery in fresh media or with 50 µM Bortezomib for the indicated times followed by either no recovery or 4h recovery in fresh media. 20 μg of total protein extract was loaded and immunoblotted with anti-GFP and anti-NBR1 antibodies. Reference protein sizes are indicated on the left side of the blots (unit: kDa). Total protein loading control was analysed by Amidoblack (AB) staining. **(B) AtCESAR2 levels do not recover after proteasome inhibition.** Western blot showing AtCESAR2-GFP protein levels after a 4h bortezomib treatment followed by 1h to 4h recovery. *A. thaliana* seedlings were grown in ½ MS + MES + 1% sucrose media for 7 days and incubated at 21°C in media supplemented with either DMSO (Control) or 50 µM Bortezomib for 4h, followed by the indicated recovery time in fresh media. 20 μg of total protein extract was loaded and immunoblotted with anti-GFP antibody. Reference protein sizes are indicated on the left side of the blots (unit: kDa). Total protein loading control was analysed by Amidoblack (AB) staining. **(C) Proteasome inhibition induces AtCESAR2 degradation in an autophagy-dependent manner.** Confocal microscopy images of *A. thaliana* root epidermal cells co-expressing AtCESAR2-GFP with mCherry-AtATG8E in wild*-*type (WT) and *atg5* mutant backgrounds. 5-d-old *A. thaliana* seedlings were incubated for 2h at 21°C in ½ MS + MES + 1% Sucrose media supplemented with 50 µM Bortezomib or DMSO, followed by a 2.5h recovery in fresh media ± 1 μM Concanamycin A (ConA). Representative images of 14-16 replicates per condition are shown. Scale bars, 10 μm. **(D) Proteasome inhibition decreases the AtCESAR2 puncta number.** Quantification of confocal experiments in Fig. S16C showing the total number of AtCESAR2-GFP puncta per normalized area (10,000 μm^2^). Bars indicate the mean ± SD of 15 replicates for both wild-type (WT) and *atg5* (DMSO – ConA); 16 replicates for WT and 14 for *atg5* (DMSO + ConA); 15 replicates for WT and 14 for *atg5* (Bortezomib – ConA); 15 replicates for WT and *atg5* (Bortezomib +ConA). Two-tailed unpaired Student’s t-tests with Welch’s corrections were performed to analyze the differences of total puncta numbers between DMSO or Bortezomib within and between the genotypes ± 1 µM ConA. ****, *P* value < 0.0001, **, *P* value < 0.005 (0.0028). **(E) Proteasome inhibition reduces the AtCESAR2 puncta size.** Quantification of confocal experiments in Fig. S16C showing the AtCESAR2-GFP puncta area size (µm^2^). The area of 218 and 191 puncta for WT and *atg5, respectively,* upon DMSO and a total of 94 and 192 puncta for WT and *atg5*, respectively, upon Bortezomib, were quantified. Two-tailed unpaired Student’s t-tests with Welch’s corrections were performed to analyse the differences of puncta sizes between wild*-*type (WT) and *atg5* backgrounds upon Bortezomib and DMSO treatments. ****, *P* value < 0.0001, *, *P* value < 0.05 (0.0228).

**Figure S17.**
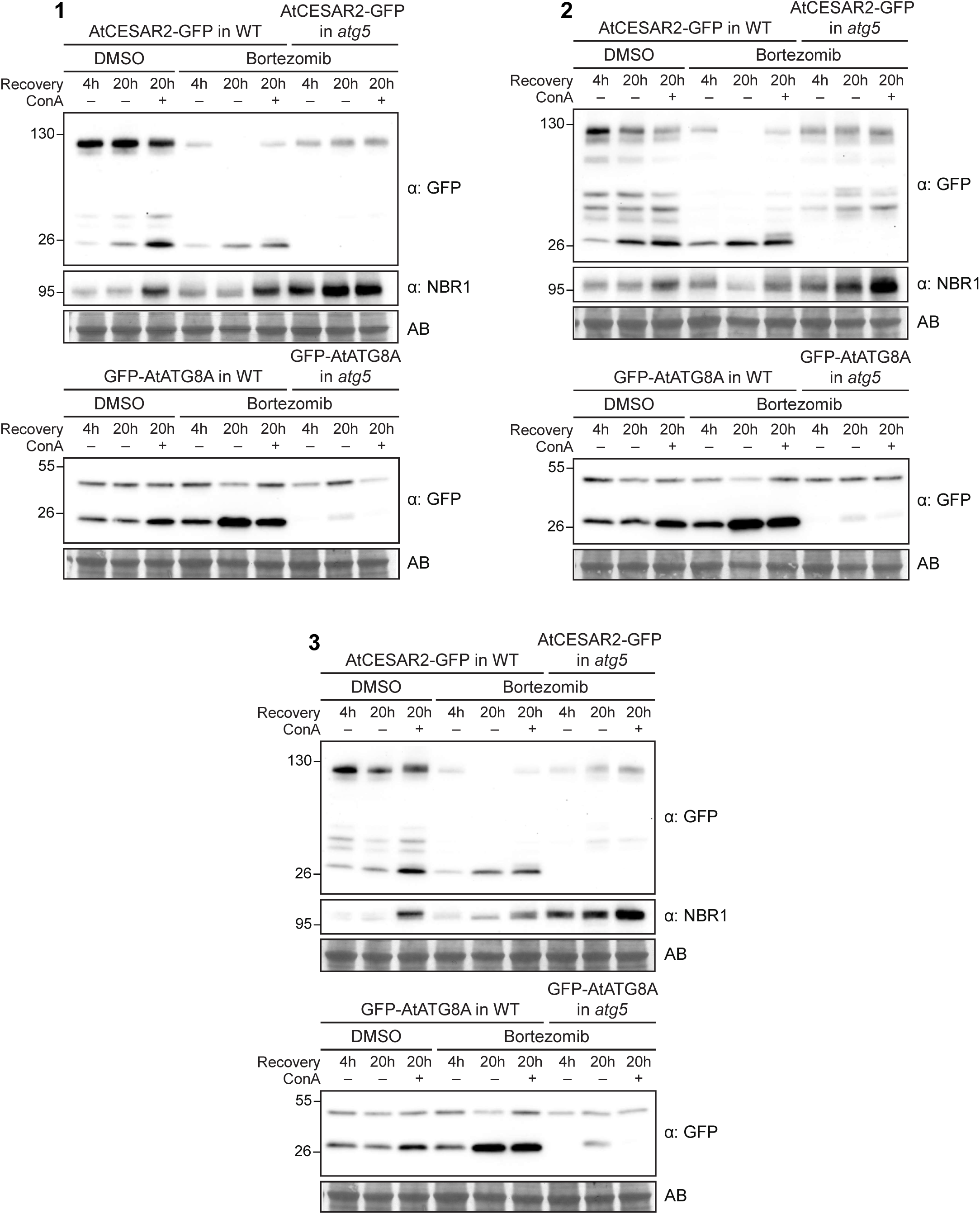
AtCESAR2 undergoes autophagic degradation upon proteasome inhibition. Three independent biological replicates of western blots showing AtCESAR2-GFP (upper panel) and GFP-AtATG8A (lower panel) autophagic flux assays during recovery from proteasome inhibition. *A*. *thaliana* seedlings were grown in ½ MS + MES + 1% sucrose media for 7 days and incubated in media supplemented with DMSO or 50 µM Bortezomib for 4h, followed either by 4h or 20h recovery in fresh media ± 1 μM Concanamycin A (ConA). 20 μg of total protein extract were loaded and immunoblotted with anti-GFP and anti-NBR1 antibodies. Reference protein sizes are indicated on the left side of the blots (unit: kDa). Total protein loading control was analysed by Amidoblack (AB) staining.

**Figure S18.**
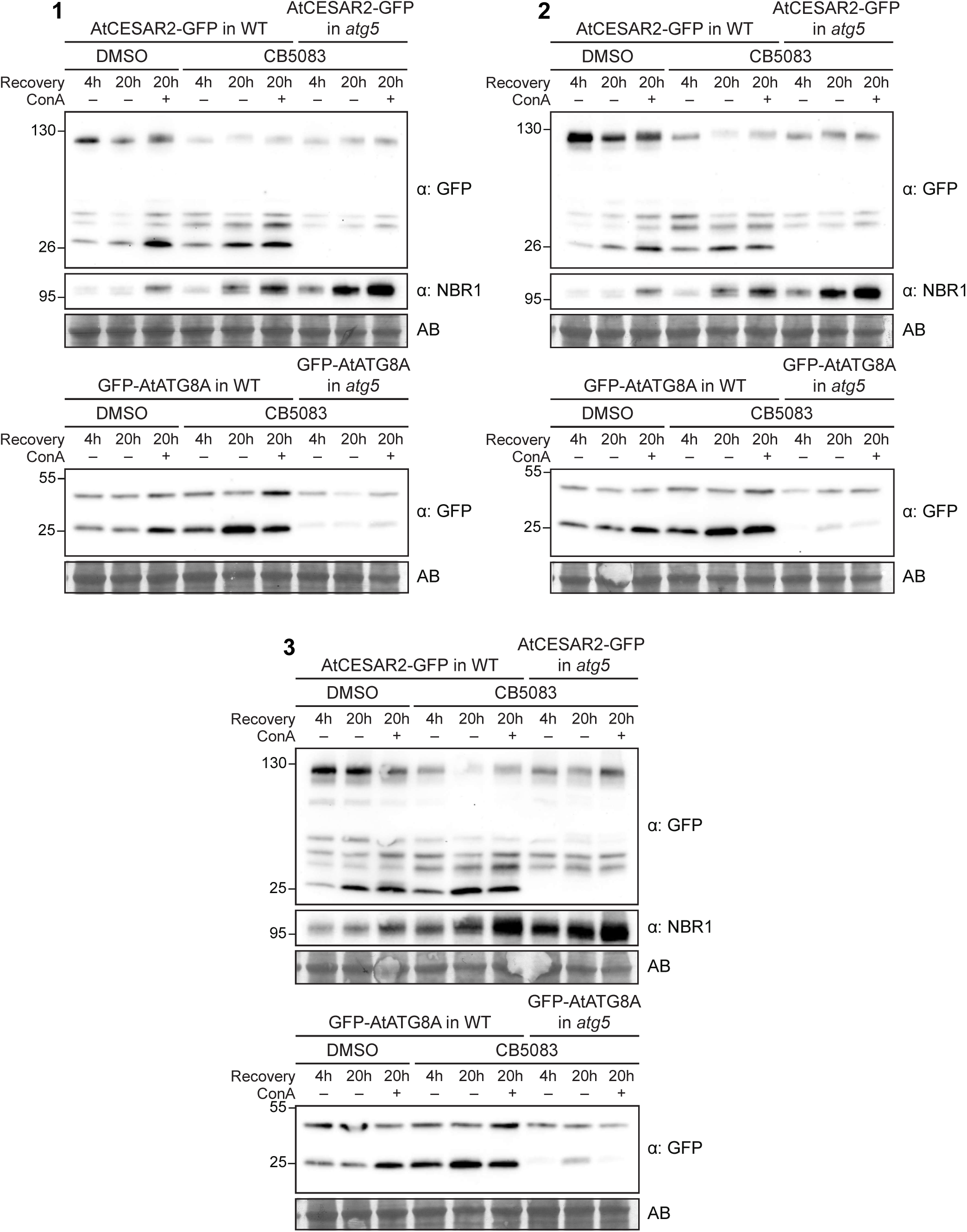
AtCESAR2 undergoes autophagic degradation upon CDC48 inhibition. Three independent biological replicates of western blots showing AtCESAR2-GFP (upper panel) and GFP-AtATG8A (lower panel) autophagic flux assays upon CB5083 treatment. *A. thaliana* seedlings were grown in ½ MS + MES + 1% sucrose media 7 days and incubated in media supplemented with DMSO or 50 µM CB5083 for 4h, followed either by either a 4h or a 20h recovery in fresh media ± 1 μM Concanamycin A (ConA). 20 μg of total protein extract were loaded and immunoblotted with anti-GFP and anti-NBR1 antibodies. Reference protein sizes are indicated on the left side of the blots (unit: kDa). Total protein loading control was analysed by Amidoblack (AB) staining.

**Figure S19.**
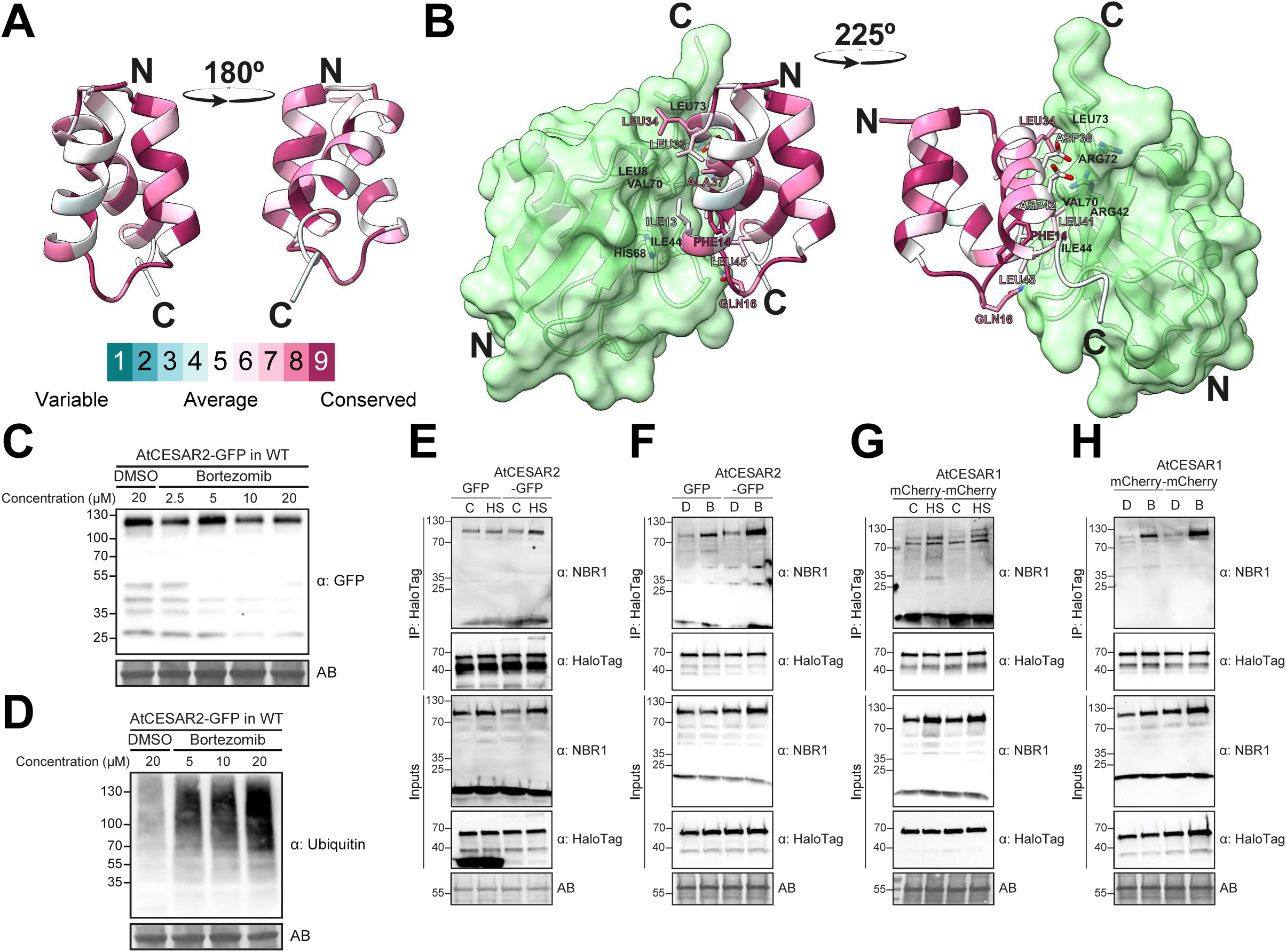
CESAR binds ubiquitin. **(A) CESAR has a conserved CUE domain.** AF2 predicted structure of MpCESAR CUE domain. The conservation scale of aminoacid residues in the CUE domain of all CESAR orthologs is mapped on the structure, N- and C-termini are shown. **(B) CESAR CUE domain binds ubiquitin.** AF2 predicted structure of MpCESAR CUE domain interaction with ubiquitin. Residues involved in the interaction are highlighted in black (ubiquitin) or coloured as shown in Fig. S19A (CESAR CUE), N- and C-termini are shown. **(C) AtCESAR2 protein levels upon proteasome inhibition change in a concentration-dependent manner.** Western blot showing AtCESAR2-GFP protein levels upon incubation with different Bortezomib concentrations. *A. thaliana* seedlings were grown in ½ MS + MES + 1% sucrose media for 7 days and incubated in media supplemented with either DMSO or the indicated Bortezomib concentrations at 21°C for 1h followed by a recovery of 1h in fresh media. 20 μg of total protein extract was loaded and immunoblotted with anti-GFP antibody. Reference protein sizes are indicated on the left side of the blots (unit: kDa). Total protein loading control was analysed by Amidoblack (AB) staining. **(D) Ubiquitinated species accumulate upon proteasome inhibition.** Western blot showing ubiquitin levels upon incubation with different Bortezomib concentrations. *A. thaliana* seedlings were grown under continuous light in ½ MS + MES +1% sucrose media for 7 days and incubated in media supplemented with either DMSO or the indicated Bortezomib concentrations at 21°C for 1h followed by a recovery of 1h in fresh media. 20 μg of total protein extract was loaded and immunoblotted with anti-ubiquitin antibody. Reference protein sizes are indicated on the left side of the blots (unit: kDa). Total protein loading control was analysed by Amidoblack (AB) staining. **(E-H) NBR1 associates with TUBEs.** Halo-tagged TUBE bait and NBR1 co-immunoprecipitation are shown as a positive control for Fig. 4F (Fig. S19E), Fig. 4G (Fig. S19F), Fig. 4H (Fig. S19G) and Fig. 4I (Fig. S19H). 5-d-old *A. thaliana* seedlings expressing either GFP-EV (GFP) or AtCESAR2-GFP (panels E,F), either mCherry-EV (mCherry) or AtCESAR1-mCherry (panels G,H) in wild*-type* (Col-0) background were incubated in liquid ½ MS medium with 1% sucrose for 4 hours at 21°C (Control, C) or 37°C (Heat stress, H) followed by a 4 hour recovery phase at 21°C (panels E,G) or for 1 hour at 21°C in DMSO (D)-supplemented or 5 µM Bortezomib (B)-supplemented media followed by a 1h recovery phase at 21°C in fresh media (panels F,H) and used for co-immunoprecipitation. Plant lysates were incubated with Magne® HaloTag ® Beads conjugated with HaloTag-TUBE. Input and bound proteins were detected by immunoblotting using the respective antibodies as indicated. Reference protein sizes are indicated on the left side of the blots (unit: kDa). Total protein loading control was analysed by Amidoblack (AB) staining.

**Figure S20.**
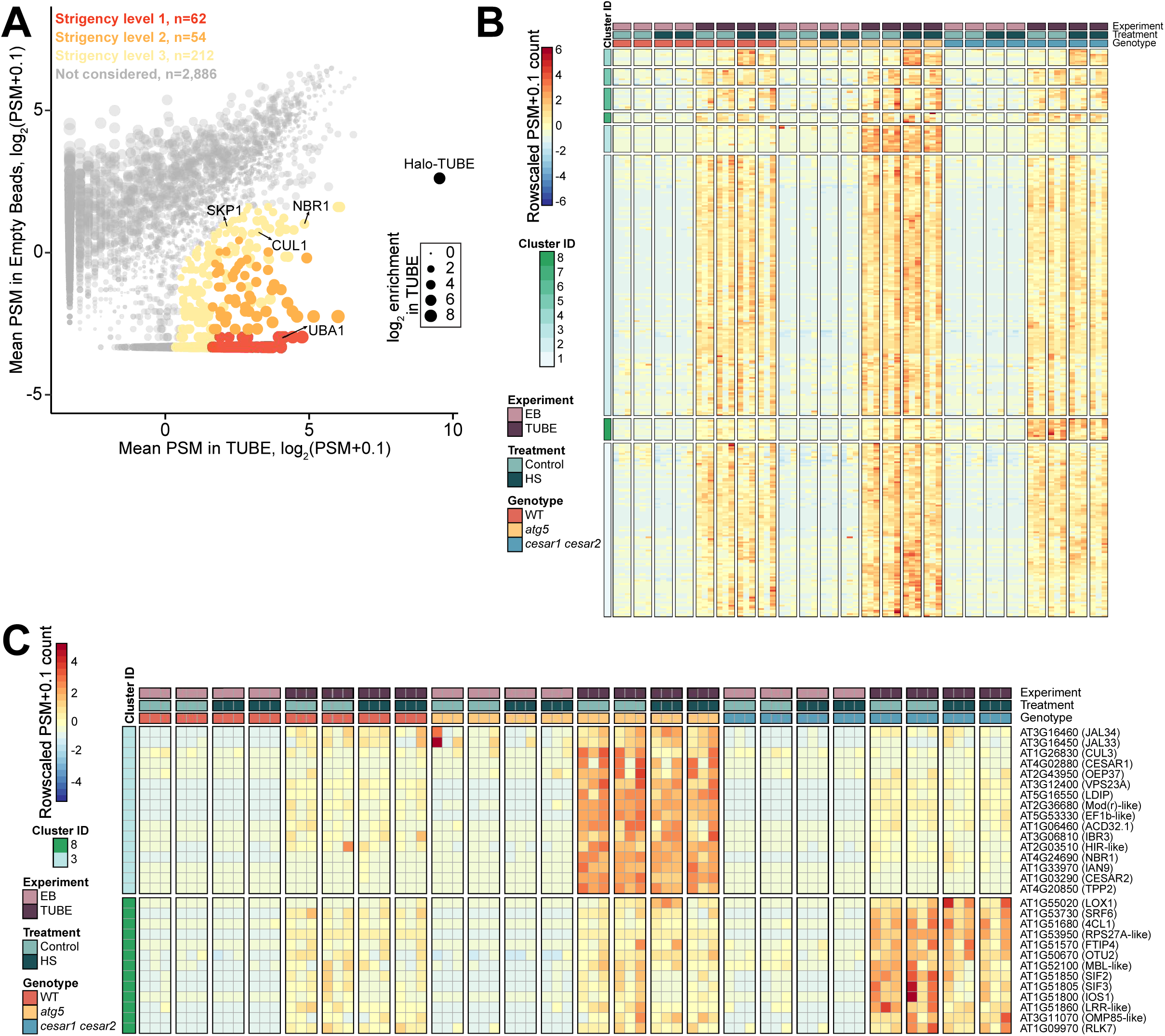
TUBE AP-MS identifies CESAR-regulated proteins upon heat stress recovery. **(A) TUBE AP-MS enriches for ubiquitinated and ubiquitin-associated proteins.** Scatter plot showing combined protein abundance as Log2(PSM+0.1) for the mean PSM value in TUBE samples (x-axis) compared to empty beads (EB) negative control (y-axis) in all genotypes irrespective of treatment. For all stringency levels, a minimum 2-fold enrichment in TUBE samples compared to EB was applied. Stringency levels for consideration of TUBE interactors were defined as follows: a minimum of 3 Peptide-Spectrum Match (PSM) mean value in TUBE samples and less than 3 PSM total value in EB samples (Level 1, red); a minimum of 3 PSM mean value in TUBE samples and a PSM value lower than 3 in any EB samples (Level 2, orange) or a total PSM value greater than the median PSM value in TUBE samples and a mean PSM value lower than 3 in EB samples (Level 3, khaki). Bait and known ubiquitin-binding proteins are highlighted. (B-C) A subset of TUBE interactors is specifically enriched in either *atg5* or *cesar1 cesar2*. Protein abundance pattern represented by a heatmap as (Log2 (Mean PSM+0.1) – mean PSM value per protein) for the 328 proteins classified in any of the stringency levels defined previously and for the 16 and 13 proteins belonging to clusters 8 and 3 in Fig. S20B, respectively. Rows were clustered using Euclidean distance and resulting dendrograms are omitted from the figures.

**Figure S21.**
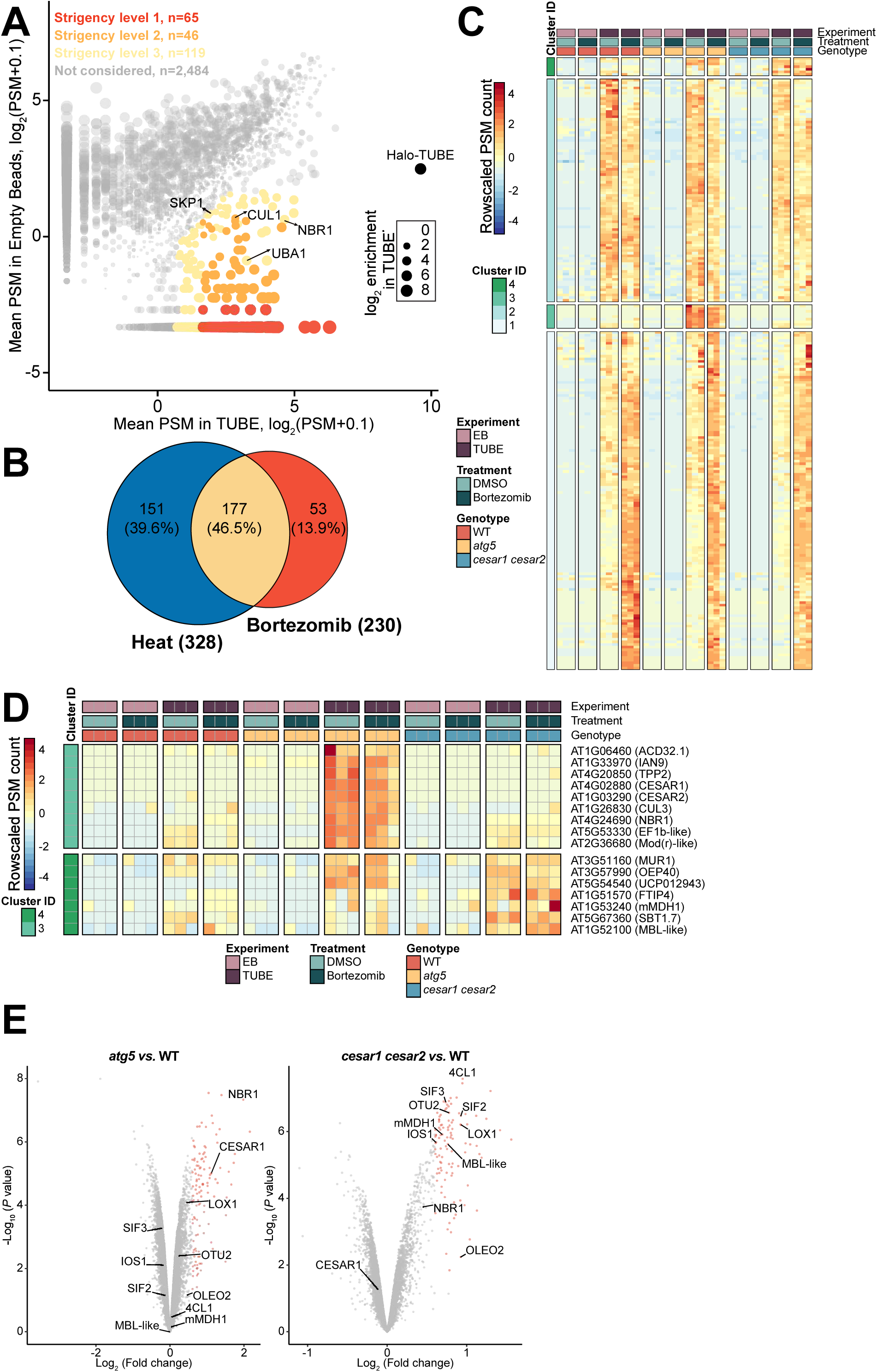
TUBE AP-MS identifies CESAR-regulated proteins upon proteasome inhibition. **(A) TUBE AP-MS enriches for ubiquitinated and ubiquitin-associated proteins.** Scatter plot showing combined protein abundance as Log2(PSM+0.1) for the mean PSM value in TUBE samples (x-axis) compared to empty beads (EB) negative control (y-axis) in all genotypes irrespective of treatment. For all stringency levels, a minimum 2-fold enrichment in TUBE samples compared to EB was applied. Stringency levels for consideration of TUBE interactors were defined as follows: a minimum of 3 Peptide-Spectrum Match (PSM) mean value in TUBE samples and less than 3 PSM total value in EB samples (Level 1, red); a minimum of 3 PSM mean value in TUBE samples and a PSM value lower than 3 in any EB samples (Level 2, orange) or a total PSM value greater than the median PSM value in TUBE samples and a mean PSM value lower than 3 in EB samples (Level 3, khaki). Bait and well-known ubiquitin-binding proteins are highlighted. **(B) TUBE interactome changes upon different proteotoxic stresses.** Venn diagram comparing TUBE interactome upon heat stress (blue circle, see Fig. S20A) or proteasome inhibition (red circle, see Fig. S21A). (C-D) A subset of TUBE interactors is specifically enriched in either *atg5* or *cesar1cesar2*. Protein abundance pattern represented by a heatmap as (Log2 (Mean PSM+0.1) – mean PSM value per protein) for the 230 proteins classified in any of the stringency levels defined previously and for the 9 and 7 proteins belonging to clusters 4 and 3 in Fig. S21C, respectively. Rows were clustered using Manhattan distance and resulting dendrograms are omitted from the figures. (E) Quantitative proteomics of *A. thaliana* reveals autophagy and CESAR-dependent degradomes. Protein enrichment in *atg5* (left) or *cesar1cesar2* (right) compared to wild*-*type (Col-0) represented by a volcano plot. Protein abundance was measured using 16-plex TMT labelling. Enriched proteins are highlighted in red.

**Figure S22.**
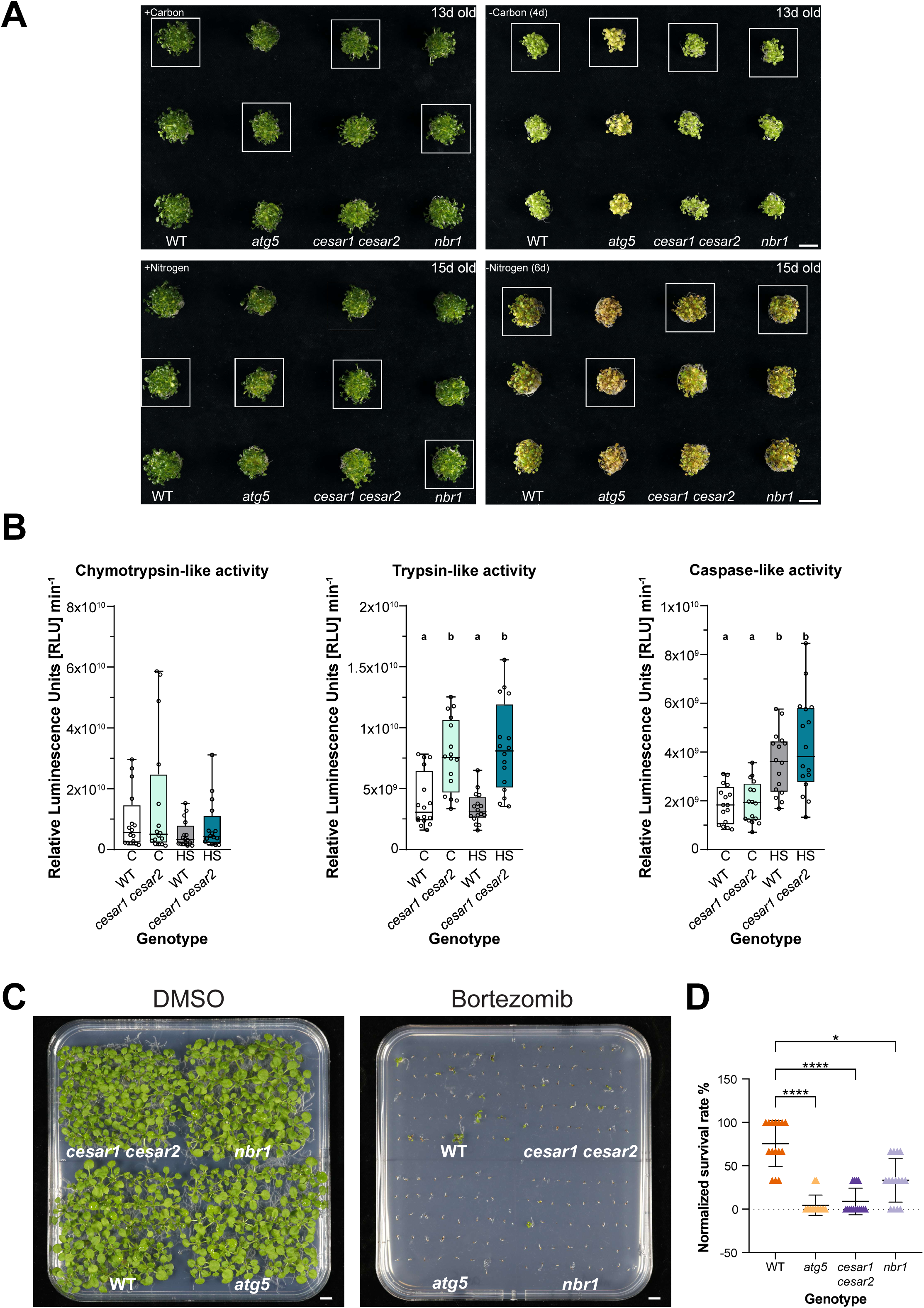
CESAR is necessary for proteotoxic stress tolerance. **(A) *cesar1 cesar2* plants are not hypersensitive to starvation.** Three independent biological replicates starvation assays. 9-d-old *A. thaliana* seedlings of the indicated genotypes were grown in ½ MS + MES + 1% sucrose for 9 days, followed by either 4 days of carbon starvation (-C, left) or 6 days of nitrogen starvation (-N, right), respectively. Representative images in Fig. 5A are highlighted by a white-boxed area. Scale bar, 1 cm. **(B) Proteasome activity is altered in *cesar1 cesar2* plants.** Boxplots representing three different protease activities of the 26S proteasome. Protease activities were measured in wild*-*type (Col-0) or *cesar1 cesar2* mutant *A. thaliana* seedlings grown at 21°C (Control) or 37°C (Heat stress, HS) for 4h, followed by 20h recovery. Protease activities are indicated in Relative Luminescence Units [RLU] min^-1^. Each boxplots represents 16 replicates from four experimental repetitions (n = 4 for each repetition, 6 seedlings per repetition). Statistical significance was determined by one-way ANOVA (*P* value<0.05). **(C) *cesar1 cesar2* plants are hypersensitive to proteasome inhibition.** A second independent biological replicate of bortezomib plates assays shown in Fig. 5D. *A. thaliana* seedlings of the indicated genotypes were grown in 1% agar ½ MS + MES + 1% sucrose plates with either DMSO (left panel) or 3.75 µM Bortezomib for 18 days before imaging. Representative images of 6 independent biological replicates are shown (n = 240). Scale bar, 1 cm. **(D) Quantification of the survival rate of seedlings grown in bortezomib-containing plates.** Normalized survival rate of the biological replicate of Bortezomib plate assays shown in Fig. 5D. Survival rate of each biological replicate was calculated by dividing the number of seedlings that showed a phenotype of size bigger than 0.3 cm and green colour by the total number of seeds sown per genotype (40) and normalized to the highest survival value for the wild*-*type (Col-0) background of all the 15 biological replicates (n = 600), each triangle represents a replicate with 40 seeds per genotype per plate). Kruskal-Wallis test followed by a Dunn’s multiple comparisons test was performed to assess the differences between the survival rates of the different phenotypes. ***, Adjusted *P* value < 0.001; *, Adjusted *P* value < 0.05. Non-significant differences are not shown.

